# The External Globus Pallidus is a Basal Ganglia Output Hub with Action-Specific Circuits

**DOI:** 10.64898/2026.04.12.718022

**Authors:** Zirong Gu, Zachary R. Lewis, Jonathan Tang, Alana Mendelsohn, Ana M. Vicente, Laudan Nikoobakht, Sofia Rosenberg, Ananya Chakravarth, Vitor Paixao, Mihir S. Tirumalasetti, Ghada A. Abuagala, Amir Khalili, Emma D. Thomas, Katherine A. Fancher, Matt Mallory, Arena Manning, Darren Bertagnolli, Jeff Goldy, Christine Rimorin, Michael Tieu, Trangthanh Cardenas, Amy Torkelson, Anish Bhaswanth Chakka, Shenqin Yao, Staci A. Sorensen, Kimberly A. Smith, Luke A. Hammond, Darcy S. Peterka, Hongkui Zeng, Bosiljka Tasic, Rui M. Costa

**Affiliations:** Zuckerman Mind Brain Behavior Institute, Columbia University, New York, NY, USA; The University of Texas at Dallas, School of Behavioral and Brain Sciences, Department of Neuroscience, Richardson, TX, USA; Allen Institute, Seattle, WA, USA; Norcliffe Foundation Center for Integrative Brain Research, Seattle Children’s Hospital, Seattle WA, USA; University of Washington School of Medicine Pediatrics, Seattle, WA, USA; Sania Therapeutics, London, UK; Champalimaud Neuroscience Programme, Champalimaud Research, Champalimaud Foundation, Lisbon, Portugal; Kinetikos, Coimbra, Portugal; Department of Neurology, The Ohio State University, Columbus, OH, USA

**Author notes:** Correspondence to: Zirong Gu; Rui M. Costa. These authors contributed equally to this work (Zirong Gu, Zachary R. Lewis).

## Abstract

The external globus pallidus (GPe) is traditionally viewed as a homogeneous relay in the indirect basal ganglia pathway that broadly suppresses movement. Using whole-brain anterograde axon mapping, rabies tracing, single-neuron reconstruction, and single-nucleus RNA sequencing, we reveal that the GPe is a major basal ganglia output nucleus, composed of anatomically and molecularly distinct populations. Besides the neurons with canonical projections, and projections to striatum and cortex, the GPe contains specific neuronal populations that target the thalamus and brainstem directly, including the parafascicular thalamus (GPe^Pf^) and the pedunculopontine nucleus (GPe^PPN^). These projection-defined subpopulations exhibit distinct behavioral functions. GPe-PPN neurons are selectively suppressed at locomotor onset, and bidirectional manipulation of this pathway is sufficient to promote or suppress locomotion. Notably, D2-MSN stimulation upstream of these neurons evokes locomotion, and co-activation of GPe^PPN^ pathway blocks this D2-MSN-evoked locomotion, demonstrating a disinhibitory circuit for action stemming from D2-MSNs. In contrast, GPe^Pf^ neurons are not engaged during locomotion but are selectively recruited during skilled forelimb actions, and their activation disrupts forelimb movements without affecting locomotion. Together, these findings establish the GPe as a basal ganglia output hub composed of modules that mediate distinct behaviors. This organization revises canonical models of the indirect pathway by demonstrating that D2-MSNs can also facilitate, rather than only suppress, movement depending on the downstream GPe output channels they engage.

**In brief:** The external globus pallidus (GPe), classically considered a relatively homogeneous relay within the indirect basal ganglia pathway, is revealed here as a major basal ganglia output hub. Distinct GPe subpopulations, defined by transcriptomic identity and projection target, form parallel channels to cortical, thalamic and brainstem structures with action-specific functions. These findings uncover a disinhibitory mechanism through which D2-MSNs can facilitate behavior, revising canonical models of basal ganglia function.

**Highlights:** Brain-wide mapping reveals the GPe as a major basal ganglia output node GPe contains novel cell types defined by transcriptomic identity and projection specificity Projection-specific GPe circuits differentially control locomotion and skilled forelimb actions D2-MSNs can disinhibit downstream targets via specific GPe projections and promote action.

## Introduction

The ability to execute specific movements in particular contexts is essential for survival. The basal ganglia are essential for this process, and disorders of the basal ganglia result in profound alterations of behavior and movement. Basal ganglia output nuclei, like globus pallidus internal segment (GPi), and substantia nigra pars reticulata (SNr), are thought to inhibit thalamocortical and brainstem motor centers through GABAergic output projections, thereby preventing unwanted movements, and disinhibition of these centers facilitates movement^1–4^. According to the canonical model, the striatum modulates these basal ganglia output nuclei via two major pathways arising from distinct populations of striatal medium spiny neurons (MSN)¹. The D1-MSN direct pathway is postulated to facilitate movement by directly inhibiting GPi and SNr and therefore disinhibiting thalamic and brainstem targets, whereas the D2-MSN indirect pathway is proposed to suppress movement by increasing the activity of basal ganglia output nuclei through its indirect polysynaptic projections via the external globus pallidus (GPe). This classical framework assumes that GPe is a relatively homogeneous relay within the indirect pathway^1–3,5^ and that it primarily projects within the basal ganglia, to the subthalamic nucleus (STN), the GPi and SNr.

However, accumulating evidence has challenged these assumptions. First, although some studies have reported that stimulation of D1-MSNs promotes movement and stimulation of D2-MSNs inhibits movement, others have shown that both populations are active during movement and that optogenetic stimulation of D2-MSNs can evoke movement and reinforcement^6–12^. Second, the GPe is increasingly recognized as a heterogeneous structure composed of distinct neuron populations with substantial variability in molecular constitution, intrinsic properties and firing dynamics^13–18^. Finally, accumulating evidence suggests that GPe neurons project not only to downstream basal ganglia nuclei (prototypical) and back to the striatum (arkypallidal) but also to targets outside the basal ganglia^19–22^.

These observations highlight the need for a comprehensive and unbiased characterization of GPe output organization. To address this, we performed unbiased brain-wide mapping of the output projections of GABAergic GPe neurons that receive input from the striatum. This approach revealed a broad and previously underappreciated set of output targets, including numerous regions outside the basal ganglia in the cortex, thalamus, and brainstem. We then combined projection mapping with single-cell and single-nucleus transcriptomic profiling to resolve the molecular diversity of these neurons, identifying seven transcriptomically defined cell types with distinct soma locations and anatomical projection patterns. To confirm these observations, we performed sparse labeling and complete (full-brain) reconstruction of individual GPe neurons. These reconstructions revealed striking diversity in axonal architecture: some neurons exhibited highly focused, target-specific projections, whereas others collateralized broadly across multiple downstream regions. Importantly, the projection patterns of individual neurons matched the transcriptomic and retrogradely traced subpopulations, providing convergent evidence for the existence of distinct output circuits from the GPe.

Building on this framework, we interrogated if different GPe output circuits would have different behavioral functions. We focused on two distinct populations of neurons directly projecting to pedunculopontine nucleus (PPN) of the brainstem and parafascicular nucleus (Pf) of the thalamus, as these regions have been traditionally viewed as being disinhibited by D1 MSNs via their direct projections to GPi and SNr. These two GPe output circuits exhibited behaviorally specific engagement: projections to PPN were preferentially modulated during locomotion, whereas projections to Pf were more strongly engaged during learned forelimb actions. Projection-specific optogenetic gain- and loss-of-function experiments demonstrated that GPe projections to PPN bidirectionally modulate locomotion. Critically, stimulation of D2-MSNs upstream of these neurons robustly evokes locomotion that is blocked by co-activation of the GPe-PPN projections, uncovering a disinhibitory circuit for action directly emanating from D2-MSNs. In contrast, activation of GPe projections to Pf selectively disrupts forelimb movements without altering locomotion, demonstrating the behavioral specificity of these circuits.

Together, these experiments redefine the GPe as a major hub composed of molecularly and anatomically defined distinct cell types that project not only within the basal ganglia but also to regions of the cortex, thalamus and brainstem. These populations specifically and directly modulate motor centers in these regions demonstrating that D2-MSNs can give rise to “direct pathways” that disinhibit motor centers.

## Results

### Brain-wide mapping of GPe projections reveals extensive outputs beyond the basal ganglia

To better understand the function of GPe and how it relates to the canonical model of basal ganglia function, there is a strong need for a comprehensive, unbiased characterization of GPe anatomical projections, in particular for the GPe neurons that receive striatal input. Therefore, to identify and quantify GPe output patterns across the brain, we performed brain-wide projection mapping of GPe neurons. We used an intersectional viral labeling strategy combined with BrainJ^23^, a machine learning/based pipeline for pixel classification and mesoscale axonal mapping. Specifically, we targeted GABAergic GPe neurons receiving input from the dorsolateral striatum (DLS), the sensorimotor region of the striatum, by injecting an anterograde adeno-associated virus (AAV) encoding Flpo recombinase into the left DLS of *vGat*-Cre mice (n=7) (**Figure 1A**). In the same animals, we delivered a Cre- and Flpo-dependent AAV-expressing enhanced green fluorescent protein (*AAV-Flex-FRT-eGFP*) into the left GPe. This strategy selectively labeled GABAergic GPe neurons that receive input from the DLS, resulting in the expression of eGFP exclusively in GABAergic GPe cells that receive inputs from DLS (**Figure 1A**). The GFP signal was enhanced by immunohistochemistry, the serial coronal brain sections were imaged and registered to the Allen Mouse Brain Common Coordinate Framework version 3 (CCFv3)^24^. This approach revealed an average of 2,303 ± 435 (n = 7) GFP-positive somata and their spatial distribution within the GPe was quantified (**Figures 1C** and **1D**). Axonal quantification using BrainJ revealed widespread GPe projections across more than 64 target regions. To systematically define putative targets, we used the projection intensity of the PPN, a functionally validated GPe target, as a reference threshold. Regions with average projection intensity exceeding the PPN value (6.8%) were classified as GPe targets (**Figures 1E, 1F, S1A,** and **S1B; Videos S1 and S2**). Notably, only 29.5% ± 3.4% of labeled GPe axons were found in the striatum and 1.7% ± 0.2% in the pallidum, while the remaining 68.7% ± 3.5% extended to cortex, midbrain, thalamus, hindbrain, hypothalamus, and other regions (**Figures 1F, 1G, S1A,** and **S1B; Video S1**). These include 18.2% ± 1.5% of projections directed to discrete cortical areas and specific cortical laminae, 7.6% ± 0.7% to midbrain structures, and 4.6% ± 1.1% to thalamic nuclei. Among the top 40 brain regions receiving GPe input, 50% are located within the cerebral cortex, followed by thalamic (20%) and midbrain (10%) targets (**Figures 1H–1R**, and **S1**; **Video S1**). This anatomical organization reveals a substantial GPe output architecture that extends far beyond the canonical basal ganglia model, which posits that the GPe primarily projects to other basal ganglia nuclei as part of the indirect pathway. Instead, these projections constitute direct GABAergic output channels that are in principle capable of conveying D2-MSN activity to disinhibit downstream motor and associative centers across the brain. In this respect, the GPe functions as an output nucleus analogous to the GPi and SNr in the D1-MSN pathway.

**Figure 1.**
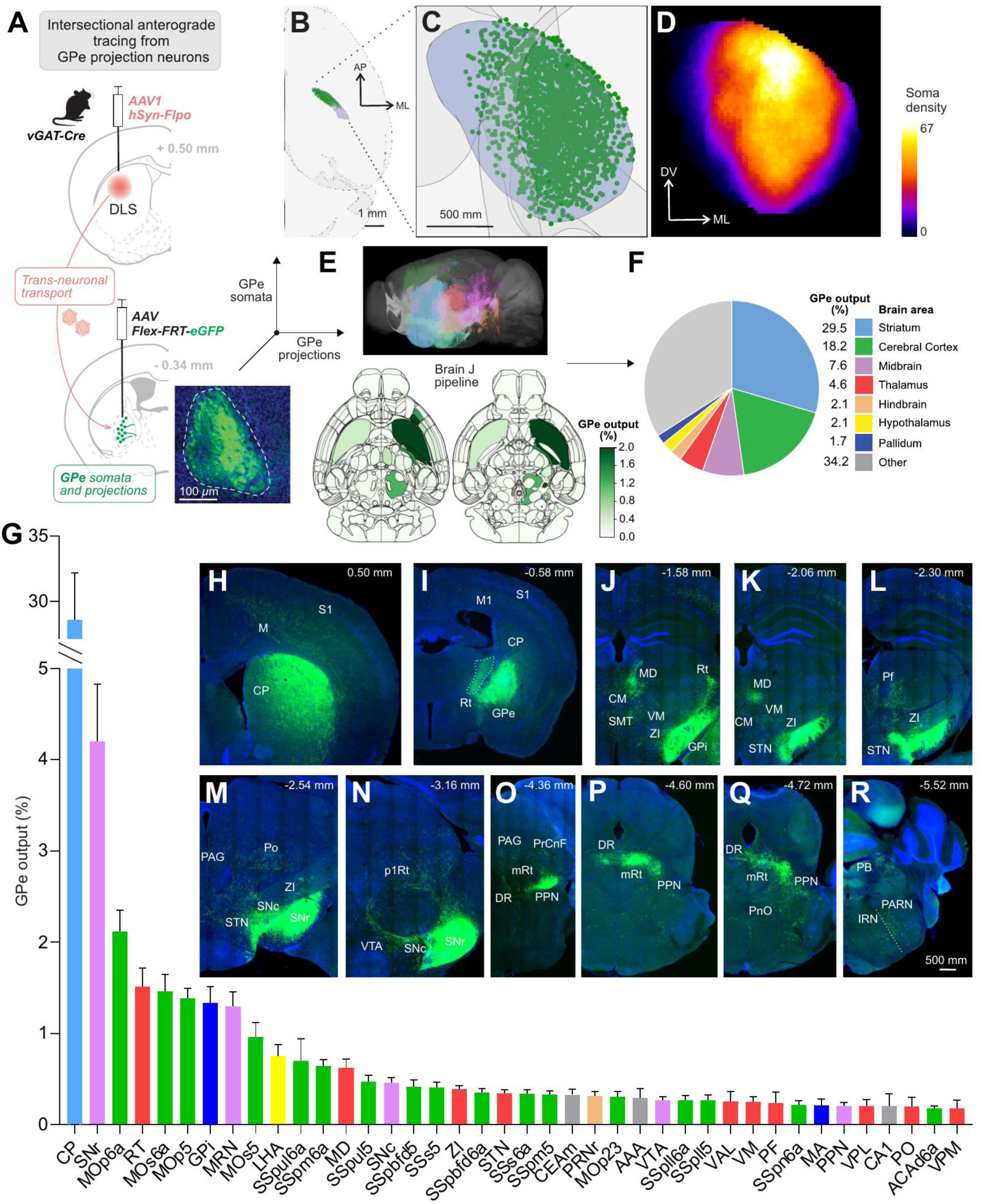
Brain-wide Mapping of GPe Output Targets. **(A)** Schematic of the viral tracing strategy. An *AAV1-hSyn-Flpo* was injected into the dorsolateral striatum (DLS) of *vGAT-Cre* mice (n=7) to express Flpo recombinase in GPe neurons receiving input from MSNs in DLS. A Flpo-dependent reporter virus (*Flex-FRT-GFP*) was injected into the GPe to label GABAergic GPe neurons receiving input from DLS. Inset, representative GFP-labeled GPe neurons labeled by the tracing strategy. (**B–D**) Three-dimensional reconstructions of labeled GPe neurons from a representative brain mapped onto the Allen Mouse Common Coordinate Framework version 3 (CCFv3, 25 µm resolution). (**B**) Top view showing labeled GPe neuron somata. (**C**) Same as (**B**), magnified. (**D**) Soma density heatmap calculated based on (**C**), illustrating the mediolateral and dorsoventral distribution of labeled GPe neurons. (**E**) Top, 3D rendering of axonal projections registered to the Allen CCFv3, revealing extensive GPe outputs to cortical, thalamic, midbrain, and brainstem regions. Bottom, two representative horizontal sections (CCF 0151 and 0180) showing the distribution of GPe axonal projections across the brain. The heat map represents the fractional distribution of total GPe axonal output across brain regions. Color intensity indicates the normalized projection density, corresponding to the fraction of total GPe projection signal detected in each region relative to the sum of all GPe projections across the brain. (**F**) Pie chart showing the relative fraction of total GPe projections across major brain divisions. **(G)** Quantification of the percentage of total GPe projections to the top 40 target regions, expressed as a fraction of the brain-wide GPe output (mean ± SEM, n = 7). **CP**, caudoputamen; **SNr**, Substantia nigra, reticular part; **MOp6a**, Primary motor area, Layer 6a; **RT**, Reticular nucleus of the thalamus; **MOs6a**, Secondary motor area, layer 6a; **MOp5**, Primary motor area, Layer 5; **GPi**, Globus pallidus, internal segment; **MRN**, Midbrain reticular nucleus; **MOs5**, Secondary motor area, layer 5; **LHA**, Lateral hypothalamic area; **SSpul6a**, Primary somatosensory area, upper limb, layer 6a; **SSpm6a**, Primary somatosensory area, mouth, layer 6a; **MD**, Mediodorsal nucleus of thalamus; **SSpul5**, Primary somatosensory area, upper limb, layer 5; **SNc**, Substantia nigra, compact part; **SSpbfd5**, Primary somatosensory area, barrel field, layer 5; **SSs5**, Supplemental somatosensory area, layer 5; **ZI**, Zona incerta; **SSpbfd6a**, Primary somatosensory area, barrel field, layer 6a; **STN**, Subthalamic nucleus; **SSs6a**, Supplemental somatosensory area, layer 6a; **SSpm5**, Primary somatosensory area, mouth, layer 5; **CEAm**, Central amygdalar nucleus, medial part; **PRNr**, Pontine reticular nucleus; **MOp23**, Primary motor area, Layer 23; **AAA**, Anterior amygdalar area; **VTA**, Ventral tegmental area; **SSpll6a**, Primary somatosensory area, lower limb, layer 6a; **SSpll5**, Primary somatosensory area, lower limb, layer 5; **VAL**, Ventral anteriorlateral complex of the thalamus; **VM**, Ventral medial nucleus of the thalamus; **PF**, Parafascicular nucleus; **SSpn6a**, Primary somatosensory area, nose, layer 6a; **MA**, Magnocellular nucleus; **PPN**, Pedunculopontine nucleus; **VPL**, Ventral posterolateral nucleus of the thalamus; **CA1**, Field CA1; **PO**, Posterior complex of the thalamus; **ACAd6a**, Anterior cingulate area, dorsal part, layer 6a; **VPM**, Ventral posteromedial nucleus of the thalamus. **(H–R)** Representative coronal sections showing GFP-labeled axons in major GPe target regions. Scale bars: (**A**) 100 µm; (**C**) 500 µm; (**I**–**S**) 500 µm. See also **Figure S1**, **Video S1**, and **Video S2**.

### Transcriptomic heterogeneity and novel GABAergic cell types in GPe

Previous studies have suggested that the GPe is composed of different cell types based on their transcriptomic profiles^13,25–27^. We therefore combined unbiased single-nucleus RNA sequencing (snRNA-seq) and projection-specific snRNA-seq (Retro-seq) to define the transcriptomic profiles of GABAergic GPe projection neurons^28,29^. We generated an unbiased reference atlas by microdissecting the GPe and neighboring regions from six adult wild-type mice (10x Genomics v3.1, n = 3 mice; 10x Genomics snMultiome, n = 3 mice) (**Figures 2A** and **S2A**). To define and enrich projection-specific subtypes, we bilaterally injected *AAVretro-hSyn-H2B-mCherry* and *AAVretro-hSyn-H2B-eGFP* into two distinct GPe output targets per mouse (**Figures 3A** and **S3A–S3F**). The fluorescent proteins, fused to histone H2B, allowed nuclear localization and provided dual-labeling of projection-defined GPe populations (**Figures 3A**, **S3E**, and **S3F**). In total, 24 mice were injected across three retrograde target-pair groups: Pf (mCherry)/PPN (eGFP), STN (mCherry)/DLS (eGFP), and SNr (mCherry)/MO, SS (eGFP) (n = 8 mice per group). Following virus expression, GPe and adjacent tissue were microdissected, fluorescently labeled nuclei were isolated via fluorescence-activated cell sorting (FACS), and analyzed by snRNA-seq (SMART-Seq v4 (SSv4), Clontech).

**Figure 2.**
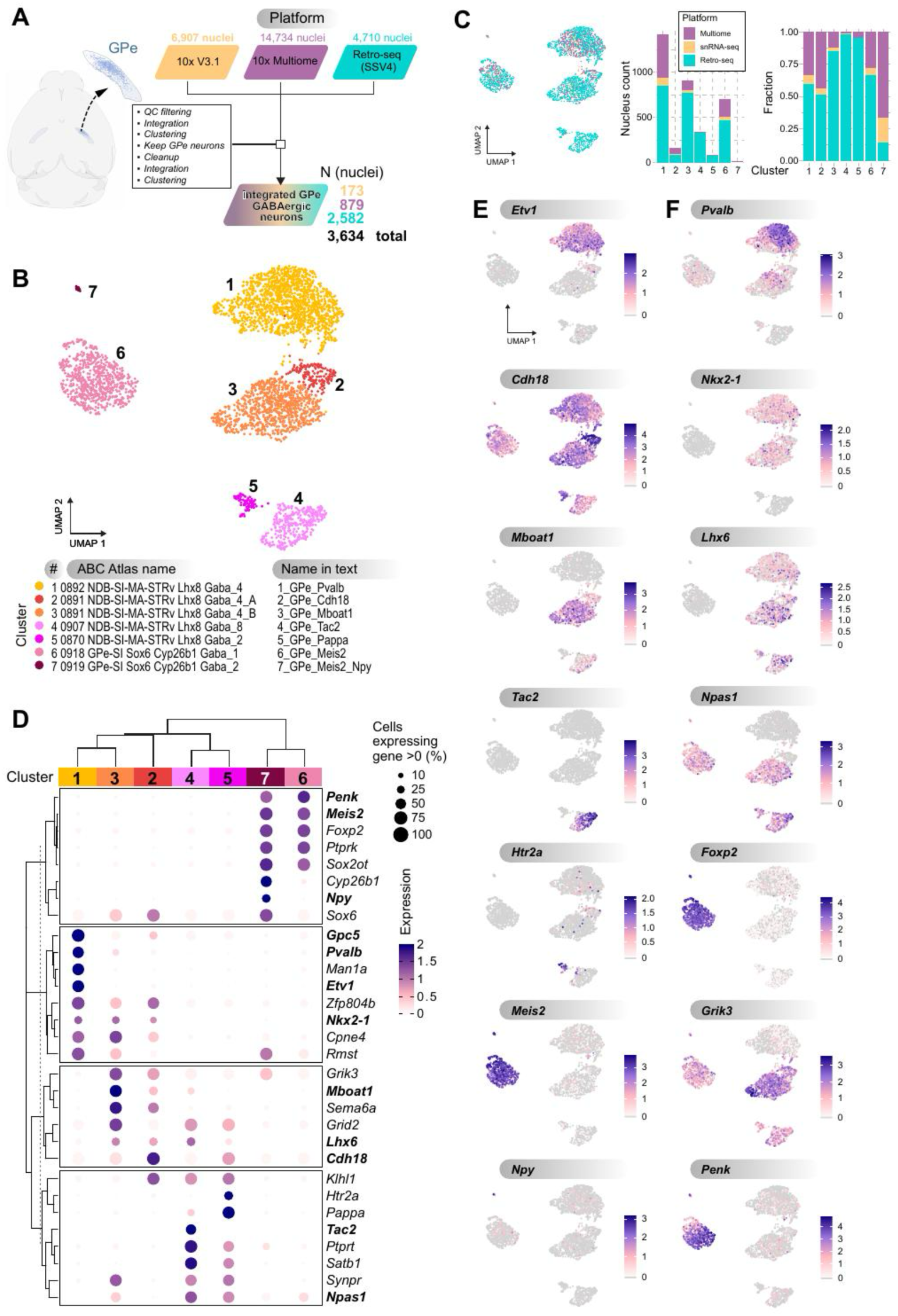
Distinct GPe Neuronal Populations Defined by sn-RNAseq. (**A**) Experimental workflow. Single-nucleus RNA sequencing (snRNA-seq 10x V3.1), single-nucleus multiome sequencing (snRNA-seq and snATAC-seq; 10x Multiome) and single nucleus Retro-seq (SMART-seq V4, SSV4) were integrated and non-GPe cells were removed to define the transcriptomic types of GPe GABAergic neurons. A more complete workflow is available in **Figure S2A**. (**B**) Transcriptomic clustering of GPe neurons. UMAP embedding of 3,634 GABAergic GPe nuclei reveals seven transcriptionally distinct GPe neuronal clusters after exclusion of non-neuronal and cholinergic (ChAT⁺) populations. (**C**) Distribution of sequencing modalities within the final clusters. Left, UMAP embedding of the integrated dataset colored by sequencing platform (Retro-seq SMART-seq v4, 10x snRNA-seq v3.1, and 10x Multiome), illustrating the distribution of nuclei from each platform across the integrated transcriptional landscape. Middle, stacked bar plot showing the absolute nucleus counts contributed by each sequencing modality within each of the seven GPe transcriptomic clusters. Right, stacked bar plot showing the fractional contribution of each sequencing modality within each cluster, demonstrating consistent representation of clusters across datasets following integration. (**D**) Cluster-specific marker genes. Dot plot showing the top differentially expressed genes defining each GPe transcriptomic cluster. Dot size indicates the percentage of cells expressing each gene; color intensity represents mean expression level. Expression is calculated with log_2_((counts/total counts) x 10,000 + 1) scaled to a maximum value of 2.0 for visualization. (**E**) Cluster-specific gene expression patterns. UMAP feature plots showing representative marker genes uniquely enriched in each GPe transcriptomic cluster identified in this study. Distinct clusters are defined by *Etv1*, *Cdh18*, *Mboat1*, *Tac2*, *Htr2a*, *Meis2*, and *Npy* (clusters 1–7, respectively). (**F**) Expression of canonical GPe marker genes. UMAP feature plots showing expression patterns of established GPe markers, including *Pvalb*, *Nkx2-1*, *Lhx6*, *Npas1*, *Foxp2*, *Grik3*, and Penk. Overlapping expression is observed for *Pvalb*, *Nkx2-1*, *Lhx6*, and *Npas1* across multiple clusters.

**Figure 3.**
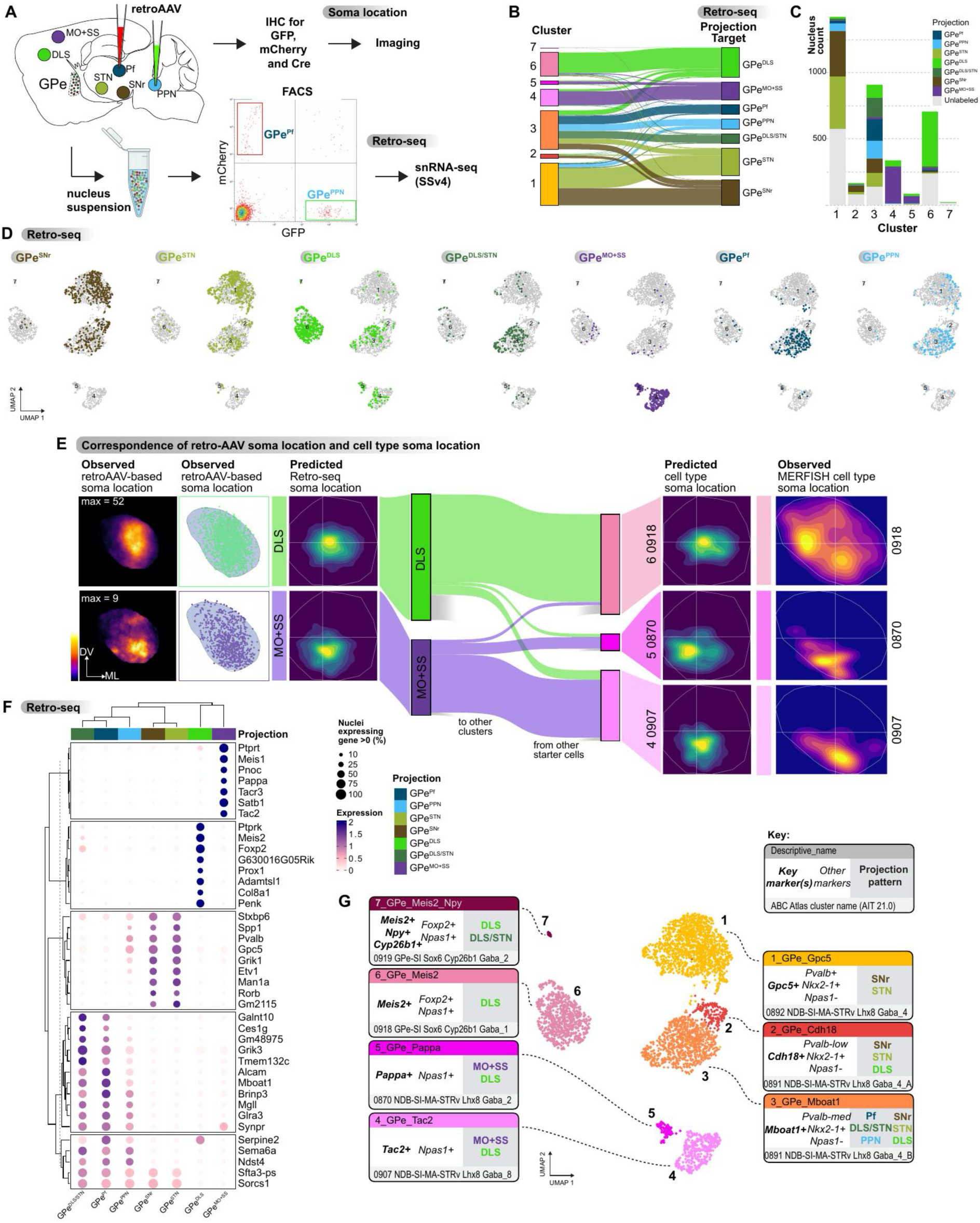
Linking Transcriptomic Identity to Projection-Specific Organization in the GPe. (**A**) Retrograde labeling strategies used to define projection-specific GPe populations and their spatial organization. Two related experimental workflows are illustrated. Top, for soma localization, retrograde AAV vectors expressing nuclear-localized reporters (H2B–mCherry and H2B–eGFP) together with Cre recombinase were injected into three distinct GPe output targets per mouse. Ten mice were used across the following target combinations: Pf (mCherry) / PPN (eGFP) / STN (Cre) (n = 4), DLS (mCherry) / MO–SS (eGFP) / STN (Cre) (n = 4), and MO–SS (mCherry) / PPN (eGFP) / SNr (Cre) (n = 2). After viral expression, brains were processed for immunohistochemistry (IHC) and imaging to determine the spatial distribution of projection-defined GPe neuron somata. Bottom, for projection-specific transcriptomic profiling (Retro-seq), pairs of retrograde AAVs expressing H2B–mCherry and H2B–eGFP were injected into two downstream targets per mouse. Across experiments, 24 mice were distributed across three target-pair groups: Pf (mCherry) / PPN (eGFP), STN (mCherry) / DLS (eGFP), and SNr (mCherry) / MO–SS (eGFP) (n = 8 mice per group). Following viral expression, the GPe and adjacent tissue were microdissected, nuclei were isolated, and fluorescently labeled nuclei were sorted by fluorescence-activated cell sorting (FACS). An example FACS plot from the Pf (mCherry) / PPN (eGFP) group is shown, illustrating separation of projection-defined nuclear populations prior to SMART-seq v4 (Retro-seq) analysis. (**B**) Sankey diagram illustrating the correspondence between transcriptomic clusters (left) and projection-defined cell groups (right). (**C**) Stacked bar plot showing absolute cell counts of each projection-defined GPe population across the seven transcriptomic clusters. (**D**) UMAP visualization of projection-defined GPe neurons (GPe^SNr^, GPe^STN^, GPe^DLS^, GPe^DLS/STN^, GPe^MO-SS^, GPe^Pf^, GPe^PPN^) overlaid onto the integrated GPe transcriptomic space. Distinct projection-defined populations occupy different regions of transcriptomic space, highlighting a structured correspondence between transcriptional identity and projection target across GPe neuron types. (**E**) Transcriptomic identity predicts soma location for projection-defined GPe populations. From left to right, heatmaps show empirically observed soma locations of GPe neurons labeled by retrograde AAV injections targeting the dorsolateral striatum (GPe^DLS^, top row) or motor and somatosensory cortices (GPe^MO-SS^, bottom row), followed by the corresponding experimentally observed soma distributions in the GPe. Next, soma locations predicted from Retro-seq gene expression using MERFISH-trained spatial models reveal distinct and partially overlapping spatial domains for GPe^DLS^ and GPe^MO-SS^ neurons. Sankey-style mappings illustrate the correspondence between projection-defined populations and transcriptomic clusters, highlighting strong alignment of GPe^DLS^ neurons with AIT21 cluster 6_0918 and GPe^MO-SS^neurons with AIT21 clusters 5_0870 and 4_0907. Predicted soma locations based on transcriptomic cluster identity closely recapitulate the spatial distributions observed in MERFISH cell-type maps (right), demonstrating that transcriptomic identity captures the spatial organization of GPe^DLS^ and GPe^MO-SS^ neurons within the GPe. (**F**) Marker genes for projection-defined GPe populations. Dot plot showing the top differentially expressed genes associated with each projection-defined GPe population, revealing four dominant projection patterns: SNr/STN, Pf/PPN, DLS, and MO–SS. Distinct molecular signatures characterize each group: *Gpc5*, *Pvalb*, *Etv1*, and *Grik1* are enriched in GPe^SNr^ and GPe^STN^ neurons; *Mboat1*, *Brinp3*, *Alcam*, and *Synpr* define GPe^Pf^ and GPe^DLS/STN^populations; *Prox1*, *Adamtsl1*, *Foxp2*, and *Meis2* mark GPe^DLS^ neurons; and *Tac2*, *Tacr3*, *Pnoc*, and *Pappa* distinguish GPe^MO-SS^ neurons. Dot size indicates the fraction of cells expressing each gene, and color intensity represents mean expression level. (**G**) Summary of defining transcriptional markers and projection targets for each of the seven GPe clusters, including novel subtypes such as cluster 3_GPe_Mboat1, which exhibits multi-target projections to DLS, STN, SNr, Pf, and PPN.

We integrated the scRNA-seq (14,734 cells), snRNA-seq (6,907 nuclei), and SSv4 Retro-seq (4,710 nuclei) datasets, yielding a total of 26,351 cells/nuclei for downstream analysis (**Figures 2A**, **3A**, **S2A**, and **S2B**). After quality control and exclusion of non-neuronal and cholinergic (ChAT⁺) clusters, we identified seven different GABAergic neuronal clusters in the GPe, more than previously acknowledged, and differentially expressed genes across the clusters (**Figures 2B, 2D,** and **S2B–S2F**). This final GABAergic neuron subset contained 3,634 neurons (173 from 10x Genomics v3.1 snRNA-seq, 879 from 10x Genomics snMultiome, and 2582 from Retro-SMART-seq).

We sought to identify new molecular markers emerging from our dataset that could define individual transcriptomic clusters. Differential expression analysis revealed that *Etv1*, *Cdh18, Mboat1, Tac2, Meis2, Pappa* and *Meis2/Cyp26b1/Npy* are reliable markers for clusters 1 through 7, respectively (**Figures 2D** and **2E**), some of which were previously identified^25–27^. These markers can be coupled with established markers for GPe neural cell types to define a variety of cell subgroupings. We then characterized the expression patterns of canonical GPe markers, including *Pvalb*, *Nkx2.1*, *Lhx6*, *Npas1*, *Foxp2*, *Grik3*, and *Penk* (**Figure 2F**). Although *Pvalb* and *Npas1* have been described as defining mutually exclusive cell types^13,15,17,20,27^, our data revealed extensive overlap between these markers, particularly in cluster 3_GPe_Mboat1 and 6_GPe_Meis2 (**Figures 2F**, **S2F**, and **S2G**). Moreover, *Pvalb* expression varied along a continuum and could be stratified into high, medium, and low levels (**Figures S2F** and **S2G**). We also observed substantial co-expression of *Pvalb* with *Nkx2.1* and *Lhx6* (**Figures S2F** and **S2G**), further challenging the notion that these markers define strictly segregated molecular or functional GPe subtypes^30^. The classically-defined prototypical neurons fall into cluster 1_GPe_Pvalb (*Pvalb+/Nkx2-1+/Npas1-*), while the classically-defined arkypallidal neurons likely belong to clusters 6_GPe_Meis2 and 7_GPe_Meis2_Npy (**Figure 2B**). Together, these transcriptomic results define seven distinct GABAergic GPe cell types, including two clusters corresponding to the classically described prototypical and arkypallidal neurons ^25^. The increased resolution helped reveal further sub-structure in the other clusters beyond what was previously described ^25^. For instance, we subdivide Saunders “subcluster 2-17” into clusters 2_GPe_Cdh18 and 3_GPe_Mboat1 and divide “subcluster 2-18” into clusters 4_GPe_Tac2 and 5_GPe_Pappa. We do not find support for subclusters 2-16, 2-15, or 2-21 (**Figure S2K**), which fell into our integrated clusters 4 and 7 and are outliers characterized by poor QC characteristics (not shown). Conversely, we assign “prototypical” GPe neurons to one cluster (1_GPe_Pvalb), while Saunders et al. assigned it to subclusters 2-13 and 2-14 (**Figure S2L**). Together, these analyses establish a high-resolution transcriptomic atlas of GPe projection neurons, identifying seven GABAergic cell types, linking projection-defined populations to specific molecular clusters, and revealing greater molecular diversity within the GPe than previously appreciated.

### Relation between projection patterns, transcriptomic identity, and soma location in GPe

We next investigated how transcriptomic identity relates to anatomical organization and projection specificity by integrating molecular profiling with soma localization and projection pattern analysis. Retrograde labeling of projection-defined GPe subpopulations followed by whole-brain registration and 3D reconstruction revealed that each subpopulation exhibited a distinct spatial bias in soma localization within the GPe^31^ (**Figures 3A** and **S3**). GPe^DLS^ somata were concentrated in the mid-to-lateral regions; GPe^MO-SS^ somata were biased ventrally; GPe^Pf^ somata were more dorsally distributed; GPe^PPN^ somata localized ventrally; GPe^STN^ somata were centrally positioned; and GPe^SNr^ somata were biased medially (**Figures 3E**, **S4E,** and **S4F**).

To determine how transcriptomic identity relates to anatomical projection, we integrated Retro-seq data from projection-defined GPe neurons with our full GPe transcriptomic dataset. This analysis revealed distinct projection patterns across the different transcriptomic clusters (**Figures 3B**, **3C**, and **3D**). First, we found that cluster 1_GPe_Gpc5 was predominantly composed of GPe neurons projecting to the STN (47%) and SNr (40%), consistent with the identity of prototypical GPe neurons known to target these downstream basal ganglia structures (**Figures 3B**, **3C**, and **3D**). Second, GPe^DLS^ neurons were highly enriched in cluster 6_GPe_Meis2 (87%), supporting the designation of this cluster as the arkypallidal cell type that projects densely back to the striatum (**Figures 3B, 3C,** and **3D**). Third, GPe^MO-SS^ neurons were primarily localized to clusters 4_GPe_Tac2 (82%) and 5_GPe_Pappa (68%) (**Figures 3B**, **3C**, and **3D**), suggesting that these populations represent output channels to the cortex. Finally, cluster 3_GPe_Mboat1 emerged as a unique transcriptional group composed of neurons that broadly project to multiple brain regions, including DLS, STN, SNr, Pf, and PPN (**Figures 3B**, **3C**, and **3D**). Notably, neurons co-labeled from both STN and DLS were predominantly found in cluster 3_GPe_Mboat1 (19%), indicating that this cluster represents a novel GPe cell type distinct from classical arkypallidal neurons (**Figures 3B**, **3C**, and **3D**). Interestingly, cluster 2_GPe_Cdh18 projected to the SNr and STN, resembling prototypical GPe neurons, while also innervating the striatum, a hallmark of arkypallidal populations. This mixed projection pattern suggests that cluster 2 represents a transcriptomically defined population that bridges features of both classical GPe cell types.

We next sought to identify gene expression signatures associated with distinct GPe projection patterns. To this end, we performed differential gene expression analysis across the projection-defined subpopulations and found that GPe neurons exhibited projection-specific transcriptional profiles. A dendrogram was constructed to visualize the relationships among these groups based on shared gene expression patterns, revealing five major expression patterns (**Figure 3F**). GPe^Pf^ neurons were characterized by a unique combination of markers, including *Mboat1*, *Brinp3*, *Alcam*, and *Synpr*. This transcriptional signature was also observed in GPe^DLS/STN^ cells, suggesting a molecular similarity between these populations (**Figure 3F**). GPe^DLS/STN^ cells represent a distinct population identified during FACS that projects to both DLS and STN. Second, GPe^PPN^ neurons shared some features with GPe^Pf^ cells but could be distinguished by higher expression of *Gpc5* and lower expression of *Mboat1*, *Brinp3*, *Alcam*, and *Synpr* (**Figure 3F**). Third, GPe^SNr^and GPe^STN^neurons exhibited highly similar gene expression profiles and were enriched for *Gpc5*, *Stxbp6*, *Grik1*, *Pvalb*, and *Etv1* (**Figure 3F**). Fourth, GPe^DLS^ neurons displayed a distinct transcriptional identity, marked by elevated expressions of *Prox1*, *Adamtsl1*, *G630016G05Rik*, *Col8a1*, *Foxp2*, *Meis2*, *Penk*, and *Ptprk* (**Figure 3F**). Finally, GPe^MO+SS^ neurons were defined by high expression of *Tac2*, *Tacr3*, *Pnoc*, *Ptprt*, *Stab1*, *Meis1*, and *Pappa* (**Figure 3F**). Together, these results reveal projection-enriched gene expression programs across the seven projection-defined GPe subpopulations, providing molecular insight into their distinct anatomical and functional identities.

To investigate the relationship between transcriptomic identity and soma location across projection-defined GPe subpopulations, we trained predictive models using spatial transcriptomic data from the multiplexed error-robust fluorescence in situ hybridization (MERFISH) data^32^ (**Figures 3D, 3E,** and **S4**). The trained MERFISH-based models were then applied to our Retro-seq dataset, using shared gene features to infer the spatial coordinates of GPe neuron somata (**Figure S4A**). Predicted soma locations were then compared with empirically observed distributions derived from MERFISH data^32^ and experimentally observed somata localization following AAVretro injections (**Figures 3E**, **S4E–S4J).** The predictive model revealed that GPe^DLS^ somata are concentrated in the mid-to-lateral regions, corresponding closely to the territory occupied by cluster 6_GPe_Meis2 (AIT21 cluster 0918) predicted from MERFISH spatial data (**Figures 3E** and **S4G–S4J**)^32^. Predicted soma distributions further indicated that GPe^MO-SS^ neurons projecting to the motor and somatosensory cortices are ventrally localized within the GPe, and that transcriptomic clusters 4_GPe_Tac2 and 5_GPe_Pappa exhibit finer spatial segregation into ventrolateral and ventromedial territories, respectively, GPe^Pf^ somata occupy more dorsal positions, GPe^STN^ somata are centrally located, and GPe^SNr^somata are biased medially (**Figures S4G–S4J**). The exception were GPe^PPN^ somata, whose predicted locations were centrally positioned, consistent with MERFISH-based predictions but differed from retrograde tracing results (**Figures S4G–S4J**). Together, these analyses demonstrate that molecular identity, projection target, and soma position are correlated features of GPe neurons.

### Single-cell reconstructions reveal diverse projection patterns of GPe neurons

We next sought to understand if individual GPe projection neurons had the projection patterns observed for the transcriptomic cell types described above. To map the projections of individual neurons in GPe, we performed sparse labeling of GPe neurons by crossing Cre-driver lines including, Npr3-IRES2-CreERT2-neo and Pvalb-T2A-CreERT2, to Ai166 transgenic reporter line, which expresses eGFP sparsely and strongly in a Cre-dependent manner^33^. Alternatively, we stereotaxically injected Penk-IRES2-Cre-neo mice with an AAV expressing eGFP sparsely and strongly in a Cre-dependent manner **(Figure 4A**, see Methods). This strategy enabled sparse but bright labeling of individual GPe neurons. Resin-embedded, GFP-labeled brains underwent chemical reactivation to recover GFP fluorescence and were imaged across the entire brain using fluorescence micro-optical sectioning tomography (fMOST)^34^, generating datasets with submicron lateral resolution (0.35 × 0.35 µm) and 1 µm axial sampling. Whole neuron morphology (WNM) reconstructions were performed using Vaa3D-TeraVR, allowing manual tracing of dendrites and complete local and long-range axonal arbors. Completed reconstructions were post-processed to ensure continuity and eliminate tracing errors. fMOST image volumes were registered to the CCFv3, and reconstructed neurons were transformed into CCF space using the resulting deformation fields^35^. To quantify projection patterns, reconstructed SWC files were resampled and used to generate projection matrices based on axonal node counts within anatomically defined target regions, including both ipsilateral and contralateral structures, with fibers of passage excluded. In total, we reconstructed 20 GPe neurons with somata localized within the GPe (**Figures 4B–4F**; **Video S3**). Confirming what was observed in Figure 1, these single neurons exhibited surprisingly vast and diverse projection patterns, with distinct subsets preferentially targeting different brain regions, including specific cortical areas and laminae, as well as subcortical structures such as the hypothalamus, hippocampus, striatum, pallidum, thalamus, and midbrain (**Figures 4B–4F** **and S5C**; **Video S3**). We performed K-means clustering and established 4 different groups of GPe neurons based on their single-cell projectomes (**Figures 4C–4E**, **S5A, and S5B**). For example, neuron 6 from group 1 is a representative example of transcriptomic type 1_GPe_Gpc5, sending a dense axonal projection to the STN and SNr with very few detectable axon collaterals in other brain regions. Neuron 7 from group 1 is a representative example of 2_GPe_Cdh18, projecting to both SNr and STN, while also innervating the striatum. Neuron 18 from group 2 exemplifies 3_GPe_Mboat1 characteristics, extending axons to cortex, striatum, thalamus, and brainstem, highlighting its multi-projection architecture. Neuron 14 from group 3 represents 4_GPe_Tac2 and 5_GPe_Pappa, projecting robustly to the motor and somatosensory cortices (MO + SS). Neuron 27 from group 4 represents 6_GPe_Meis2 and 7_GPe_Meis2_Npy, exhibiting a focused projection to the striatum with no collaterals to other regions. Together, these single-neuron reconstructions illustrate that GPe neurons exhibit a spectrum of axonal projection patterns, from highly target-specific, one-to-one projections to broader, multi-target arborizations. This diversity aligns closely with molecularly and retrogradely defined subpopulations, supporting a model in which distinct GPe transcriptomic clusters form parallel and minimally overlapping output channels. Transcriptomic cluster 3_GPe_Mboat1 includes neurons with broad projection patterns. However, we did not observe neurons projecting simultaneously to Pf in the thalamus and PPN in the midbrain, two prominent downstream targets of canonical basal ganglia output mediated by GPi and SNr. Therefore, we examined specifically if GPe^Pf^ and GPe^PPN^ neurons exhibit distinct or overlapping patterns of axonal collateralization. To do this, we employed a cell type- and projection-specific monosynaptic rabies tracing strategy, enabling precise visualization of the axon collateral profiles of each population. We first injected *AAV1.hSyn.DIO.TVA(66T).3XHA.2A.N2C(G)* into the Pf (n=6 mice, **Figures S6A–S6D**) or PPN (n=5 mice, **Figures S6M–S6P**) of *vGlut2-Cre* mice to drive Cre-dependent expression of the avian TVA(66T) receptor and rabies glycoprotein N2C(G), which are required for selective infection and transsynaptic labeling using the EnvA-pseudotyped rabies virus^36^ (**Figures 4L** and **4R**). In the same animals, we injected *AAV5-EF1a-fDIO-EYFP* into the GPe to enable Flpo-dependent reporter expression. Two weeks later, we delivered EnvA-pseudotyped, G-deleted CVS-N2c rabies virus encoding Flpo recombinase fused to mCherry (*EnvA-N2cΔG-FlpO.mCherry*) into either Pf or PPN^37^. This approach retrogradely labels GPe neurons that provide monosynaptic input to Pf or PPN and enables Flpo-mediated EYFP expression exclusively in those presynaptic GPe neurons (**Figures 4G–4J**). Immunohistochemical enhancement of EYFP followed by serial coronal section imaging and BrainJ-based analysis revealed sparse but specific labeling of GPe neurons (**Figures 4K** and **4Q**). Registration to the CCFv3 and 3D reconstruction showed that GPe^Pf^ and GPe^PPN^ neurons were spatially segregated (**Figures 4G–4J**). GPe^Pf^ neurons were predominantly located in the dorsal GPe, whereas GPe^PPN^ neurons were biased ventrally (**Figures 4G–4J**), consistent with the soma distributions observed in our AAVretro tracing experiments (**Figures S4E** and **S4F**). We next analyzed the axonal collateralization of these projection-defined populations. GPe^Pf^ neurons sent dense axonal projections to the Pf with minimal or no collaterals to PPN, DLS, M1, GPi, STN, or SNr (**Figures 4M–4P** and **S6A–S6L**). Conversely, GPe^PPN^ neurons projected densely to the PPN but exhibited little to no collateralization to Pf or other GPe output targets (**Figures 4S–4V** and

**Figure 4.**
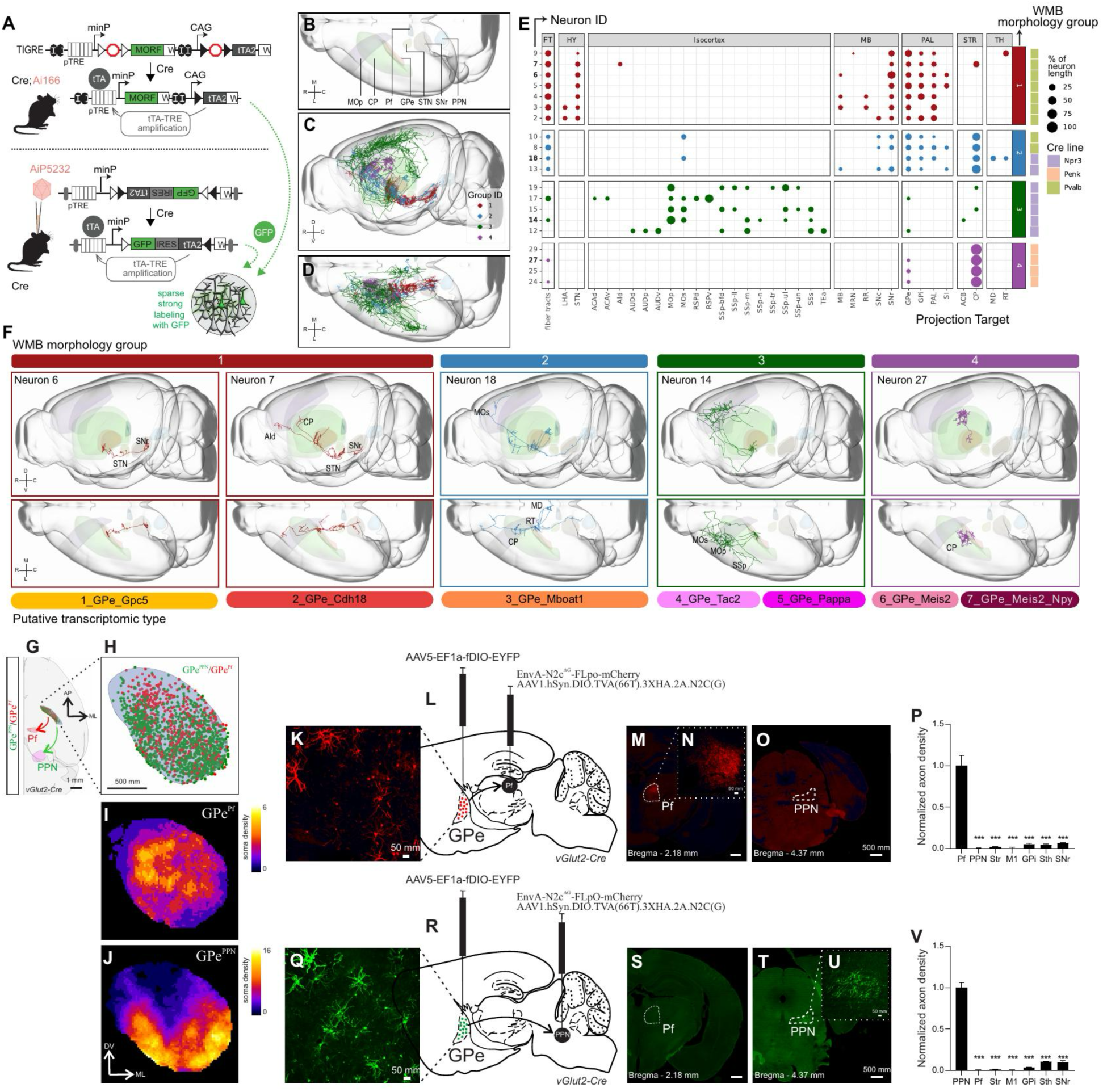
Cellular and Subpopulation Architecture of GPe Output Collateralization. (**A**) Strategy for sparse and strong labeling of individual GPe neurons for complete single-cell reconstruction. Top, schematic illustrating the all-transgenic strategy: Cre-driver mouse lines (Npr3-IRES2-CreERT2-neo or Pvalb-T2A-CreERT2) were crossed with the Ai166 reporter line. Bottom, a complementary transgenic + viral strategy: Penk-IRES2-Cre-neo mice were injected with a Cre-dependent AAV expressing GFP (AiP5232). (**B**) Schematic top view of the mouse brain illustrating the external globus pallidus (GPe) and major downstream target regions of GPe projections, including primary motor cortex (MOp), dorsal striatum (STRd), parafascicular thalamus (Pf), subthalamic nucleus (STN), substantia nigra pars reticulata (SNr), and pedunculopontine nucleus (PPN). (**C** and **D**) 3D-renderings of 20 single-neuron reconstructed GPe neurons illustrating the diversity of axonal trajectories and projection targets across cortical, thalamic, and brainstem regions. Reconstructions are shown from a dorsal view (**C**) and from lateral and dorsal views (**D**). Neurons are colored according to their projection group. (**E**) Hierarchical clustering of single-cell axon projection maps showing groups of neurons with similar collateralization patterns. (**F**) Representative single-neuron reconstructions illustrating correspondence between individual GPe projectomes and transcriptomic-defined clusters. Neuron 6 (cluster 1_GPe_Gpc5; Pvalb/prototypical) sends dense projections to STN and SNr with minimal collateralization to other regions. Neuron 7 (cluster 2_GPe_Cdh18) projects to STN and SNr and additionally innervates the striatum, demonstrating mixed projection features. Neuron 18 (cluster 3_GPe_Mboat1) exhibits broad, multi-target projections to cortex, striatum, thalamus, and brainstem. Neuron 14 corresponds most closely to clusters 4_GPe_Tac2 and 5_GPe_Pappa and projects prominently to motor and somatosensory cortex. Neuron 27 represents arkypallidal clusters 6_GPe_Meis2 and 7_GPe_Meis2_Npy and shows a focused projection to the striatum with minimal collateralization elsewhere. Each neuron is color-coded and grouped according to its collateralization pattern. (**G–J**) Soma localization and 3D reconstructions of GPe neurons labeled by monosynaptic rabies tracing projecting to Pf and PPN in *vGluT2-Cre* mice. (**G**) Example horizontal view showing segregated Pf- and PPN-projecting GPe neurons. (**H**) Coronal view of 3D distribution of Pf-projecting (red) and PPN-projecting (green) neurons mapped to the Allen Brain Atlas CCFv3. Soma density heatmaps showing dorsal enrichment of GPe^Pf^neurons (**I**) and ventral bias of GPe^PPN^neurons (**J**). (**K–P**) Projection-specific monosynaptic rabies identification of GPe^Pf^ neurons. (**K**) High-magnification image of labeled GPe^Pf^ neurons. (**L**) Experimental schematic showing viral injection and monosynaptic rabies tracing strategy to identify GPe^Pf^ neurons in *vGlut2-Cre* mice. **(M–O)** Representative coronal sections showing dense GPe^Pf^ axon terminals in Pf (**M**, **N**) and absence of collaterals in PPN (**O**). Inset (**N**), magnified view of dense axonal labeling within Pf. (**P**) Quantification of normalized GPe^Pf^ axon density across target regions (mean ± SEM, *n* = 6). (**Q–V**) Projection-specific monosynaptic rabies identification of GPe^PPN^ neurons. (**Q**) High-magnification image of labeled GPe^PPN^ neurons. (**R**) Experimental schematic showing viral injection and monosynaptic rabies tracing strategy to identify GPe^PPN^ neurons in *vGlut2-Cre* mice. (**S–U**) Representative coronal sections showing the absence of GPe^PPN^ collateral in Pf (**S**) and the presence of dense GPe^PPN^axon collaterals in PPN (**T**, **U**). Inset (**U**), magnified view of dense axonal labeling within PPN. (**V**) Quantification of normalized GPe^PPN^ axon density across target regions (mean ± SEM, n = 5 mice).

**S6M–S6X**). Together, these findings reveal that GPe^Pf^ and GPe^PPN^ neurons form largely segregated output channels that project in parallel to the thalamus and brainstem, respectively, with minimal collateral overlap.Our single-cell reconstructions demonstrate that the diverse projection patterns observed at the population level are recapitulated by individual GPe neurons. These single-neuron projectomes align closely with transcriptomically-defined cell types while also uncovering additional organization within clusters. Even within a given transcriptomic cell type, projection-specific subpopulations can be spatially segregated within the GPe and extend axons to largely non-overlapping downstream targets, supporting a model in which anatomically and molecularly defined GPe cell types form parallel and functionally distinct output channels (**Figures 3G, 4E, 4F,** and **S5**).

### GPe projection to PPN but not Pf modulates locomotion

Previous studies have identified the PPN as a locomotor command center responsible for initiating and maintaining locomotion, while the Pf has been implicated in the execution of learned forelimb action sequences^38–41^. Classic basal ganglia output structures SNr and GPi are thought to exert tonic inhibitory control over their downstream targets and can disinhibit them by reducing their firing rates^6,42,43^. To investigate how GPe projections to the PPN and Pf are engaged during behavior and whether they decrease their activity to relieve inhibition of their targets, we monitored the activity of GPe^Pf^ and GPe^PPN^ neurons during open field exploration. To achieve this, we expressed axon-targeted GCaMP6f in the GPe and implanted optical fibers above the axonal terminals of GPe projections to the PPN and Pf (**Figures 5A–5C**), enabling projection-specific fiber photometry recordings^44^. We collected continuous inertial sensor data and applied affinity propagation to identify behavioral clusters during open field^45,46^. To quantify how GPe outputs to the PPN and Pf are modulated during locomotion, we aligned calcium signals to the onset of locomotor bouts, defined as 600-ms epochs corresponding to two consecutive locomotion clusters (300 ms per cluster bin) identified through unsupervised behavioral clustering. We observed a robust decrease in the activity of GPe axons projecting to the PPN at the onset of locomotion (**Figure 5D**), consistent with a disinhibitory mechanism facilitating movement initiation. Direct statistical comparison revealed that GPe→PPN axons showed a significantly greater decrease in activity at locomotor onset than GPe→Pf axons (**Figures 5E** and **5F**).

**Figure 5.**
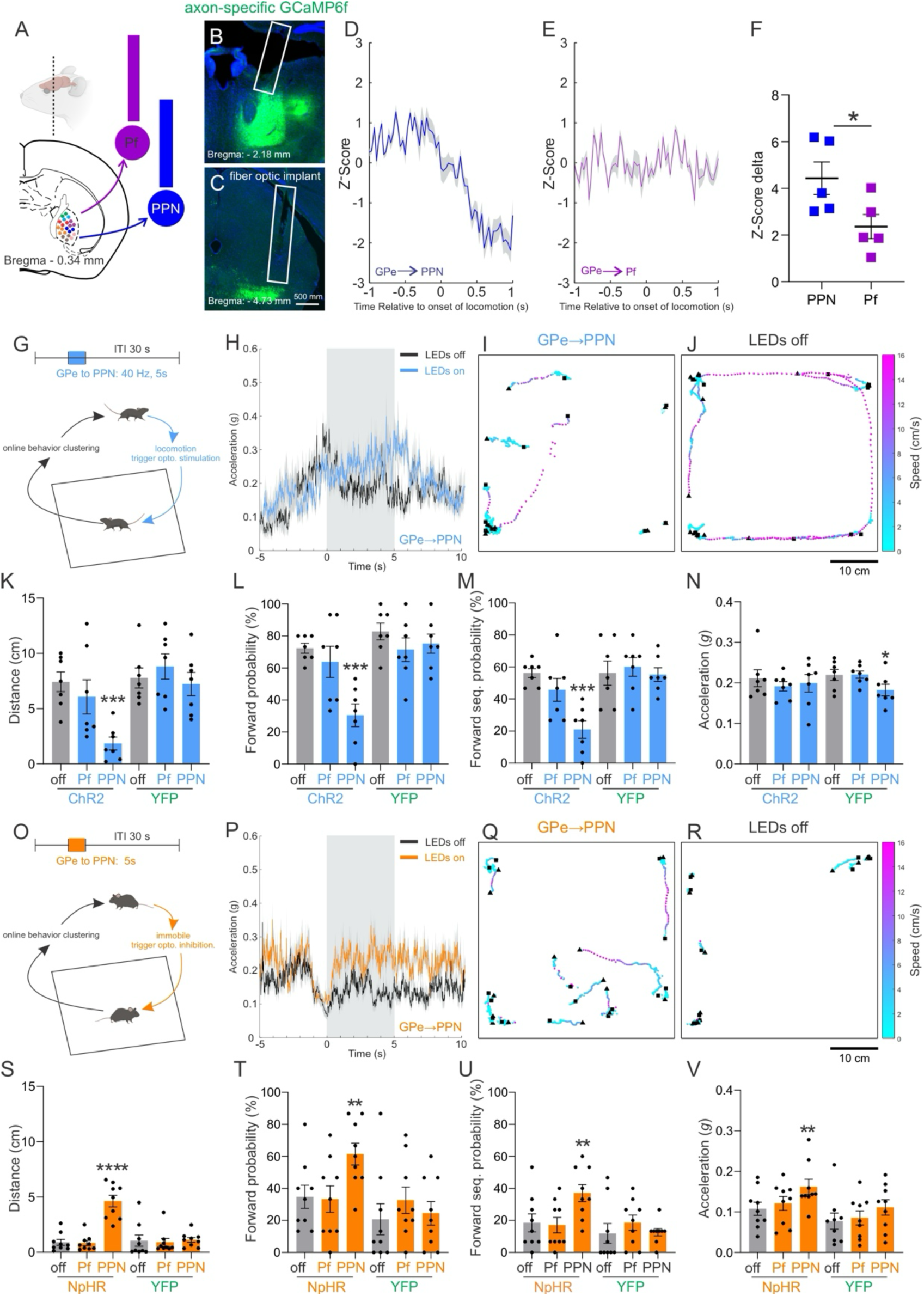
GPe→PPN but Not GPe→Pf Projections Modulate Locomotion. (**A–C**) Schematic (**A**) and histology showing projection-specific fiber-photometry targeting of GPe axons in Pf and PPN. (**B**, **C**) Representative sections showing GCaMP6f-labeled GPe axons and fiber-optic placements targeting Pf (**B**) and PPN (**C**) (n = 5 mice per group). (**D**, **E**) Average z-scored calcium activity aligned to locomotion onset for GPe→PPN (**D**) and GPe→Pf (**E**). Note the robust decrease in GPe→PPN activity at the onset of locomotion. (**F**) Quantification of Z-score delta at locomotion onset at locomotion onset (mean ± SEM; n = 5 mice in each group, P = 0.0437). (**G**) Closed-loop optogenetic stimulation paradigm for real-time detection of locomotion clusters and optogenetic activation of GPe→PPN or GPe→Pf axons (40 Hz, 5 s pulses; inter-trial interval = 30 s) during open field behavior. (**H**) Representative trace of total body acceleration during LED-on (blue) and LED-off (black) trials from a mouse expressing ChR2 in GPe neurons. The gray shaded interval indicates the period of optogenetic stimulation targeting GPe→PPN projections. **(I, J)** Representative plots showing centroid trajectories from 15 trials each of LED-on (**I**) and LED-off (**J**) conditions in a mouse expressing ChR2 in GPe neurons. Speed (cm/s) is color-coded along each trajectory; black triangles mark the start of each trial, and black squares mark the end of the 5 s stimulation or control period. (**K–N**) Quantification of distance (**K**), forward probability (**L**), forward sequence probability (**M**), and acceleration (**N**) across stimulation conditions (LED off, LED on Pf, LED on PPN) in ChR2 and YFP groups (ChR2: n = 7; YFP: n = 7). In ChR2 mice, optogenetic stimulation significantly reduced distance (F(1.399, 8.395) = 14.66, P = 0.0029; PPN: P = 0.0002; Pf: P = 0.4836), forward probability (F(1.851, 11.11) = 25.07, P < 0.0001; PPN: P = 0.0006; Pf: P = 0.4252), and forward sequence probability (F(1.977, 11.86) = 28.75, P < 0.0001; PPN: P = 0.0005; Pf: P = 0.1215), with no effect on acceleration (F(1.520, 9.119) = 0.3724, P = 0.6438). Effects were specific to PPN stimulation, with no effect of Pf stimulation. In YFP mice, no consistent effects were observed (distance: F(1.788, 10.73) = 0.5165, P = 0.5911; forward probability: F(1.487, 8.924) = 1.455, P = 0.2752; forward sequence probability: F(1.666, 9.994) = 0.1538, P = 0.8227; acceleration: F(1.416, 8.498) = 5.188, P = 0.0418). (**O**) Closed-loop optogenetic inhibition paradigm for real-time detection of immobile clusters and optogenetic inhibition of GPe→PPN or GPe→Pf axons (5 s continuous illumination; inter-trial interval = 30 s) during open field behavior. (**P**) Representative trace of total body acceleration during LED-on (orange) and LED-off (black) trials from a mouse expressing NpHR in GPe neurons. The gray shaded interval indicates the period of optogenetic inhibition targeting GPe→PPN projections. (**Q**, **R**) Representative plots showing centroid trajectories from 15 trials each of LED-on (**Q**) and LED-off (**R**) conditions in a mouse expressing NpHR in GPe neurons. Speed (cm/s) is color-coded along each trajectory; black triangles mark the start of each trial, and black squares mark the end of the 5 s inhibition or control period. (**S–V**) Quantification of distance (**S**), forward probability (**T**), forward sequence probability (**U**), and acceleration (**V**) across inhibition conditions (LED off, LED on Pf, LED on PPN) in NpHR and YFP groups (NpHR: n = 9; YFP: n = 9). In NpHR mice, optogenetic inhibition significantly increased distance (F(1.366, 10.93) = 37.13, P < 0.0001; PPN: P < 0.0001; Pf: P = 0.9777), forward probability (F(1.282, 10.26) = 4.316, P = 0.0567; PPN: P = 0.0028; Pf: P = 0.9885), forward sequence probability (F(1.178, 9.421) = 5.048, P = 0.0458; PPN: P = 0.0004; Pf: P = 0.9785), and acceleration (F(1.525, 12.20) = 6.543, P = 0.0162; PPN: P = 0.0016; Pf: P = 0.6618), with effects driven by inhibition of PPN-projecting neurons and not Pf-projecting neurons. In YFP mice, no consistent effects of optogenetic inhibition were observed (distance: F(1.100, 8.803) = 0.1094, P = 0.7724; forward probability: F(1.493, 11.95) = 0.6043, P = 0.5161; forward sequence probability: F(1.493, 11.95) = 0.6043, P = 0.5161; acceleration: F(1.964, 15.71) = 1.096, P = 0.3575). Data are mean ± SEM. ****P < 0.0001, ***P < 0.001, **P < 0.01, *P < 0.05; ns, not significant.

To probe the functional relevance of GPe projections to the PPN and Pf, we implemented a closed-loop bidirectional optogenetic manipulation strategy (**Figures 5G** and **5O**). We first performed offline clustering of open field behavior to build individualized libraries of behavioral clusters for each mouse. We then extracted video frames corresponding to specific behavioral clusters and compiled cluster-specific videos to visually confirm the identity of individual clusters identified through unsupervised clustering. During online sessions, we deployed a computationally efficient clustering and matching pipeline to continuously identify target behavioral clusters in real time^46^. Upon detection of a predefined cluster, specifically locomotion or immobility, we triggered optogenetic manipulation of GPe output projections. This approach allowed us to assess the contributions of GPe^PPN^and GPe^Pf^pathways to both locomotor execution (triggered by the locomotion cluster) and locomotor initiation (triggered by the immobility cluster).

To test whether GPe-mediated disinhibition is necessary for locomotor execution, we expressed channelrhodopsin-2 (ChR2–YFP) or control YFP bilaterally in the GPe and implanted fiber optics above the axonal terminals in either the PPN or Pf. Our real-time closed-loop system reliably detected locomotion clusters and triggered optogenetic stimulation (40 Hz, 5 s duration, 30 s inter-trial interval) (**Figure 5G**). We found that optogenetic enhancement of GPe→PPN inhibition significantly reduced locomotor distance traveled compared to LED-off trials, whereas stimulation of GPe→Pf projections had no significant effect (**Figures 5I–5K**; **Video S4**). Further analysis revealed that stimulation of GPe→PPN projections significantly decreased the probability of sustaining locomotor clusters and disrupted the formation of locomotor sequences (defined as two consecutive bins of locomotion clusters, 600 ms total) (**Figures 5L** and **5M**). In contrast, GPe→Pf stimulation did not significantly affect these behavioral metrics (**Figures 5K–5M**; **Video S4**). Importantly, neither manipulation altered overall body acceleration (**Figures 5H** and **5N**), indicating that GPe→PPN influence on locomotor execution is not simply due to changes in general movement or vigor but rather reflects modulation of specific cluster of action. These results suggest a selective role for GPe^PPN^ in locomotor control. In the control YFP group, LED illumination of the PPN had no detectable impact on locomotor behavior. Specifically, optogenetic stimulation did not alter the distance traveled, the probability of sustaining locomotor clusters, the formation of locomotor sequences, or total body acceleration when comparing LED-on versus LED-off trials (**Figures 5K–5N**; **Video S5**). Similarly, stimulation of GPe→Pf terminals in YFP-expressing mice failed to produce any noticeable changes in these behavioral metrics (**Figures 5K–5N**; **Video S5**). These control experiments confirm that the observed behavioral effects are not due to light delivery or nonspecific activation. Taken together, these results strongly support the conclusion that decreased GPe-mediated inhibition of the PPN but not the Pf is necessary for the proper execution of locomotion.

To test whether reducing GPe-mediated inhibition of the PPN is sufficient to induce locomotion, we expressed Halorhodopsin (NpHR–YFP) or control YFP bilaterally in the GPe and implanted fiber optics above the axonal terminals in either the PPN or Pf. Our real-time closed-loop system reliably detected immobility clusters and triggered optogenetic inhibition via continuous yellow light illumination (5 s duration, 30 s inter-trial interval) (**Figures 5O** and **5P**). We found that optogenetic inhibition of the GPe→PPN projection, thereby releasing the PPN from GPe-mediated inhibition, significantly increased locomotor distance traveled compared to LED-off trials, whereas inhibition of the GPe→Pf projection had no significant effect (**Figures 5Q–5S**; **Video S6**). Furthermore, GPe→PPN inhibition increased the probability of initiating locomotor clusters and enhanced the formation of locomotor sequences, defined as two consecutive bins of locomotor clusters (600 ms total) (**Figures 5T** and **5U**). In contrast, optogenetic inhibition of GPe→Pf terminals did not significantly alter these behavioral measures (**Figures 5T** and **5U**; **Video S6**). Notably, GPe→PPN inhibition produced a robust increase in total body acceleration (**Figure 5V**), suggesting that relieving GPe-mediated inhibition of the PPN not only promotes locomotor initiation but also invigorates movement. In control mice expressing YFP alone, LED illumination of the PPN failed to elicit any locomotor effects: there were no changes in distance traveled, probability of initiating locomotor clusters, sequence formation, or body acceleration between LED-on and LED-off trials (**Figures 5S–5V**; **Video S7**). Similarly, illumination of Pf terminals in YFP controls did not produce significant changes in these parameters (**Figures 5S–5V**; **Video S7**). Together, these data demonstrate that reducing GPe inhibition of the PPN but not Pf is sufficient to facilitate locomotion. These findings show that disinhibition via GPe→PPN, but not GPe→Pf projections, plays a critical role in the initiation and execution of locomotor behavior, and demonstrate a specific direct effect of particular GPe populations on brainstem targets.

### A “direct” D2-MSN striatopallidal pathway to GPe^PPN^ neurons modulates locomotion

Our findings reveal that GPe projections to the PPN exhibit a classic disinhibitory pattern during locomotion, reminiscent of the well-established mechanisms by which the SNr and GPi regulate downstream targets through tonic inhibition and transient pauses in firing to facilitate movement^42,47^. This observation raises an important conceptual prediction: if GPe, as a downstream target of D2-MSNs (classically named as the origin of the indirect pathway), can drive disinhibition of basal ganglia downstream targets like the PPN, then D2-MSNs themselves may contribute to movement initiation through mechanisms classically ascribed to the direct (D1-MSNs) pathway. To directly test this hypothesis, we next investigated whether a direct striatopallidal pathway from D2-MSNs to the PPN exists and is sufficient to drive both the initiation and maintenance of locomotion.

We first asked whether the segregated GPe^PPN^ and GPe^Pf^populations receive inputs from overlapping or distinct sets of MSNs by performing cell type- and projection-specific monosynaptic rabies tracing from each GPe subpopulation. Specifically, we injected *AAVretro-EF1a-Flpo* into either the PPN or Pf to express Flpo recombinase in GPe^PPN^ or GPe^Pf^neurons, respectively (**Figure 6A**). In the same animals, we delivered a mixture of *AAV1.EF1a.FlpX.H2B.3XHA.2A.N2C(G)* and *AAVDJ.hSyn.FlpX.TVA950* into the GPe to drive Flpo-dependent expression of the avian TVA950 receptor and rabies glycoprotein N2C(G), elements required for selective infection and transsynaptic labeling using pseudotyped rabies virus^36^. Two weeks later, we injected an EnvA-pseudotyped, G-deleted CVS-N2c rabies virus encoding GFP (EnvA-N2cΔG-GFP) into the GPe (**Figures 6A and 6B**). This approach enabled retrograde labeling of striatal MSNs that provide monosynaptic input to either GPe^PPN^ or GPe^Pf^neurons. Following immunohistochemical enhancement of GFP and serial coronal section imaging, we performed quantitative analysis using BrainJ. Registration to the CCFv3 and 3D reconstruction with Brainrender revealed sparse but topographically specific labeling of MSNs (**Figures 6C, 6D, and S7E–S7H**). Notably, MSNs labeled from GPe^PPN^ (i.e., MSNs→GPe^PPN^→PPN pathway; hereafter referred to MSNs-GPe^PPN^) and from GPe^Pf^ (i.e., MSNs→GPe^Pf^→Pf pathway; MSNs-GPe^Pf^) were spatially segregated, with the most prominent separation observed in DLS (**Figures 6**C**–6E, and S7E–S7H**). Quantification showed a significantly greater fraction of MSNs-GPe^PPN^neurons located laterally compared to MSNs-GPe^Pf^neurons (**Figure 6E**). Together, these results (**Figures 4** and **6**) demonstrate that GPe^PPN^ and GPe^Pf^neurons form anatomically segregated output channels that project to distinct brainstem and thalamic targets with minimal collateralization and that these GPe subpopulations receive input from spatially distinct MSN populations.

**Figure 6.**
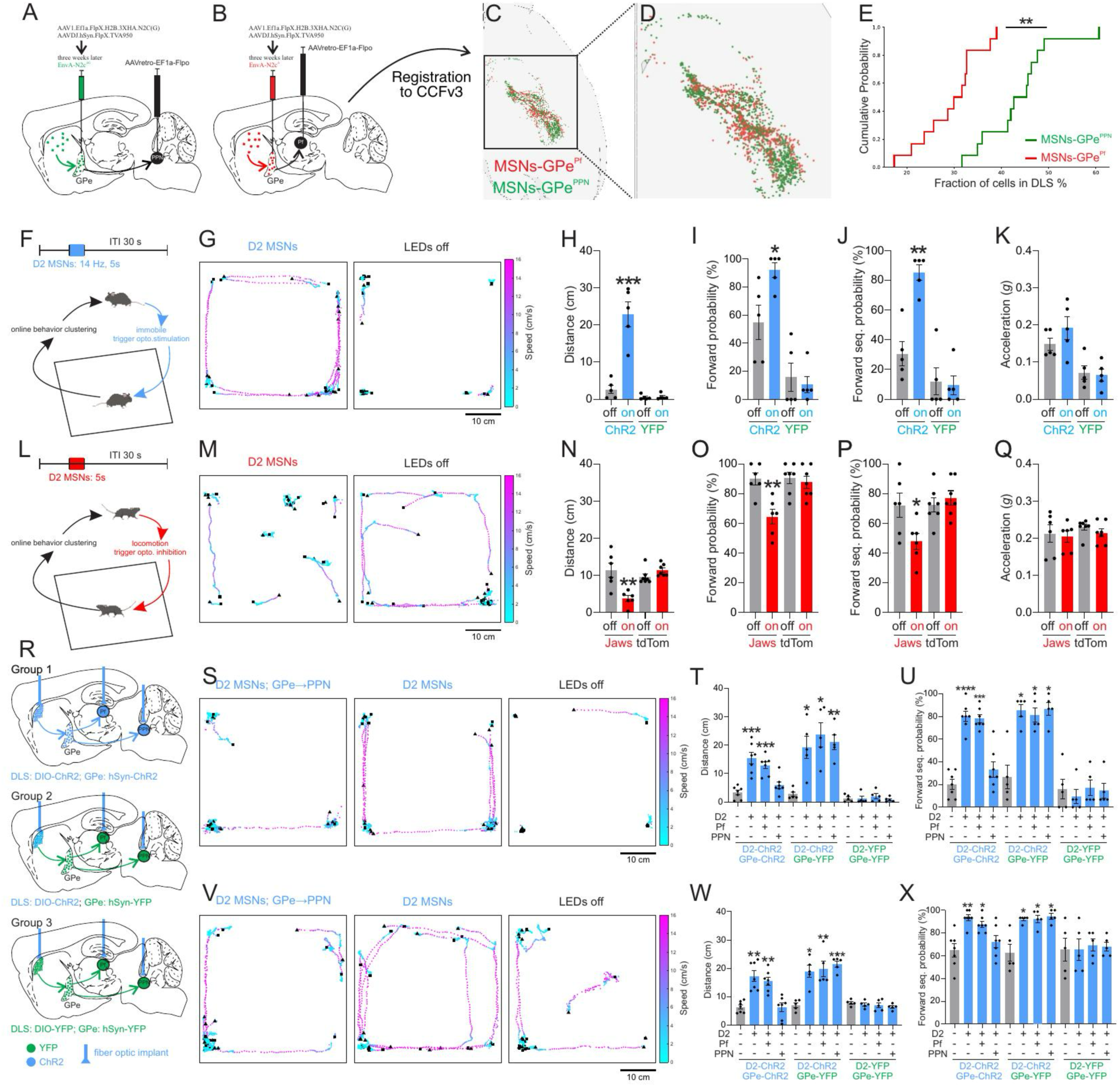
A Direct D2-MSNs Pathway to the PPN that Initiates and Sustains Locomotion. (**A–B**) Cell-type- and projection-specific monosynaptic rabies tracing strategy to identify striatal MSN inputs to GPe neurons projecting to the PPN and Pf. AAVretro-EF1α-Flpo was injected into the PPN or Pf to label GPe^PPN^ or GPe^Pf^ neurons, respectively. In the GPe, Flpo-dependent helper viruses (AAV1.EF1a.FlpX.H2B.3XHA.2A.N2C(G) and AAVDJ.hSyn.FlpX.TVA950) enabled selective expression of TVA and rabies glycoprotein. Three weeks later, EnvA-N2cΔG-GFP was injected into the GPe to label monosynaptic striatal inputs. (**C–D**) BrainJ-based cell detection and registration to Allen CCFv3 with 3D reconstruction of rabies-labeled striatal MSNs projecting to GPe^PPN^ (green) and GPe^Pf^ (red), revealing spatially segregated input populations. (**D**) shows a magnified view of the boxed region in (**C**). (**E**) Cumulative distribution analysis of MSN fractions in the DLS reveals a significant lateral shift in MSNs-GPe^PPN^ (green) relative to MSNs-GPe^Pf^ (red) (Kolmogorov–Smirnov test: D = 0.444, P = 0.0014). **(F–K)** Closed-loop optogenetic activation of D2-MSNs in DLS, triggered by real-time detection of immobility clusters, induces locomotion. (**F**) Schematic of closed-loop stimulation paradigm: immobility clusters detected in real time trigger 14 Hz, 5 s blue-light stimulation (30 s inter-trial interval) of D2-MSNs in DLS. (**G**) Representative plots showing centroid trajectories from 15 trials each of LED-on (left) and LED-off (right) conditions in a mouse expressing ChR2 in D2-MSNs. Speed (cm/s) is color-coded along each trajectory; black triangles mark the start of each trial, and black squares mark the end of the 5 s stimulation or control period. (**H–K**) Quantification of locomotor behaviors in ChR2 and YFP mice across conditions (LED off, LED on). Activation of D2-MSNs significantly increased distance traveled (H), locomotor cluster initiation probability (I), and locomotor sequence probability (J), with no effect on acceleration (K) (mean ± SEM; n = 5 mice per group). In the ChR2 group, two-way repeated-measures ANOVA revealed a significant effect of optogenetic stimulation on distance traveled (F(1,8) = 36.44, P = 0.0003), locomotor probability (F(1,8) = 3.716, P = 0.0900, with significant post hoc), and sequence probability (F(1,8) = 25.35, P = 0.0010), but not acceleration (F(1,8) = 0.6230, P = 0.4527). Sidak’s multiple comparisons test showed that LED stimulation significantly increased distance traveled (P < 0.0001), locomotor probability (P = 0.0258), and sequence probability (P = 0.0001), with no effect on acceleration (P = 0.3935). In the YFP group, no significant effect of optogenetic stimulation was observed across measures (all P > 0.45), and LED stimulation did not alter distance traveled (P = 0.9971), locomotor probability (P = 0.8855), sequence probability (P = 0.9241), or acceleration (P = 0.9739). **(L–Q)** Closed-loop optogenetic inhibition of D2-MSNs during locomotion attenuates ongoing locomotion. **(L)** Schematic of closed-loop inhibition paradigm: locomotor clusters detected in real time trigger 5 s continuous red-light illumination (30 s inter-trial interval) delivered at locomotor onset. **(M)** Representative plots showing centroid trajectories from 15 trials each of LED-on (left) and LED-off (right) conditions in a mouse expressing Jaws in D2-MSNs. Speed (cm/s) is color-coded along each trajectory; black triangles mark the start of each trial, and black squares mark the end of the 5 s stimulation or control period. **(N–Q)** Inhibition of D2-MSNs (Jaws) significantly reduces distance traveled (**N**), probability of sustaining locomotor clusters (**O**), probability of forming locomotor sequence (**P**) without affecting acceleration (**Q**). Tdtomato controls show no effect. (**N–Q**) Inhibition of D2-MSNs (Jaws) reduces locomotor behavior (Jaws: n = 6 mice; tdTomato: n = 7 mice). Distance traveled (N), forward locomotion probability (O), and locomotor sequence probability (P) were significantly reduced during LED inhibition in Jaws mice, with no effect observed in tdTomato controls. In contrast, acceleration (Q) was not altered by inhibition. Two-way repeated-measures ANOVA revealed significant effects of optogenetic inhibition on distance traveled (N: F(1,11) = 12.70, P = 0.0044) and forward locomotion probability (O: F(1,11) = 14.49, P = 0.0029), as well as a significant group effect and interaction for locomotor sequence probability (P: group F(1,11) = 5.634, P = 0.0369; interaction F(1,11) = 7.353, P = 0.0202). No significant effects were observed for acceleration (Q: all P > 0.11). Sidak’s post hoc comparisons confirmed LED-dependent reductions in Jaws mice (N: P = 0.0001; O: P = 0.0014; P: P = 0.0204), with no effect in tdTomato controls (all P > 0.30). Acceleration was unchanged in both groups (Jaws: P = 0.6940; tdTomato: P = 0.2446). **(R–X)** Concurrent manipulation of GPe output identifies a D2-MSNs →GPe→PPN disinhibitory pathway mediating locomotion initiation and maintenance. (**R**) Experimental design for three groups: (Group 1, top) dual ChR2 expression in D2-MSNs and GPe neurons with optical fibers over DLS, PPN, and Pf terminals; (Group 2, middle) ChR2 in D2-MSNs with GPe-YFP control; (Group 3, bottom) dual YFP control. **(S–U)** Across all groups, immobility-triggered D2-MSN stimulation induces locomotion. In Group 1, this effect is selectively attenuated by co-activation of GPe→PPN but not GPe→Pf terminals, whereas no projection-specific effects are observed in control groups. (Group 1: n = 7 mice; Group 2: n = 5 mice; Group 3: n = 5 mice). (**S**) Representative centroid trajectories from 15 trials under LED-on concurrent stimulation of D2-MSNs and GPe→PPN terminals (left), D2-MSN stimulation alone (middle), and LED-off (right) conditions in a mouse expressing ChR2 in D2-MSNs and GPe neurons. Stimulation was triggered by immobility clusters. Trajectories are color-coded by speed (cm/s). Black triangles indicate trial onset, and black squares mark the end of the 5 s stimulation or control window. (**T**) Quantification of distance traveled (cm) during immobility-triggered stimulation. In Group 1 (D2-MSN-ChR2 + GPe-ChR2), repeated-measures one-way ANOVA revealed a significant effect of stimulation (F(1.990, 11.94) = 24.91, P < 0.0001). D2-MSN stimulation increased distance (P = 0.0005), which was maintained with GPe→Pf activation (P = 0.0001), but reduced to baseline by GPe→PPN activation (P = 0.2164). Sidak comparisons confirmed suppression by GPe→PPN relative to D2-only and D2+Pf (P = 0.0097 and P = 0.0472), with no difference between D2-only and D2+Pf (P = 0.8749). In Group 2 (D2-ChR2 + GPe-YFP), stimulation increased distance across all conditions (F(1.547, 6.190) = 9.845, P = 0.0143; DLS: P = 0.0223; Pf: P = 0.0251; PPN: P = 0.0026). In Group 3 (D2-YFP + GPe-YFP), no effects were observed (F(1.618, 6.473) = 0.5030, P = 0.5897; all P > 0.82). (**U**) Quantification of forward sequence probability (%) during immobility-triggered stimulation. In Group 1, stimulation had a significant effect (F(2.409, 14.46) = 37.69, P < 0.0001): D2-MSN activation increased sequence probability (P < 0.0001), which was preserved with GPe→Pf (P = 0.0011) but abolished by GPe→PPN (P = 0.4845). Sidak comparisons confirmed suppression by GPe→PPN relative to D2-only and D2+Pf (P = 0.0090 and P = 0.0073), with no difference between D2-only and D2+Pf (P > 0.9999). In Group 2, stimulation increased sequence probability (F(1.267, 5.068) = 14.51, P = 0.0105; DLS: P = 0.0175; PPN: P = 0.0403), while Pf showed a non-significant trend (P = 0.0500). In Group 3, no effects were observed (F(2.095, 8.379) = 0.2335, P = 0.8060; all P > 0.37). **(V–X)** When stimulation is delivered at locomotion onset, D2-MSNs activation enhances ongoing locomotion (**V–X**), an effect reversed by co-activation of GPe→PPN but not GPe→Pf terminals. (**V**) Representative centroid trajectories from 15 trials under LED-on concurrent stimulation of D2-MSNs and GPe→PPN terminals (left), D2-MSN stimulation alone (middle), and LED-off (right) conditions in a mouse expressing ChR2 in D2-MSNs and GPe neurons. Stimulation was triggered by locomotor clusters. Trajectories are color-coded by speed (cm/s). Black triangles indicate trial onset, and black squares mark the end of the 5 s stimulation or control window. (**W**) Quantification of distance traveled (cm) across stimulation conditions (LED off, DLS, Pf, PPN). In Group 1, stimulation had a significant effect (F(2.421, 14.53) = 15.67, P = 0.0001): D2-MSN activation increased distance (P = 0.0024), which was maintained with GPe→Pf (P = 0.0033) but abolished by GPe→PPN (P > 0.9999). In Group 2, stimulation increased distance across all conditions (F(1.541, 6.164) = 15.73, P = 0.0048; DLS: P = 0.0251; Pf: P = 0.0091; PPN: P = 0.0007). In Group 3, no effects were observed (F(1.822, 7.289) = 0.7173, P = 0.5071; DLS: P = 0.8820; Pf: P = 0.6484; PPN: P = 0.3376). (**X**) Quantification of forward sequence probability (%) across stimulation conditions (LED off, DLS, Pf, PPN). In Group 1, stimulation had a significant effect (F(1.763, 10.58) = 8.098, P = 0.0086): D2-MSN activation increased sequence probability (P = 0.0054), which was maintained with GPe→Pf (P = 0.0114) but not with GPe→PPN (P = 0.7434). In Group 2, stimulation had a significant effect (F(1.323, 5.293) = 12.11, P = 0.0136), with all conditions increasing sequence probability relative to LED off (DLS: P = 0.0490; Pf: P = 0.0498; PPN: P = 0.0447). In Group 3, no effects were observed (F(2.223, 8.891) = 0.1353, P = 0.8935; all P > 0.95). Data are mean ± SEM. ****P < 0.0001, ***P < 0.001, **P < 0.01, *P < 0.05; ns, not significant.

Given these spatial biases, we next tested whether activation of D2-MSNs in DLS projecting to GPe^PPN^ is sufficient to initiate locomotion (**Figure 6F**), and conversely, whether inhibition of D2-MSNs in this area suppresses locomotion (**Figure 6L**). We again used a state-dependent, closed-loop bidirectional optogenetic strategy targeting the D2-MSNs in the specific area of DLS. We took advantage of the real-time behavioral clustering and matching approach described above to continuously identify immobility or locomotor onset clusters (**Figures 6F** and **6L**)^46^, and precisely time D2-MSN manipulations upon the identification of targeted clusters. We bilaterally injected *AAV1.EF1a.DIO.hChR2(H134R)-eYFP* (ChR2–YFP) or *AAV1.EF1a.DIO.eYFP* (control) into the DLS of A2A-Cre mice and implanted fiber optics over the same region. Using our real-time closed-loop system, we detected immobility clusters and triggered optogenetic stimulation of D2-MSNs with a stimulation protocol previously validated (14 Hz, 5 s duration, 30 s inter-trial interval)^8,46^ (**Figure 6F**). We found that optogenetic activation of D2-MSNs reliably evoked robust locomotion, significantly increasing distance traveled compared to LED-off trials (**Figures 6G** and **6H**; **Video S8**). Further analysis showed that D2-MSN stimulation in this region of DLS significantly increased the probability of initiating locomotor clusters and promoted the formation of locomotor sequences, without affecting overall body acceleration (**Figures 6I–6K**). In contrast, in control animals expressing eYFP, blue LED illumination of the DLS produced no detectable behavioral effects: distance traveled, initiation probability of locomotor clusters, locomotor sequence formation probability, and body acceleration all remained unchanged between LED-on and LED-off conditions (**Figures 6H–6K**; **Video S9**). Thus, this state-dependent, closed-loop stimulation of D2-MSNs in the DLS during immobility reliably triggered locomotor initiation. We next sought to determine whether D2-MSN activity is necessary to sustain locomotion once initiated. To this end, we bilaterally injected *AAV8-hSyn-Flex-Jaws-GFP* (Jaws) into the DLS of *A2A-Cre* mice and implanted fiber optics over the same region. Control animals received either *AAV1-CAG-FLEX-tdTomato* or *AAV1-CAG-FLEX-GFP*. We delivered optogenetic inhibition at the onset of locomotion (continuous red light, 5 s duration, 30 s inter-trial interval; **Figure 6L; Video S10**). We found that the inhibition of D2-SPNs significantly attenuated ongoing locomotion, reducing distance traveled relative to LED-off trials (**Figures 6M** and **6N**; **Video S10**). Additional analysis revealed that D2-MSN inhibition significantly decreased the probability of sustaining locomotor clusters and disrupted the formation of locomotor sequences, without affecting overall total body acceleration (**Figures 6O–6Q**). In contrast, red LED illumination in control animals had no effect on locomotor metrics: distance traveled, cluster initiation probability, locomotor sequence formation probability, and total body acceleration all remained unchanged between LED-on and LED-off conditions (**Figures 6N–6Q**; **Video S11**). Together, these results demonstrate that D2-MSN activity in the region of DLS projecting to GPe^PPN^ neurons is sufficient and necessary for locomotor behavior.

We next tested whether the effects of D2-MSN activation on locomotion were mediated by GPe, and more by specific GPe projections. To achieve this, we simultaneously activated GPe^PPN^ or GPe^Pf^ neurons upon D2-MSN activation, reasoning that if D2-MSN mediated locomotion occurs via inhibition of one of these downstream projections, their simultaneous activation would block D2-MSN mediated locomotion. We designed an experimental paradigm with three groups of mice (**Figure 6R**). In the first group, we bilaterally injected *AAV1.EF1a.DIO.hChR2(H134R)-eYFP* into the DLS and *AAV1-hSyn-hChR2(H134R)-eYFP* into the GPe of *A2A-Cre* mice. We then implanted fiber optics over the DLS and GPe axonal terminals in the PPN and Pf, enabling concurrent optogenetic activation of D2-MSNs and elevation of GPe inhibition to downstream targets in the same mouse (**Figure 6R**, top). In the second group (ChR2 in D2-MSN only group), we injected *AAV1.EF1a.DIO.hChR2(H134R)-eYFP* into the DLS and *AAV1-hSyn-eYFP* into the GPe (**Figure 6R**, middle). This allowed for selective activation of D2-MSNs while controlling for nonspecific effects of light delivery to GPe axonal terminals. In the third group (full control), we injected *AAV1.EF1a.DIO.eYFP* into the DLS and *AAV1-hSyn-eYFP* into the GPe (**Figure 6R**, bottom). These mice received light illumination to both DLS and GPe axonal terminals but expressed YFP instead of opsin, providing a baseline control for photo illumination effects. These three groups of mice allowed us to determine whether the behavioral effects of D2-MSN activation are mediated specifically via disinhibition of the PPN through a GPe subcircuit. In Group one, we first confirmed that optogenetic stimulation of D2-MSNs reliably evoked robust locomotion when stimulation was triggered by immobility cluster (**Figure S8A**), significantly increasing distance traveled compared to LED-off trials (**Figures 6R–6T**, and **S8B**; **Video S12**). This activation also significantly increased the probability of initiating locomotor clusters and promoted the formation of locomotor sequences, without altering overall total body acceleration (**Figures 6U**, **S8C**, and **S8D**). Within the same session and in the same animals, we next performed trials in which stimulation of D2-MSNs was paired with simultaneous optogenetic activation of GPe axon terminals targeting either the PPN or Pf, thereby enhancing GABAergic inhibition onto these downstream targets while we stimulated D2-MSNs in DLS. We observed that D2-MSN-evoked locomotion was significantly attenuated by concurrent stimulation of GPe→PPN projections but not GPe→Pf projections (**Figure 6S and 6T**; **Video S12**). Specifically, GPe→PPN activation reduced the distance traveled to levels comparable to LED-off baseline trials (**Figures 6S** and **6T**; **Video S12**). Further analysis showed that only concurrent stimulation of GPe→PPN but not GPe→Pf significantly reduced the elevated probability of initiating locomotor clusters and the increased formation of locomotor sequences evoked by D2-SPN stimulation alone (**Figures 6U**, **S8C**, and **S8D**; **Video S12**). Importantly, none of these manipulations affected overall total body acceleration (**Figure S8C**), indicating a specific modulation of locomotor clusters rather than general motor output. In contrast, in group 2 mice expressing only eYFP in GPe axon terminals, blue LED illumination of the PPN or Pf terminals had no effect on D2-MSN-evoked locomotion (**Figure 6T**; **Video S13**). There were no significant changes in distance traveled, locomotor cluster initiation, sequence formation, or total body acceleration between LED-on and LED-off conditions (**Figures 6T**, **6U**, **S8C**, and **S8D**). Similarly, in group 3 animals expressing YFP in both D2-MSNs and GPe neurons, blue LED illumination of the DLS alone or in combination with GPe terminal illumination failed to alter behavior (**Figures 6T**, **6U**, **S8C**, and **S8D**; **Video S14**). Together, these findings demonstrate that locomotion evoked by D2-MSN activation is selectively channeled through GPe output to the PPN, but not to the Pf. These results support the model that D2-MSNs initiate locomotion via projection-specific disinhibition of the PPN through a dedicated GPe subcircuit. We next asked whether stimulation of D2-MSNs in the DLS at locomotion onset could enhance ongoing movement (**Figure S8E**), and whether this effect is mediated by the same GPe→PPN pathway that supports movement initiation. In group 1 animals, we first confirmed that optogenetic stimulation of D2-SPNs triggered at locomotion onset further enhanced ongoing movement compared to LED-off trials (**Figure 6V**). This enhancement was reflected by increased distance traveled, higher probability of sustaining locomotor clusters, and greater probability of forming locomotor sequences, without changes in overall body acceleration (**Figures 6V–6X**, and **S8F–S8H**). Within the same session and animals, we then performed trials in which D2-MSN stimulation was paired with simultaneous optogenetic activation of GPe axon terminals in either the PPN or Pf, enhancing GPe GABAergic output to these respective targets. Strikingly, concurrent activation of GPe→PPN projections significantly attenuated the pro-locomotor effects of D2-SMSN stimulation, reducing distance traveled to levels comparable to LED-off baseline trials (**Figures 6V** and **6W**). Additional analysis revealed that only GPe→PPN but not GPe→Pf concurrent stimulation reversed the D2-MSN induced increase in the probability of sustaining locomotor cluster and forming locomotor sequences (**Figures 6V–6X**, and **S8F-S8H**). None of these manipulations affected total body acceleration (**Figures S8F** and **S8G**), suggesting a selective modulation of locomotor clusters rather than general motor output. In contrast, group 2 animals expressing eYFP in GPe terminals showed no changes of D2-MSN enhanced behavior with LED illumination of the PPN or Pf during D2-SPN stimulation. Locomotor metrics, including distance traveled, cluster continuation probability, sequence formation probability, and total body acceleration, were unchanged between LED-on and LED-off conditions during D2-MSN stimulation paired with blue LEDs illumination on GPe axonal terminal trials (**Figures 6W**, **6X**, and **S8F–S8H**). Similarly, in group 3 animals expressing YFP in both D2-MSNs and GPe neurons, locomotion-triggered illumination of the DLS alone or combined with GPe terminal stimulation did not alter behavior (**Figures 6W**, **6X**, and **S8F–S8H**).

These temporally precise, state-dependent, closed-loop optogenetic manipulations, combined with rabies tracing data showing that GPe^PPN^ and GPe^Pf^ neurons receive input from spatially distinct SPN populations, provide strong evidence that a subset of D2-MSNs facilitates locomotion through selective engagement of a GPe→PPN output channel, establishing a functional direct D2-MSN striatopallidal pathway to the PPN that supports both the initiation and maintenance of locomotion.

### GPe projections to Pf but not PPN modulate the execution of forelimb sequences

Our imaging and functional perturbation studies revealed a projection- and action-specific role for GPe→PPN outputs in locomotion: only GPe→PPN activity decreased at the onset of locomotion, and only suppression of GPe→PPN (but not GPe→Pf) facilitated locomotion initiation and execution (**Figure 5**). Prior studies have implicated the Pf in skilled motor behaviors, including the execution of learned forelimb action sequences^38^. We therefore investigated how the activity of GPe axons projecting to Pf and PPN are modulated during the performance of a trained forelimb lever-press sequence, and whether these projections show differential activity patterns aligned to the execution of learned, skilled forelimb behavior. We used the same cohort of mice (n = 5) in which axon-targeted GCaMP6f was expressed in the GPe, and optical fibers were implanted above GPe axonal terminals in the PPN and Pf (**Figure 7A**). These mice were trained to perform a rapid lever-pressing task requiring a sequence of eight presses under a fixed ratio 8 schedule (FR8) to receive a drop of 10% sucrose reward (**Figure 7B**). The training paradigm began with 2 days of magazine training, during which mice learned to retrieve the sucrose reward from the reward port. This was followed by 7 days of continuous reinforcement (CRF; one press = one reward), consisting of 2 days each of CRF5 and CRF15, and 3 days of CRF30, with increasing maximum reward limits per session. Following CRF, animals were trained under a progressively demanding fixed-ratio schedule: 2 days of FR4 (4 presses = one reward), followed by 14 days of FR8 training. During this period, mice exhibited steady increases in total lever presses and presses per minute and progressively began to structure their behavior into self-paced bouts or action sequences. After 14 days of FR8 training, we conducted fiber photometry recordings during task performance, in which mice were required to complete FR8 lever-press sequences to earn sucrose rewards (maximum of 30 rewards per session). This preparation enabled us to monitor GPe axonal activity in the PPN and Pf during the execution of well-trained sequences of self-initiated forelimb actions (**Figures 7C** and **7D**). We observed a pronounced decrease in the activity of GPe→Pf axons at the onset of a lever press sequence (**Figure 7D**), consistent with a disinhibitory mechanism facilitating lever presses. Direct statistical comparison revealed that GPe→Pf axons exhibited a significantly greater decrease in activity during lever press sequence execution than GPe→PPN axons (**Figures 7C** and **7E**).

**Figure 7.**
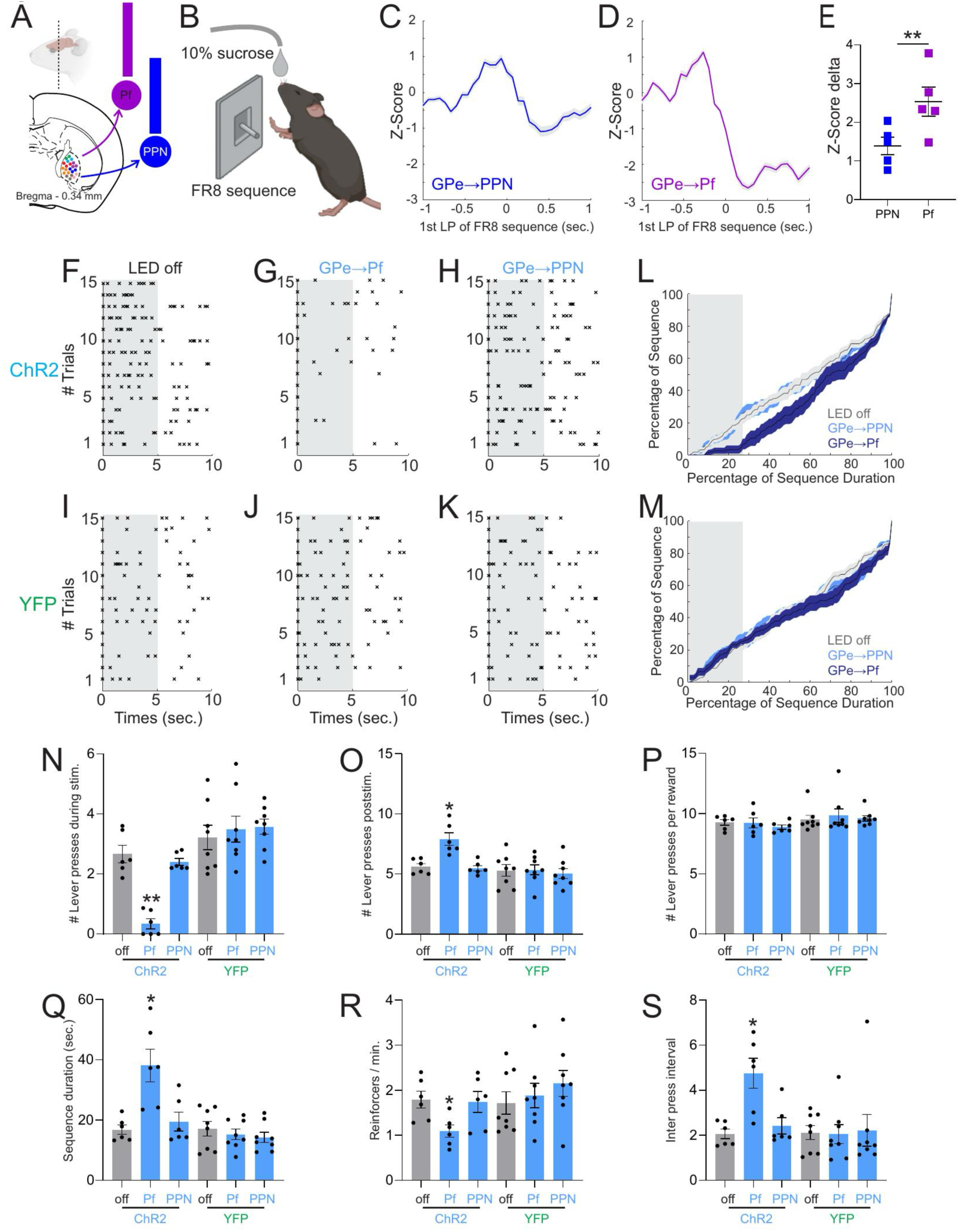
GPe Projection to Pf but Not PPN Modulates Skilled Forelimb Behavior. (**A**) Illustration of the fixed-ratio (FR8) lever-press task, in which mice were required to complete eight consecutive lever presses (fixed ratio 8) to obtain a drop of 10% sucrose reward. (**B**) Schematic of the experimental setup for projection-specific fiber photometry recordings from GPe axon terminals in the Pf (purple) and PPN (blue). (**C**, **D**) Example traces of GPe→PPN (**C**) and GPe→Pf (**D**) axonal calcium activity aligned to the first lever of FR8 lever pressing sequence (time 0). (**E**) Quantification of Z-score delta during lever press sequence onset, showing a significantly greater decrease in GPe→Pf than in GPe→PPN terminals (mean ± SEM; n = 5 mice per group; P = 0.015). **(F–H)** Raster plots of FR8 lever-press sequences during LED-off (**F**) and LED-on trials for GPe→Pf (**G**) and GPe→PPN (H) optogenetic stimulation in ChR2-expressing mice. The gray shaded region denotes the 5-second epoch of 40 Hz blue-light illumination. Notably, lever pressing was largely suppressed during stimulation in GPe→Pf trials, whereas LED-off or GPe→PPN trials showed normal lever-pressing performance. (**L**) Cumulative distributions showing the percentage of lever-press sequences completed over normalized sequence duration during LED-on and LED-off trials in ChR2-expressing mice. The gray shaded region denotes the 5-second epoch of 40 Hz blue-light illumination. Notably, lever pressing was largely suppressed during stimulation in GPe→Pf trials, whereas LED-off or GPe→PPN trials showed normal lever-pressing performance. **(I–K)** Raster plots of lever-press sequences during LED-off (**I**) and LED-on trials for GPe→Pf (**J**) and GPe→PPN (**K**) in YFP control mice. The gray shaded region denotes the 5-second epoch of 40 Hz blue-light illumination. Lever-pressing performance remained unchanged during the 5-second illumination in both GPe→Pf and GPe→PPN trials, confirming that behavioral effects observed in ChR2 mice were not due to nonspecific light delivery. (**M**) Cumulative distributions showing the percentage of lever-press sequences completed over normalized sequence duration during LED-on (dark) and LED-off (light) trials in YFP-expressing mice. The gray shaded region denotes the 5-second epoch of 40 Hz blue-light illumination. Lever-pressing performance remained unchanged during the 5-second illumination in both GPe→Pf and GPe→PPN trials, confirming that behavioral effects observed in ChR2 mice were not due to nonspecific light delivery. (**N–S**) Quantification of lever pressing during stimulation (N), post-stimulation lever pressing (O), lever presses per reward (P), sequence duration (Q), reward rate (R), and inter-press interval (**S**) across stimulation conditions (LED off, LED on Pf, LED on PPN) in ChR2 and YFP groups (ChR2: n = 6; YFP: n = 8). In ChR2 mice, optogenetic stimulation significantly reduced lever pressing during stimulation (F(1.261, 6.306) = 31.57, P = 0.0009; Pf: P = 0.0049; PPN: P = 0.5482), increased post-stimulation lever pressing (F(1.278, 6.389) = 13.54, P = 0.0075; Pf: P = 0.0366; PPN: P = 0.9353), prolonged sequence duration (F(1.157, 5.784) = 12.07, P = 0.0127; Pf: P = 0.0196; PPN: P = 0.3311), decreased reward rate (F(1.847, 9.233) = 5.317, P = 0.0309; Pf: P = 0.0315; PPN: P = 0.9658), and increased inter-press interval (F(1.158, 5.788) = 11.64, P = 0.0137; Pf: P = 0.0246; PPN: P = 0.2915), with no effect on lever presses per reward (F(1.118, 5.591) = 0.5925, P = 0.4911). Effects were driven by stimulation of Pf-projecting neurons, with no effect of PPN stimulation. In YFP mice, no significant effects of optogenetic stimulation were observed across metrics (lever pressing during stimulation: F(1.952, 13.67) = 0.5236, P = 0.5996; post-stimulation: F(1.179, 8.250) = 0.3695, P = 0.5941; lever presses per reward: F(1.350, 9.453) = 0.3154, P = 0.6538; sequence duration: F(1.504, 10.53) = 0.6937, P = 0.4815; reward rate: F(1.754, 12.28) = 0.9895, P = 0.3889; inter-press interval: F(1.636, 11.45) = 0.0631, P = 0.9091). Data are mean ± SEM. ****P < 0.0001, ***P < 0.001, **P < 0.01, *P < 0.05; ns, not significant.

To directly test the functional contribution of GPe projections to the Pf and PPN during skilled forelimb behavior, we bilaterally expressed Channelrhodopsin-2 (ChR2–YFP, n = 6) or a control YFP (n = 8) in the GPe and implanted optical fibers above the GPe axonal terminals in either the Pf or PPN. Optogenetic stimulation (40 Hz, 5-second duration) was triggered immediately after the first lever press in a lever-press sequence, thereby enhancing inhibitory output from the GPe to its downstream targets. Stimulation of the GPe→Pf projection markedly impaired the execution of lever-press sequences (**Figures 7G**, **7L**, and **7N–7S**; **Video S15**). Mice expressing ChR2 exhibited little to no lever pressing during the 5-second stimulation window (**Figure 7G**), resulting in a significant reduction in lever presses during this period relative to LED-off trials (**Figures 7F**, **7G**, **7L**, and **7N**). Interestingly, lever pressing rebounded after stimulation ended, with a significant increase in presses during the 5 seconds following stimulation (**Figure 7O**), resembling rebound effects reported after optogenetic perturbation of D2-MSNs in DLS^8^. This suggests that this stimulation disrupted the execution of the sequence but not the motivation to press or the number of elements that the sequence should contain. Sequence duration was significantly prolonged during LED-on trials (**Figure 7Q**), and mice earned significantly fewer reinforcers per minute (**Figure 7R**). Consistently, the temporal structure of the action sequence was degraded, as evidenced by a significant increase in inter-press intervals (**Figure 7S**). In contrast, stimulation of GPe→PPN projections had no significant effect on any behavioral metrics measured, including total lever presses, sequence duration, reinforcer rate, or inter-press interval (**Figures 7H**, **7L**, and **7N–7S**; **Video S15**). Likewise, light delivery to either Pf or PPN in YFP-expressing control mice had no detectable behavioral effects (**Figures 7I–7S**; **Video S16**), confirming that the observed deficits were not due to nonspecific effects of light delivery. Taken together, these results demonstrate that GPe-mediated inhibition of the Pf but not the PPN disrupt the execution of forelimb sequences, showing a double dissociation between the behavioral effects of GPe→PPN and GPe→Pf projections.

## Discussion

Our findings redefine the functional role of the GPe within the basal ganglia circuitry. Traditionally, the GPe has been viewed as an intermediate node in the indirect pathway, receiving inhibitory input from D2-MSNs and sending inhibitory projections to the STN as well as to the basal ganglia output nuclei GPi and SNr. In contrast, our results reveal that the GPe is a major hub with vast and diverse projections beyond the basal ganglia, targeting a broad array of cortical, thalamic, and brainstem structures. Furthermore, our results reveal that GABAergic projection neurons in GPe can be classified into at least seven molecularly distinct cell types, of which only one completely matches the prototypical neurons of the canonical model. Single-neuron reconstructions further resolved the diverse and distinct projection patterns of GPe neurons. We show that GPe projections to PPN and Pf directly and specifically modulate locomotion and forelimb movements. Furthermore, our optogenetic manipulations targeting different stages of the basal ganglia signal flow, including input alone (D2-MSNs in the DLS), specific GPe output pathways, or combined manipulation of basal ganglia input and defined GPe output channels, provide strong evidence for the existence of direct disinhibitory circuits linking D2-MSNs to brainstem and thalamic targets. These results call for a conceptual revision of basal ganglia organization, integrating both classical intranuclear relay circuits and novel extrinsic output modules, where GPe, GPi and SNr can all be output inhibitory circuits, and where both D1 and D2-MSNs can disinhibit downstream circuits with specific behavioral functions.

### The GPe as an Output Modular Hub of the Basal Ganglia

Classical models posit that basal ganglia output arises exclusively from the SNr and GPi complex, which exert tonic inhibitory control over thalamic and brainstem motor centers^1,42^. Within this framework, the GPe is viewed as an intermediate relay in the indirect pathway, transmitting inhibition from D2-MSNs to the STN, thereby contributing to movement suppression.

Our results fundamentally revise this view by demonstrating that although some GPe neurons project to other basal ganglia nuclei, the majority extend beyond the basal ganglia to the cortex, thalamus, midbrain, and brainstem. Specific GPe subpopulations, particularly those projecting to Pf in thalamus and the PPN in brainstem, send direct long-range connections totargets outside the basal ganglia and exhibit projection-specific disinhibition. Beyond these functionally characterized examples, our whole-brain reconstructions reveal that GPe neurons also extend to a wide array of cortical, thalamic, and midbrain targets, suggesting a broader and modular architecture for pallidal output.

Functionally, D2-MSNs classically considered movement-suppressing, can instead promote locomotion by disinhibiting the GPe→PPN pathway. Optogenetic activation of D2-MSNs reliably triggered locomotor bouts, an effect abolished by concurrent activation of GP projections to PPN. This defines a D2-MSNs→GPe→PPN disinhibitory route through which D2-MSNs facilitate movement. Together, these findings challenge the rigid direct/indirect dichotomy, positioning the GPe as both an internal relay and a distributed output hub that shapes distinct behavioral programs through parallel inhibitory channels. This refined framework has important implications for understanding how pathway-specific GPe dysfunction contributes to diverse motor impairments in Parkinson’s and related basal ganglia disorders.

### Transcriptomic and Anatomical Logic of Modular GPe Organization

By integrating projection-specific single-nucleus RNA sequencing, retrograde labeling, and soma localization, we establish a comprehensive molecular and anatomical framework for the modular organization of the GPe. Single-cell and projection-defined transcriptomic analyses reveal that GPe GABAergic neurons are molecularly diverse with at least 7 clusters, only one of which corresponds entirely to the classical prototypical neurons. Distinct transcriptional clusters align with specific projection patterns and soma localization, forming discrete subpopulations with distinct projection combinations. The convergence of molecular identity, projection topology, and soma positioning suggests a genetically determined blueprint for parallel output channels within the pallidum.

GPe neurons projecting to Pf in thalamus and the PPN in brainstem form largely non-overlapping, spatially segregated populations, consistent with our retrograde tracing, 3D reconstruction, and soma registration to the Allen Brain Atlas. Single-neuron reconstructions further confirm that these neurons exhibit highly specific axonal arborization with minimal collateralization, establishing that GPe outputs are organized into parallel, functionally specialized inhibitory circuits rather than diffuse networks. This spatial segregation is complemented by projection-specific gene expression programs, indicating that molecular identity predicts both connectivity and anatomical position.

Importantly, our transcriptomic analysis reveals that classical molecular markers such as *Pvalb*, *Npas1*, *Lhx6*, and *Nkx2.1*, long used to define GPe subtypes, exhibit extensive overlap across clusters and variable expression levels, indicating that they may not serve as precise markers of GPe neuron identity. *Pvalb* expression spans a continuum from high to low and frequently co-occurs with *Lhx6* and *Nkx2.1*, indicating that these canonical markers delineate overlapping transcriptional continua rather than discrete cell types. Consequently, the traditional binary classification of “prototypic” versus “arkypallidal” neurons likely oversimplifies the molecular and functional diversity within the GPe.

Previous studies have proposed that Nkx2.1+/Npas1+ GPe neurons preferentially project to the cortex, whereas Nkx2.1+/Npas1− neurons correspond to prototypical cells and Foxp2+/Npas1+ neurons to arkypallidal populations^27^. In our dataset, however, transcriptomic clusters corresponding to Npas1+ populations exhibit more diverse projection patterns. In particular, cluster 3_GPe_Mboat1, which is enriched for markers associated with Nkx2.1+/Npas1+ neurons, displays broad projection targets rather than selective cortical innervation, whereas cortical-projecting neurons are enriched in transcriptomic clusters 4_GPe_Tac2 and 5_GPe_Pappa. One possible explanation for this difference is that prior projection mapping relied on Npas1-Cre driver lines, which label a heterogeneous population of Npas1+ neurons^17^. In contrast, our approach integrates projection-defined labeling with transcriptomic resolution, allowing finer separation of Npas1+ subpopulations with distinct projection patterns. These findings suggest that Npas1+ GPe neurons are not a uniform class but comprise multiple transcriptionally and anatomically distinct subtypes.

Our integrated single-cell analysis identifies novel, cluster-specific marker genes that more accurately define distinct GPe subtypes and predict their projection logic. Newly delineated populations include *Tac2⁺* and *Pappa⁺* neurons in ventral GPe projecting to motor and somatosensory cortex, *Meis2⁺* neurons targeting the striatum, and *Mboat1⁺*neurons with broad projections encompassing Pf, PPN, STN, and SNr. These cluster-specific markers offer a refined molecular taxonomy that bridges transcriptomic identity with projection topology and spatial organization, providing a mechanistic substrate for modular pallidal output.

Together, these results demonstrate that transcriptional identity, spatial topology, and connectivity converge to define a structured and modular organization of the GPe, in which distinct molecular modules give rise to anatomically segregated channels that collectively orchestrate diverse aspects of behavior.

### Projection- and Action-Specific Encoding in GPe Outputs

Our functional imaging and perturbation experiments reveal that the GPe is organized into projection- and action-specific modules, each tuned to distinct behavioral domains. Projection-specific calcium imaging and closed-loop optogenetic manipulations show that GPe→PPN neurons decrease their activity at locomotor onset, and that suppressing their output facilitates movement initiation through a disinhibitory mechanism. In contrast, GPe→Pf neurons exhibit minimal modulation during locomotion but display robust decreases during the initiation and execution of learned forelimb action sequences. Thus, the GPe governs locomotor and skilled forelimb behaviors through anatomically segregated inhibitory pathways that act as parallel output channels for behaviorally distinct actions.

Monosynaptic rabies tracing further reveals that these output-defined GPe populations receive inputs from distinct subsets of D2-MSNs in striatum. This input specificity indicates that the indirect pathway is not a uniform, movement-suppressing stream but a modular network of separable input–output channels tuned to different motor functions. Subsets of D2-MSNs preferentially engage the GPe→PPN circuit to modulate locomotor control, whereas others target the GPe→Pf circuit to shape the sequencing and precision of forelimb actions.

Together, these findings demonstrate a fine-grained, behavior-specific wiring logic within the D2-MSNs pathway, establishing that D2-MSNs can facilitate or suppress movement depending on the output module they engage. This modular organization provides a circuit-level mechanism for how basal ganglia networks achieve target-specific disinhibition, extending the principle of functional modularity, long recognized in striatal ensemble organization, into the pallidal and brainstem levels of motor control.

### A Revised View of D2-MSN Pathway Function

Our findings support a conceptual revision of the D2-MSN pathway. Rather than a single inhibitory stream that uniformly suppresses movement, the D2-MSN→GPe circuit comprises multiple projection-defined subcircuits that shape motor output through target-specific disinhibition. Within this modular framework, distinct subsets of D2-MSNs recruit discrete GPe output channels to control separate behavioral domains. The D2-MSN→GPe→PPN route operates as a disinhibitory loop that facilitates locomotion by transiently relieving GPe-mediated inhibition of brainstem motor centers, whereas the D2-MSN→GPe→Pf pathway regulates the precision and sequencing of skilled forelimb actions through dynamic inhibition of thalamocortical circuits. This division of labor redefines the indirect pathway as a flexible, context-dependent network capable of both facilitating and constraining movement, depending on which GPe module is engaged. The discovery of parallel disinhibitory subcircuits thus reconciles the long-standing paradox that D2-MSNs can both promote and suppress movement, providing a mechanistic framework for adaptive action selection^6–8,10,11^.

Finally, this modular redefinition of the GPe shifts its conceptual role within the basal ganglia from an intermediate relay to a flexible output control hub where striatal, pallidal, and subcortical computations converge, and information is then broadcasted out through specific channels. Future studies should dissect the synaptic and neuromodulatory mechanisms governing plasticity within D2-MSNs→GPe pathways and determine how dopaminergic tone, behavioral context, and learning dynamically rebalance activity across distinct GPe output modules in health, disease, and recovery.

### Implications for basal ganglia disorders

These insights also have direct relevance to the pathophysiology of Parkinson’s disease. Differential vulnerability or maladaptive plasticity across GPe output modules could underlie the heterogeneity of motor symptoms. Hypofunction of the GPe→PPN channel may contribute to gait freezing and akinesia, whereas disruption of the GPe→Pf pathway could impair fine motor coordination and action sequencing. Importantly, our transcriptomic analyses further revealed that these projection-defined pathways correspond to molecularly distinct GPe neuron populations, linking specific projection targets to defined transcriptional identities. More broadly, the identification of molecularly and projection-defined GPe modules provides a blueprint for circuit-level therapeutic strategies, including targeted neuromodulation, chemogenetic, or optogenetic interventions designed to selectively restore function within specific disinhibitory channels rather than globally modulate basal ganglia activity.

## Supplemental Figure Legends

**Figure S1. Brain-wide distribution of GPe projections quantified across coronal levels, related to Figure 1.**

(**A**) Coronal sections registered to the Allen Common Coordinate Framework (CCFv3) showing the fractional distribution of total GPe axonal output across brain regions at representative rostrocaudal levels (CCF 0160, 0260, 0300, 0340, and 0405). Color intensity indicates the percentage of total GPe projection signal within each region, normalized to the sum of all detected GPe projections across the brain.

(**B**) Corresponding coronal sections showing GPe projection density, defined as the volume of GPe axonal signal within each brain region normalized by the volume of that region. Color intensity reflects relative projection density, highlighting regions with particularly dense GPe innervation independent of regional size.

**Figure S2. Transcriptomic Heterogeneity and Data Integration of GPe Neurons, related to Figure 2**

(**A**) Complete data processing and integration workflow. Single-cell RNA-seq (10x snRNA-seq), single-nucleus RNA-seq + ATAC-seq (10x snRNA-seq Multiome), and SMART-Seq v4 Retro-seq datasets were independently quality-controlled, filtered, and annotated prior to iterative integration, benchmarking, clustering, and removal of low-confidence or non-neuronal populations.

(**B**) Broad cell-type annotation of all sequenced cells and nuclei following initial quality control. UMAP visualization showing major cellular classes identified across the combined dissectate dataset after only performing QC filtering, including cholinergic, dopaminergic, GABAergic, glutamatergic, immature neuronal, and non-neuronal populations, based on mapping to the AIT21 taxonomy using MapMyCells (CCN20230722)^48^. Non-GABAergic and non-neuronal populations were excluded from downstream analyses of GPe neuronal organization.

(**C**) UMAP visualization showing cell-class annotation of all sequenced cells and nuclei based on mapping to the AIT21 atlas. Major classes include GABAergic neuronal populations from multiple brain regions (CNU-HYa GABA, CNU-LGE GABA, CNU-MGE GABA, CTX-CGE GABA, CTX-MGE GABA, HY GABA, MB GABA, MY GABA, OB-IMN GABA, P GABA), glutamatergic populations (CNU-HYa Glut, HY Glut, IT-ET Glut, MB Glut, MY Glut, TH Glut), striatal projection neurons (SPNs), as well as non-neuronal populations including astrocyte/ependymal cells, oligodendrocyte precursor cells (OPC-Oligo), immune cells, and vascular cells. This classification was used to exclude non-GPe and non-GABAergic populations prior to downstream analyses.

(**D**) UMAP visualization colored by sequencing platform, demonstrating robust mixing of scRNA-seq, snRNA-seq Multiome, and SSv4 retro-seq data following integration.

(**E**) UMAP highlighting included versus excluded clusters after quality control, doublet removal, spatial filtering, and exclusion of low-confidence annotations, yielding a refined population enriched for bona fide GPe neurons.

(**F**) Violin plots showing expression distributions of canonical GPe markers (*Pvalb*, *Lhx6*, *Npas1*, *Foxp2*) across the seven integrated GPe GABAergic transcriptomic clusters, illustrating graded and overlapping expression patterns rather than strict binary segregation. These plots reveal substantial overlap among classical molecular markers across GPe neuron types, including prominent co-expression of *Pvalb* and *Lhx6* in clusters 1 and 3, *Pvalb* and *Npas1* in clusters 3 and 6, and *Pvalb* and *Foxp2* in cluster 6, highlighting the continuous and non-mutually exclusive nature of these classical markers across GPe neuron populations.

(**G**) UMAP feature plots showing the expression of *Pvalb*, *Npas1*, and *Foxp2* overlaid on the integrated GPe transcriptomic clusters. These plots reveal substantial overlap among these three classical molecular markers across GPe neuron types, including prominent co-expression of *Pvalb* and *Npas1* in clusters 3 and 6, and *Pvalb* and *Foxp2* in cluster 6, highlighting the continuous and non-mutually exclusive nature of these classical markers across GPe neuron populations.

(**H**) Comparison of GPe neurons identified in the present study versus Saunders et al.^25^, shown in a shared UMAP space. Cells unique to each dataset and overlapping populations are indicated, demonstrating both correspondence and expansion of previously defined GPe cell types.

(**I**) UMAP visualization colored by AIT21 transcriptomic cell types, showing correspondence between AIT21 annotations and the integrated GPe clusters defined in this study.

(**J**) UMAP visualization colored by the seven integrated GPe transcriptomic clusters identified here, representing de novo clustering after integration of all datasets.

(**K**) UMAP visualization colored by Saunders et al. subclusters^25^, allowing direct comparison between prior classifications and the present integrated clustering framework.

(**L**) Sankey diagram linking AIT21 cell types, integrated GPe clusters from the present study, and Saunders et al. subclusters^25^. This analysis highlights both conserved relationships and novel subdivisions, including previously unrecognized GPe transcriptomic populations.

**Figure S3. Retrograde labeling strategy for projection-defined GPe subpopulations, related to Figure 3**

(**A**) Schematic illustrating the retrograde AAV labeling strategy used to define projection-specific GPe subpopulations. AAVretro viruses were injected into distinct downstream targets of the GPe, including Pf, PPN, STN, DLS, MO+SS, and SNr, enabling labeling of corresponding GPe projection populations.

(**B**) Representative coronal sections showing retrogradely labeled GPe neurons following injections into individual target regions. Fluorescent signals indicate projection-defined GPe populations corresponding to MO+SS, DLS, Pf, PPN, STN, and SNr. DAPI counterstain marks nuclei.

(**C**) Schematic of the dual-color retrograde labeling strategy used for fluorescence-activated cell sorting (FACS) prior to single-nucleus sequencing. AAVretro-H2B-mCherry and AAVretro-H2B-eGFP were injected into Pf and PPN, respectively, allowing nuclear-localized labeling of GPe^Pf^ and GPe^PPN^ neurons.

(**D**) Coronal brain section containing the GPe showing parvalbumin (Pvalb) immunostaining used to assess marker expression in projection-defined neurons.

(**E**, **F**) High-magnification views of the boxed region in (**D**) showing H2B-eGFP (green), H2B-mCherry (red), and Pvalb immunostaining. Arrowheads indicate representative single-labeled eGFP⁺ (green), mCherry⁺ (red), and dual-labeled neurons (yellow). Note that mCherry⁺ GPe^Pf^ neurons co-express Pv.

**Figure S4. Spatial prediction of GPe neuron soma locations from transcriptomic identity, related to Figure 3**

(**A**) Schematic illustrating the strategy for predicting soma locations from gene expression. Spatial transcriptomic data from MERFISH provide observed soma coordinates and gene expression profiles, which are used to train predictive models. The trained models are then applied to retro-seq transcriptomic data to infer soma locations based on shared gene features. (**B**–**D**) Performance of three predictive models trained on MERFISH data to infer soma location from gene expression. Scatter plots show predicted versus observed spatial coordinates for the linear regression model (**B**), random forest model (**C**), and XGBoost model

(**D**). Each point represents a single cell; predicted and observed positions are shown in different colors.

(**B**) Ground-truth soma locations from MERFISH data shown in three-dimensional coordinate space, serving as the reference distribution for model evaluation.

(**C**) Observed soma locations of retrogradely labeled GPe neurons following rAAVretro injections into different target regions (DLS, MO+SS, Pf, PPN, DLS/STN, STN, SNr), shown as density maps. Color scale indicates the absolute number of labeled cells; the maximum value for each panel is indicated on the panel.

(**D**) Quantitative comparison of prediction accuracy across models and spatial dimensions. Mean absolute error is shown for each model (linear, random forest, XGBoost) along the mediolateral (ML), dorsoventral (DV), and anteroposterior (AP) axes.

(**E** and **F**) Observed soma distributions from rAAVretro tracing for the corresponding projection groups, plotted as individual soma locations within the GPe.

(**G**) Predicted soma locations for retro-seq GPe neurons based on transcriptomic identity using the trained MERFISH-derived model, shown as density maps for each projection group.

(**H**) Sankey diagram illustrating the correspondence between projection-defined GPe populations and AIT21 cell-type clusters based on inferred soma locations.

(**I**) Predicted soma location density maps for MERFISH-defined GPe cell types corresponding to retro-seq projection groups.

(**J**) Observed soma location density maps for the same MERFISH-defined AIT21 cell types, shown for comparison with predicted distributions.

**Figure S5. Clustering of single-neuron GPe projectomes, related to Figure 4.**

(**A**) Soma localization of all 20 reconstructed GPe neurons plotted separately for the left and right hemispheres in top view. Neurons are color-coded by projection-defined group. Dot size reflects the dorso-ventral coordinate (Y coordinate) of each soma.

(**B**) Soma localization of the same 20 neurons plotted separately for the left and right hemispheres in coronal view. Neurons are color-coded by projection-defined group. Dot size reflects the rostro-caudal (X coordinate) of each soma. The dashed outline denotes the GPe boundary.

(**C**) Single-neuron axonal projection matrices derived from whole-brain reconstructions of individual GPe neurons were used for unsupervised clustering based on similarity of axonal projection patterns. Each column represents one reconstructed GPe neuron, and each row represents an anatomical target region registered to the Allen Common Coordinate Framework. Hierarchical clustering is shown as a dendrogram, revealing four distinct projection-defined groups. The color bar indicates cluster assignment for each neuron. Bar plots summarize total reconstructed axonal length (neuron_length) for each neuron, and scatter plots indicate the mediolateral soma position (y_position) within the GPe. Together, these analyses reveal four distinct projectome groups, ranging from highly target-specific neurons to neurons with broader, multi-target collateralization patterns, and highlight relationships between projection pattern, axonal extent, and soma location.

**Figure S6. Cell-type– and projection-specific monosynaptic rabies tracing of GPe^Pf^ and GPe^PPN^ neurons, related to Figure 4.**

(**A**–**D**) Representative coronal section through the Pf showing EYFP expression driven by Flpo-dependent reporter activation in rabies-labeled axons originating from GPe^Pf^ neurons (green), mCherry signal from the rabies virus (red), HA immunostaining for TVA/G expression (magenta), and DAPI (blue).

(**E**–**L**) High-contrast inverted images illustrating the distribution of labeled axons across major brain regions following monosynaptic rabies tracing from GPe^Pf^ neurons. Robust labeling is observed in the GPe and the intended downstream target region (Pf), with minimal or no labeling in other brain areas. Regions shown include M1, DLS, GPe, GPi, STN, Pf, SNr, and PPN, as indicated.

(**M**–**P**) Representative coronal section through the PPN showing EYFP expression driven by Flpo-dependent reporter activation in rabies-labeled axons originating from GPe^PPN^ neurons (green), mCherry signal from the rabies virus (red), HA immunostaining for TVA/G expression (magenta), and DAPI (blue).

(**Q**–**X**) High-contrast inverted images illustrating the distribution of labeled axons across major brain regions following monosynaptic rabies tracing from GPe^PPN^ neurons. Robust labeling is observed in the GPe and the intended downstream target region (PPN), with minimal or no labeling in other brain areas. Regions shown include M1, DLS, GPe, GPi, STN, Pf, SNr, and PPN, as indicated.

**Figure S7. Projection-specific monosynaptic rabies tracing reveals distinct striatal MSN inputs to GPe^Pf^and GPe^PPN^neurons, related to Figure 6.**

(**A–B**) Cell-type- and projection-specific monosynaptic rabies tracing strategy to identify striatal MSN inputs to GPe neurons projecting to the PPN and Pf. AAVretro-EF1α-Flpo was injected into the PPN or Pf to label GPe^PPN^ or GPe^Pf^ neurons, respectively. In the GPe, Flpo-dependent helper viruses (AAV1.EF1a.FlpX.H2B.3XHA.2A.N2C(G) and AAVDJ.hSyn.FlpX.TVA950) enabled selective expression of TVA and rabies glycoprotein. Three weeks later, EnvA-N2cΔG-GFP was injected into the GPe to label monosynaptic striatal inputs. Experiments targeting PPN and Pf were performed in separate animals; identical viral strategies were used, and rabies-labeled neurons are color-coded for visualization (green, PPN; red, Pf).

(**C**) 3D reconstruction of rabies starter neurons within the GPe, showing the spatial distribution of GPe^Pf^ (red) and GPe^PPN^ (green) neurons.

(**D**) Quantification of labeled cell counts for projection-specific monosynaptic rabies tracing. Bar plots show the number of starter GPe neurons (GPe^PPN^, green; GPe^Pf^, red) and monosynaptically labeled striatal MSNs providing input to each GPe subpopulation (MSNs-GPe^Pf^, red; MSNs-GPe^PPN^, green). No significant differences were observed between PPN- and Pf-targeted groups for either starter cells (P = 0.8636) or input cells (P = 0.7240; two-tailed unpaired t tests; n = 3 mice per group).

(**E**) Top view of representative 3D reconstructions of striatal MSNs providing monosynaptic input to GPe subpopulations (MSNs-GPe^Pf^, red; MSNs-GPe^PPN^, green). For visualization, a random subset of labeled neurons is shown (1,400 per group), illustrating the spatial segregation of input populations within the striatum.

(**F**–**H**) Zoomed-in views of striatal MSN input populations. (**F**) MSNs-GPe^Pf^ neurons (red).

(**G**) MSNs-GPe^PPN^ neurons (green). (**H**) Merged view showing the relative spatial segregation of MSNs-GPe^Pf^ and MSNs-GPe^PPN^ populations.

**Figure S8 Closed-loop excitation of GPe→PPN but not GPe→Pf output suppresses D2-MSN–driven locomotion, related to Figure 6.**

(**A**) Schematic of the closed-loop optogenetic excitation paradigm. Online behavioral clustering was used to detect immobility, which triggered optogenetic excitation of GPe→PPN or GPe→Pf axons (40 Hz, 5 s; inter-trial interval 30 s).

(**B**) Example acceleration traces aligned to immobility-triggered excitation, comparing LED-off and LED-on conditions. Shaded region indicates the 5 s excitation window.

(**C**) Quantification of the total body acceleration during the excitation window across experimental groups. No significant differences were observed between conditions or groups.

(**D**) Quantification of forward locomotion probability during immobility-triggered excitation. Excitation of GPe→PPN but not GPe→Pf significantly reduces the probability of forward locomotion following D2-MSN activation. Control groups show no significant change.

(**E**) Schematic of the closed-loop optogenetic stimulation paradigm. Online behavioral clustering was used to detect locomotion onset, which triggered concurrent stimulation of D2-MSNs (14 Hz, 5 s) with or without GPe→PPN or GPe→Pf excitation.

(**F**) Example acceleration traces aligned to locomotion-triggered stimulation, comparing LED-off and LED-on. Shaded region denotes the stimulation window.

(**G**) Quantification of the total body acceleration during locomotion-triggered stimulation across experimental groups. No significant differences were observed between conditions or groups.

(**H**) Quantification of forward locomotion probability during locomotion-triggered stimulation. Activation of D2-MSNs significantly increases the probability of forward locomotion, an effect that is selectively suppressed by co-activation of GPe→PPN but not GPe→Pf terminals. No effects were observed in YFP control mice.

## Supplementary videos

**Video S1. Brain-wide distribution of GPe axonal projections across major brain regions, related to Figure 1**

Sequential coronal sections spanning the entire anterior–posterior extent of the mouse brain showing the distribution of axonal projections from GPe neurons. One coronal section is displayed at a time, progressing from anterior to posterior, with projection signal registered to the Allen Brain Atlas. Color intensity represents the fraction of total GPe axonal output within each brain region, illustrating widespread GPe projections beyond classical basal ganglia targets, including cortex, thalamus, midbrain, and brainstem regions.

**Video S2. Brain-wide density map of GPe axonal projections reveals extensive extra–basal ganglia outputs, related to Figure 1**

Sequential coronal sections spanning the full anterior–posterior extent of the mouse brain showing the density of axonal projections from dorsolateral striatum–innervated GPe neurons. One coronal section is displayed at a time, progressing from anterior to posterior, with axonal density calculated as projection volume normalized to the volume of each brain region and registered to the Allen Brain Atlas. Color intensity reflects projection density within each region, highlighting dense GPe innervation of cortical, thalamic, midbrain, and brainstem structures in addition to classical basal ganglia targets.

**Video S3. Single-neuron reconstructions of GPe neurons, related to Figure 4**

3D visualizations of individually reconstructed GPe neurons showing whole-brain axonal arbors from single cells. Each neuron is rendered separately and registered to the Allen Brain Atlas, illustrating the diversity of GPe output architectures, ranging from highly target-specific projections to neurons with extensive multi-target collateralization across cortical, thalamic, basal ganglia, and brainstem regions. Somata are located within the GPe, and axons are traced continuously throughout the brain volume to reveal projection specificity at the single-neuron level.

**Video S4. Optogenetic stimulation of GPe projections to the PPN, but not the Pf, impairs locomotion, related to Figure 5**

This video shows representative open-field behavior during optogenetic activation of GPe axonal projections to downstream targets. Three video segments are stitched in the following order: (1) activation of GPe→PPN terminals in ChR2-expressing mice, (2) activation of GPe→Pf terminals in ChR2-expressing mice, and (3) an LED-off control trial from the same behavioral context. Blue-light stimulation of GPe→PPN projections markedly suppresses ongoing locomotion, whereas stimulation of GPe→Pf projections does not produce a comparable effect. No locomotor impairment is observed during LED-off trials. This video illustrates the pathway-specific role of GPe→PPN projections in regulating locomotor behavior.

**Video S5. Light delivery to GPe projections to the PPN and Pf in YFP control mice does not affect locomotion, related to Figure 5**

This video presents representative open-field behavior from YFP control mice during light delivery to GPe axonal projections. Three video segments are stitched in the following order: (1) light delivery to GPe→PPN terminals in YFP-expressing mice, (2) light delivery to GPe→Pf terminals in YFP-expressing mice, and (3) an LED-off control trial. In contrast to ChR2-expressing animals, light delivery to either GPe→PPN or GPe→Pf projections in YFP controls does not alter locomotor behavior relative to LED-off trials, confirming that the effects observed in Figure 5 are not due to nonspecific effects of light delivery.

**Video S6. Optogenetic inhibition of GPe→PPN projections, but not GPe→Pf projections, induces locomotion, related to Figure 5**

This video shows representative behavioral trials from mice expressing NpHR in GPe neurons, with optical fibers positioned over axonal terminals in the PPN or Pf. Three video segments are stitched in the following order: (1) optogenetic inhibition of GPe→PPN projections during LED-on trials, (2) optogenetic inhibition of GPe→Pf projections during LED-on trials, and (3) an LED-off control trial. Inhibition of GPe→PPN terminals reliably induces locomotion relative to LED-off trials, whereas inhibition of GPe→Pf terminals does not elicit locomotor behavior, demonstrating pathway-specific disinhibition of locomotion through the GPe→PPN circuit.

**Video S7. Light delivery to GPe→PPN and GPe→Pf projections in YFP control mice does not induce locomotion, related to Figure 5**

This video presents representative behavioral trials from YFP control mice with optical fibers positioned over GPe axonal terminals in the PPN or Pf. Three video segments are stitched in the following order: (1) light delivery to GPe→PPN projections in YFP-expressing mice, (2) light delivery to GPe→Pf projections in YFP-expressing mice, and (3) an LED-off control trial. In all conditions, light delivery fails to induce locomotion relative to LED-off trials, confirming that the locomotor effects observed in experimental animals require opsin-mediated manipulation and are not due to nonspecific effects of light delivery.

**Video S8. Closed-loop optogenetic activation of D2-MSNs triggered during immobility initiates locomotion, related to Figure 6**

This video shows representative behavioral trials from mice expressing ChR2 in D2-MSNs in the dorsolateral striatum. Two video segments are stitched in sequence: (1) an LED-on trial in which closed-loop optogenetic activation of D2-MSNs is triggered upon detection of an immobility cluster, and (2) a corresponding LED-off control trial. Optogenetic activation of D2-MSNs during immobility reliably initiates locomotion, whereas no locomotor initiation is observed during LED-off trials.

**Video S9. Immobility-triggered light delivery to D2-MSNs in YFP control mice does not induce locomotion, related to Figure 6**

This supplementary video is compiled from two trials recorded from YFP control mice. In the first trial, top-down behavioural tracking is shown during a 14 Hz blue-light stimulus delivered whenever immobility is detected; the mouse expresses the YFP control construct in D2-MSNs and therefore does not respond to the light, remaining immobile throughout the stimulation window. In the second trial, no light is delivered (LED-off), and the mouse likewise remains stationary. These trials collectively demonstrate that light delivery alone does not initiate locomotion in YFP control mice, supporting the specificity of the locomotor responses observed with optogenetic activation of D2-MSNs.

**Video S10. Closed-loop, locomotion-triggered optogenetic inhibition of D2-MSNs suppresses ongoing locomotion, related to Figure 6**

This video comprises two trials recorded from mice expressing the inhibitory opsin Jaws in dorsolateral striatal D2-MSNs. In the first trial, a continuous red-light stimulus is delivered when the animal’s movement is detected, implementing a closed-loop inhibition paradigm; this real-time silencing of D2-MSNs causes the mouse’s forward locomotion to slow and stop during the light-on period. In the second trial, no light is delivered during locomotion (LED-off control), and the animal maintains its locomotor behavior without interruption. Together, these trials demonstrate that closed-loop, locomotion-triggered inhibition of D2-MSNs is sufficient to suppress ongoing locomotion, while light-off controls confirm that the effect requires optogenetic silencing.

**Video S11. Locomotion-triggered light delivery to D2-MSNs in tdTomato control mice does not alter ongoing locomotion, related to Figure 6**

This supplementary video consists of two trials recorded from tdTomato control mice. In the first trial, a closed-loop setup detects when the mouse is already moving and triggers continuous red-light illumination of striatal D2-MSNs. Since the neurons only express tdTomato (and not the light-sensitive inhibitor), the optical stimulation does not affect the mouse’s behavior, and ongoing locomotion continues uninterrupted. In the second trial (LED-off control), the locomotor activity proceeds without any light delivery, showing comparable behavior. Together, these trials demonstrate that locomotion-triggered light delivery does not alter ongoing movement in tdTomato control mice, confirming that any suppression seen with optogenetic inhibition in experimental mice is due to the specific optogenetic manipulation rather than the light itself.

**Video S12. Immobility-triggered activation of D2-MSNs induces locomotion that is selectively blocked by concurrent activation of GPe→PPN, but not GPe→Pf, projections, related to Figure 6**

This supplementary video consists of four immobility-triggered optogenetic stimulation trials illustrating the interplay between striatal D2-MSN activation and GPe efferent modulation in the regulation of locomotion. In the first trial (D2-MSNs-ChR2 only), real-time detection of immobility triggers 14 Hz blue-light pulses delivered to ChR2-expressing D2-MSNs in the dorsolateral striatum; the mouse rapidly transitions from immobility to robust locomotor activity, demonstrating that selective activation of D2-MSNs is sufficient to initiate movement. In the second trial (D2-MSNs-ChR2 + GPe-PPN-ChR2), the same D2-MSN stimulation is paired with simultaneous blue-light activation of GPe→PPN terminals; despite the continued D2-MSN stimulation, the mouse remains immobile, indicating that concurrent activation of the GPe→PPN projection blocks the locomotion-initiating effect of D2-MSN activation. In the third trial (D2-MSNs-ChR2 + GPe-Pf-ChR2), immobility-triggered activation of D2-MSNs is paired with co-activation of GPe→Pf terminals; the mouse still initiates locomotion, showing that GPe→Pf activation does not impede the pro-locomotor effect of D2-MSN stimulation. The fourth trial is an LED-off baseline (no light delivery to any structure), in which the mouse remains immobile throughout the detection window. Together, the four trials demonstrate that immobility-triggered D2-MSN activation reliably initiates locomotion and that this effect is specifically suppressed by co-activation of GPe→PPN, but not GPe→Pf, projections.

**Video S13. D2-MSN–evoked locomotion is unaffected by light delivery to GPe→PPN or GPe→Pf terminals in YFP control mice, related to Figure 6**

This compiled video consists of four trials recorded from mice in which striatal D2-MSNs express ChR2 to drive locomotion, whereas GPe terminals express only YFP. Trial 1 shows locomotion evoked by optogenetic activation of D2-MSNs alone, illustrating robust movement initiation. Trial 2 shows simultaneous light delivery to YFP-labeled GPe→PPN projections during D2-MSN stimulation; locomotion remains unaffected. Trial 3 shows concurrent light delivery to YFP-labeled GPe→Pf projections during D2-MSN activation, again with no change in locomotion. Trial 4 is an LED-off baseline in which D2-MSN activation alone initiates locomotion. Together, these trials demonstrate that light stimulation of YFP-tagged GPe terminals does not alter D2-MSN-evoked locomotion, confirming that the modulatory effects observed in functional GPe projections are specific to optogenetically active channels rather than due to light delivery itself.

**Video S14. Immobility-triggered light delivery to D2-MSNs and GPe projections in YFP-only control mice does not induce locomotion, related to Figure 6**

This supplementary movie assembles four trials in which immobility-triggered optogenetic stimulation was applied in animals expressing only YFP. In the first trial, the YFP-expressing D2-MSNs received 14 Hz blue-light pulses during an immobile period; the mouse remained immobile throughout the stimulation window. In the second trial, both D2-MSNs and GPe→PPN projections expressed YFP and were stimulated; the animal again failed to initiate movement. The third trial delivered stimulation to D2-MSNs and GPe→Pf projections in another YFP-only mouse and likewise did not induce locomotion. In the final LED-off baseline trial, no light was delivered during immobility, and the animal stayed stationary. Collectively, these trials illustrate that simultaneous stimulation of D2-MSNs and GPe projections does not trigger locomotion in YFP-only control mice, confirming that light delivery alone is insufficient to evoke the behaviors observed when opsins are present.

**Video S15. Optogenetic activation of GPe→Pf, but not GPe→PPN, disrupts execution of skilled forelimb lever-press sequences, related to Figure 7F**

This supplementary video comprises three trials, demonstrating how stimulation of distinct GPe projections affects a skilled lever-press task. In the first trial, a mouse expressing ChR2 in GPe terminals receives light pulses targeting the GPe→PPN pathway while performing a lever-press sequence. Despite the optogenetic activation, the animal completes the lever presses without interruption. In the second trial, the same optogenetic stimulation is delivered to the GPe→Pf pathway. Here, the mouse begins pressing the lever but fails to complete the sequence; the task is disrupted shortly after light onset. The final trial is a control with no light delivery (LED-off condition); the animal successfully executes the full lever-press sequence. Collectively, these clips illustrate that activating GPe→Pf projections specifically disrupts ongoing lever-press behavior, whereas activating GPe→PPN projections does not alter task performance.

**Video S16. Light delivery to GPe projections to Pf or PPN in YFP control mice does not affect execution of skilled forelimb lever-press sequences, related to Figure 7F**

This supplementary video compiles three trials recorded from YFP control mice performing a trained forelimb lever-press task. In Trial 1, the mouse receives continuous blue-light stimulation targeting GPe→PPN terminals while executing a lever-press sequence. Because GPe neurons express only YFP, the photostimulation produces no optogenetic effect and lever pressing proceeds normally. In Trial 2, the same photostimulation is applied to GPe→Pf terminals; again, the animal’s lever-pressing behavior is unaffected, confirming that light delivery alone does not disrupt task performance. Trial 3 presents an LED-off condition under otherwise identical parameters, serving as a baseline. Together, these trials demonstrate that optogenetic activation of GPe projections to PPN or Pf does not alter execution of skilled forelimb lever-press sequences in YFP control mice, underscoring that the behavioral effects observed with ChR2 expression are specific to optogenetic activation of GPe neurons.

## Supplementary Information

Supplemental Table 1: summarizes the RNA-seq datasets and analysis workflow, including platform-specific cell counts, projection percentages, cluster inclusion or exclusion, Louvain clustering results, differentially expressed markers, cluster renaming, mapping percentages, and 10x sequencing metrics.

Supplemental Table 2: provides metadata for the SWC files used in the single-cell reconstruction analysis of GPe neurons. It includes file identifiers and paths, reconstruction type, registered and raw soma coordinates, hemisphere, manual brain region annotation, distance to the GPe border, Cre line information, notes on reconstruction quality, and the assigned GPe projection subtype for each reconstructed neuron.

Supplementary Videos: Provide representative visualizations of the key anatomical and behavioral findings, including brain-wide GPe projection maps, single-neuron reconstructions, projection-specific optogenetic manipulations of locomotion and skilled forelimb behavior, and closed-loop experiments demonstrating that D2-MSNs promote locomotion through a projection-specific GPe→PPN disinhibitory pathway.

## Resource availability

### Lead contact

Lead contact Further information and requests for resources and reagents should be directed to and will be fulfilled by the lead contact, Rui M. Costa.

## Materials availability

This study did not generate new unique reagents.

## Data and code availability

Single-nucleus sequencing transcriptomic will be deposited at NCBI GEO. This paper used existing python, R and MATLAB code packages and the original code for snRNA sequencing analysis can be found here: https://github.com/AllenInstitute/gpe and https://github.com/AllenInstitute/GPe_projection. Any additional information required to reanalyze the data reported in this paper is available from the lead contact upon request.

## Acknowledgments

We thank Gabi Martins and Drew Baughman for excellent laboratory management and administrative support. We thank Ana Vaz and Mariana Lopes Correia for mouse colony management and Helio Rodrigues for assistance with behavioral box development and implementation. We thank members of the Costa laboratory for thoughtful discussions. We thank David Ng and Katie Balaguer for producing AAV and rabies viruses. We thank Ira Shieren and the Zuckerman Institute Flow Cytometry Platform for assistance with FACS. We thank Humberto Ibarra for assistance with imaging and analysis. We thank Lin Tian for providing AAV1-hSyn-axon-targeted GCaMP6f. Imaging was performed with support from the Zuckerman Institute Cellular Imaging Platform. This work was supported by A Parkinson’s Foundation Summer Student Fellowship (696639) to A.C.; a Brain & Behavior Research Foundation Young Investigator Award (PG010635) and NIH Pathway to Independence Awards (K99 phase; K99NS119788 and R00 phase; R00NS119788) to Z.G.; and by NIH funding (5U19NS104649) and Aligning Science Across Parkinson’s (ASAP-020551) through the Michael J. Fox Foundation for Parkinson’s Research (MJFF) to R.M.C.

## AUTHOR CONTRIBUTIONS

Conceptualization of study: Z.G. and R.M.C., with the contribution of Z.R.L and B.T. on transcriptomic and single neuron reconstruction; Surgery, brain-wide tracing, rabies tracing: Z.G.; Histology and imaging analysis: Z.G., L.N, S.R.; BrainJ analysis and visualization: Z.G., L.H., and D.P., L.N., S.R., M.S.T., A.K.; Single-cell and single-nucleus sequencing conceptual design and analyses: Z.G., Z.R.L. A.M. R.M.C. and B.T.; Dissection, nuclear isolation and flow cytometry: Z.G., L.N., A.M.; 10x and Retro-SMART sequencing: E.T., D.B., J.G., C.R., M.T., T.C., A.T., A.B.C., K.F., K.A.S.; Sequencing analysis and visualization: Z.R.L., A.M., E.T., R.T.; Genomics to spatial positioning modeling; Z.R.L., Single cell reconstruction: S.Y., M.M., A.M., S.S., H.Z.; Single cell reconstruction analysis: Z.R.L. and M.M.; closed-loop optogenetic system design: Z.G., J.C.Y.T., and V.P.; closed-loop optogenetic experiment and analysis: Z.G., G.A.A., and A.C.; Calcium imaging experiment and analysis: Z.G. and D.P.; Lever-pressing experiment and analysis: Z.G. and M.V.; Manuscript – original draft: Z.G. and R.M.C. Original figures and manuscript editing: Z.G. and Z.R.L., B.T., and R.M.C.; Supervision: Z.G., Z.R.L. B.T., and R.M.C.; Funding acquisition: Z.G., B.T., H.Z. and R.M.C.

## Declaration of interests

The authors declare no competing interests.

STAR★Methods

Key resource table

## Experimental model and study participant details

### Experimental animals

All experiments and procedures were performed according to National Institutes of Health (NIH) guidelines and approved by the Institutional Animal Care and Use Committee of Columbia University. Adult mice of both sexes, aged 2–6 months, were used for all experiments, including slice electrophysiology. The strains used were: C57BL/6J (Jackson Laboratories, 006660); vGat-Cre (Jackson Laboratories, 016962); vGlut2-Cre (Jackson Laboratories, 016963); A2a-Cre, B6.FVB(Cg)-Tg(Adora2acre), (MMRRC, #036158-UCD); Animals were heterozygous for the transgene and generated from a cross between heterozygote Cre male mice with C57/BL6J female mice. Mice used for behavioral experiments were individually housed, and all mice were kept under a 12-h light–dark cycle.

## Method details

### Stereotaxic viral injections and fiber implants

All surgeries were performed under sterile conditions and isoflurane (1–3%) was used to anesthetize mice. Throughout each surgery, mouse body temperature was maintained at 37°C using an animal temperature controller (ATC2000, World Precision Instruments). Analgesia in the form of subcutaneous injection of carprofen (5 mg per kg body weight) or buprenorphine XR (0.5–1 mg per kg body weight) was administered on the day of the surgery, along with bupivacaine (2 mg per kg body weight). Mice were placed in a stereotaxic holder (Kopf) and a midline incision was made to expose the skull. A craniotomy was made over the injection site with a drill (Osada, Los Angeles, CA, USA). All injections were performed using a Nanoject III Injector (Drummond Scientific, Broomall, PA, USA) at a rate of 4.6 nL every 5 s. Pulled glass injection pipettes were left in place for 5 min after injection to allow for virus absorption, and incisions were closed with Vetbond tissue adhesive (3M, Maplewood, MN, USA). Mice were subsequently allowed to recover from the anesthesia in their homecage on a heating pad.

To label GABAergic GPe neurons receiving input from the DLS, we used an intersectional viral labeling strategy. AAV1-hSyn-Flpo (Zuckerman Institute Molecular Tools Core Facility, ZI Virology; RID:SCR_026201, 1.60 × 10¹³ GC/mL) was injected into the DLS of vGat-Cre mice (AP +0.50 mm, ML −2.50 mm, DV −2.20 mm, 500 nL). In the same animals, AAV-Flex-FRT-eGFP (ZI Virology; 2.18 × 10¹³ GC/mL) was injected into the GPe (AP −0.34 mm, ML −1.90 mm, DV −3.50 mm, 240 nL). Related to Figure 1 and Figure S1.

To retrogradely label projection-defined GPe subpopulations and quantify their spatial distribution, we injected AAVretro vectors expressing nuclear-localized fluorescent reporters or Cre recombinase into three distinct GPe output targets within each mouse. Specifically, we used scAAVretro-hSyn-H2B-mCherry (Salk Institute, 1.21 × 10¹³ GC/mL), scAAVretro-hSyn1-H2B-eGFP (ZI Virology, 1.10 × 10¹³ GC/mL), and AAVretro-hSyn1-Cre (Addgene #105540, 1.17 × 10¹³ GC/mL). Ten mice were used across the following target combinations: Pf (H2B-mCherry) / PPN (H2B-eGFP) / STN (Cre), n = 4; DLS (H2B-mCherry) / MO-SS (H2B-eGFP) / STN (Cre), n = 4; and MO-SS (H2B-mCherry) / PPN (H2B-eGFP) / SNr (Cre), n = 2. Injection coordinates and volumes were as follows: Pf: AP −2.30 mm, ML −0.69 mm, DV −3.125 mm, 150 nL; PPN: AP −4.36 mm, ML −1.25 mm, DV −3.25 mm, 100 nL; STN: AP −2.06 mm, ML −1.50 mm, DV −4.40 mm, 50 nL; DLS: AP +0.50 mm, ML −2.38 mm, DV −2.13 mm, 100 nL; MO-SS: AP +0.25 mm, ML −1.25 mm, DV −0.75 mm, 100 nL; and SNr: AP −3.28 mm, ML −1.35 mm, DV −4.38 mm, 150 nL. Related to Figure 3 and supplemental figures 3 and 4.

To retrogradely label projection-defined GPe subpopulations for single-nucleus sequencing, we bilaterally injected AAVretro-hSyn-H2B-mCherry and AAVretro-hSyn-H2B-eGFP into two distinct GPe output targets per mouse. In total, 24 mice were injected across three retrograde target-pair groups: Pf (H2B-mCherry) / PPN (H2B-eGFP), STN (H2B-mCherry) / DLS (H2B-eGFP), and SNr (H2B-mCherry) / MO-SS (H2B-eGFP) (n = 8 mice per group). In total, 24 mice were injected across three retrograde target-pair groups: Pf (mCherry)/PPN (eGFP), STN (mCherry)/DLS (eGFP), and SNr (mCherry)/MO, SS (eGFP) (n = 8 mice per group). Injection coordinates (relative to bregma) and volumes were as follows: Pf: AP −2.30 mm, ML ±0.69 mm, 150 nL; PPN: AP −4.36 mm, ML ±1.25 mm, DV −3.25 mm, 100 nL; STN: AP −2.06 mm, ML ±1.50 mm, DV −4.40 mm, 50 nL; DLS: AP +0.50 mm, ML ±2.38 mm, DV −2.13 mm, 100 nL; MO-SS: AP +0.25 mm, ML ±1.25 mm, DV −0.75 mm, 100 nL; and SNr: AP −3.28 mm, ML ±1.35 mm, DV −4.38 mm, 150 nL. Related to Figures 2, 3, supplemental figures 4, and 5.

To sparsely label GPe neurons for single-cell whole-brain projectome reconstruction, we injected AAV-hSyn-Flex-eGFP into the GPe of Gnb4-Cre, Npr3-Cre, Penk-Cre, Pvalb-Cre, and Tle4-Cre mice. Injections were performed at the following coordinates (relative to bregma): GPe, AP −0.34 mm, ML ±2.15 mm, DV −3.50 mm, with a volume of 150 nL. This strategy enabled sparse but bright labeling of individual GPe neurons for subsequent whole-brain projectome reconstruction. Related to Figure 4, supplemental figures 2, 3, and 4.

To characterize axonal collateralization patterns of GPe^Pf^ and GPe^PPN^ neurons, we used a cell type- and projection-specific monosynaptic rabies tracing strategy. In vGlut2-Cre mice, AAV1-hSyn-DIO-TVA(66T)-3×HA-2A-N2C(G) (ZI Virology, 8.0 × 10¹² GC/mL) was injected into either the Pf (AP: −2.30 mm, ML: +0.69 mm, DV: −3.13 mm; 300 nL; n = 6 mice) or PPN (AP: −4.36 mm, ML: +1.25 mm, DV: −3.25 mm; 200 nL; n = 3 mice) to drive Cre-dependent expression of TVA and rabies glycoprotein. In the same animals, AAV5-EF1a-fDIO-eYFP (UNC Viral Vector Core) was injected into the GPe (AP: −0.34 mm, ML: −2.15 mm, DV: −3.50 mm; 400 nL) to enable Flpo-dependent reporter expression in presynaptic GPe neurons. After two weeks, EnvA-pseudotyped, glycoprotein-deleted CVS-N2c rabies virus encoding Flpo-mCherry (EnvA-N2cΔG-Flpo-mCherry; ZI Virology, 2.0 × 10⁹ ffv/mL) was injected into the same Pf or PPN site (500 nL) to initiate monosynaptic retrograde labeling of projection-defined GPe neurons. Related to Figures 4 and S6.

To identify MSNs projecting to segregated GPe^PPN^ and GPe^Pf^ subpopulations, we performed cell type- and projection-specific monosynaptic rabies tracing. AAVretro-EF1a-Flpo (Addgene #55637, 2.20 × 10¹³ GC/mL) was injected into the PPN (AP: −4.36 mm, ML: +1.25 mm, DV: −3.25 mm; 100 nL; n = 3 mice) or Pf (AP: −2.30 mm, ML: +0.69 mm, DV: −3.13 mm; 150 nL; n = 3 mice) of adult C57BL/6J mice to express Flpo recombinase in the corresponding GPe projection neurons. In the same animals, 300 nL of a helper AAV mixture containing AAV1-EF1a-FlpX-H2B-3×HA-2A-N2C(G) and AAVDJ-hSyn-FlpX-TVA950 (ZI Virology; each 8.0 × 10¹² GC/mL) was injected into the GPe (AP: −0.34 mm, ML: −2.15 mm, DV: −3.50 mm) to drive Flp-dependent expression of TVA950 and rabies glycoprotein N2C(G). Two weeks later, 300 nL of EnVA-N2cΔG-GFP (ZI Virology, 2.0 × 10⁹ ffv/mL) was injected into the GPe to initiate monosynaptic labeling. Related to Figures 6 and S7.

To perform projection-specific fiber photometry recordings from axon terminals of GPe projections to the PPN and Pf, we injected AAV1-hSyn-axon-targeted GCaMP6f (8.52 × 10^12^ GC/mL; gift from Dr. Lin Tian) into the GPe (AP −0.34 mm, ML +2.15 mm, DV −3.50 mm, 240 nL). A mono fiberoptic cannula (MFC_400/430-0.57_5mm_MF1.25_FLT, Doric Lenses) was then implanted above the axonal terminals in either the PPN (AP −4.36 mm, ML +1.25 mm, DV −2.95 mm) or the Pf (AP −2.30 mm, ML +0.50 mm, DV −2.88 mm, 15° angle). Related to Figures 5, and 7.

To perform projection-specific optogenetic stimulation of GPe axon terminals in the PPN or Pf, we bilaterally injected AAV2-hSyn-ChR2(H134R)-eYFP (n = 7, UNC Viral Vector Core, 3.10 × 10¹² GC/mL) or control AAV1-hSyn-EYFP (n = 7, UNC Viral Vector Core, 4.70 × 10¹² GC/mL) into the GPe (240 nL per hemisphere; AP: −0.34 mm, ML: ±2.15 mm, DV: −3.50 mm) of A2a-Cre mice. In the same mice, we also injected AAV1-EF1a-DIO-hChR2(H134R)-eYFP (n = 7, addgene #20298, 3.40 × 10¹³ GC/mL) or AAV1-EF1a-DIO-eYFP (n = 7, addgene #27056, 4.05 × 10¹³ GC/mL) into DLS (600 nL, AP: +0.5; ML: ±2.5; DV: 2.2). Optical fiber implants (OPT/FS-Flat-200/230, 5.50 mm length, Plexon) were then bilaterally placed above GPe axon terminals in either the PPN (AP: −4.36 mm, ML: ±1.25 mm, DV: −2.95 mm) or the Pf (AP: −2.30 mm, ML: ±0.50 mm, DV: −2.88 mm; 15° angle). In addition, a pair of optical fiber implants (OPT/FS-Flat-200/230, 4.00 mm length, Plexon) were then bilaterally targeted to DLS (AP: +0.5; ML: ±2.5; DV: 2.0) in individual mice. Related to Figures 5–7.

To perform projection-specific optogenetic inhibition of GPe axon terminals in the PPN or Pf, we bilaterally injected AAV5-hSyn-eNpHR3.0-EYFP (n = 10, UNC Viral Vector Core, 5.00 × 10¹² GC/mL) or control AAV1-hSyn-EYFP (n = 9, UNC Viral Vector Core, 4.70 × 10¹² GC/mL) into the GPe (240 nL per hemisphere; AP: −0.34 mm, ML: ±2.15 mm, DV: −3.50 mm) of A2a-Cre mice. In the same mice, we also injected AAV8-hSyn-Flex-Jaws-KGC-GFP (UNC Viral Core, 3.20 × 10¹² GC/mL) or AAV1-CAG-FLEX-tdTomato (UNC Viral Vector Core, 5.90 × 10¹² GC/mL) into DLS (600 nL, AP: +0.5; ML: ±2.5; DV: 2.2) for performing optogenetic inhibition of D2-MSNs in DLS. Optical fiber implants (OPT/FS-Flat-200/230, 5.50 mm length, Plexon) were then bilaterally placed above GPe axon terminals in either the PPN (AP: −4.36 mm, ML: ±1.25 mm, DV: −2.95 mm) or the Pf (AP: −2.30 mm, ML: ±0.50 mm, DV: −2.88 mm; 15° angle). In addition, a pair of optical fiber implants (OPT/FS-Flat-200/230, 4.00 mm length, Plexon) were then bilaterally targeted to DLS (AP: +0.5; ML: ±2.5; DV: 2.0) in individual mice. Related to Figures 5 and 6.

Once all fiber implants were lowered to their target coordinates, a small amount of clear Loctite Liquid Super Glue was applied to secure the ferrules in place. Dental cement was then applied according to the C&B METABOND Quick Cement System protocol to cement the implants to the skull. A holder (3-position, 0.079″ pitch gold PCB receptacle; Harwin Inc., manufacturer part number 122-7130342; DigiKey 952-1350-5-ND) was cemented adjacent to the DLS fiber implants, positioned along the midline and aligned parallel to the skull surface. This holder served as a head-mounted interface for connecting the wired inertial measurement unit (IMU) used for motion recording. Finally, additional layers of black Ortho-Jet powder and liquid acrylic resin (Lang Dental, USA) were applied over the METABOND layer to reinforce and light-shield the implant assembly.

### Histology and immunohistochemistry

Mice were deeply anesthetized with isoflurane and transcardially perfused with ice-cold 1x PBS followed by ice-cold 4% paraformaldehyde (PFA). Brains were postfixed overnight in 4% PFA, washed in 1x PBS, and cryoprotected in 30% sucrose for 48 hr at 4°C. Tissue was embedded in optimal cutting temperature (OCT) compound (Tissue-Tek), frozen, and stored at −80°C. Coronal sections (50 µm) were cut on a cryostat. Primary antibodies used were as follows: chicken anti-GFP (1:2000, Aves, GFP-1020), rabbit anti-RFP (1:2000, Rockland, 600-401-379), goat anti-mCherry (1:2000, Biorbyt, orb11618), rat anti-HA (1:2000, Roche, 11867423001), rabbit anti-Cre (1:500, BioLegend, 908001), and rabbit anti-PV (1:500, Swant, Pv27). Fluorophore-conjugated secondary antibodies were obtained from Jackson ImmunoResearch and Invitrogen. Free-floating immunohistochemistry was performed by incubating sections with primary antibodies overnight at room temperature, followed by three 1x PBS washes and incubation with secondary antibodies (1:1000) for 4 hr at room temperature. Sections were washed three times in 1x PBS and counterstained with DAPI (1:1000) for 15 min at room temperature. Sections were then mounted with Liquid Fluoromount-G® Mounting Medium (SouthernBiotech, 0100-01) and coverslipped for imaging.

### Imaging

Automated high-throughput imaging of tissue sections was performed on a custom-built automated slide scanner using an AZ100 microscope equipped with a 4x 0.4NA Plan Apo objective (Nikon Instruments Inc) and P200 slide loader (Prior Scientific), controlled by NIS-Elements using custom acquisition scripts (Nikon Instruments Inc.). Confocal imaging was performed on a W1-Yokogawa inverted spinning disk confocal using a 10x or 20x objective.

### Image processing, anatomical reconstructions, and analysis

Automated reconstruction of slide-scanned images was performed in ImageJ/Fiji using BrainJ as previously described^20^. BrainJ is an ImageJ-based analysis pipeline that enables high-throughput processing, registration, and quantification of whole-brain histological datasets^20^. Briefly, brain sections were aligned and registered using two-dimensional rigid-body registration^38^. A seven-pixel rolling-ball filter was applied to reduce background fluorescence. Machine-learning-based pixel classification was performed using Ilastik to identify cell bodies and neuronal processes^39^. Background-subtracted images from each fluorescence channel were imported into Ilastik, and representative examples of cell bodies, neurites, and background signal were manually annotated to train the classifier across all sections. Probabilistic classification outputs were inspected using live preview to ensure segmentation accuracy. The trained models were then applied to generate probability maps for each fluorescence channel, which were subsequently used for segmentation of labeled cells and processes. To assign anatomical location, three-dimensional registration was performed using Elastix relative to a reference brain^39^. Coordinates of detected cells and processes were projected into the Allen Brain Atlas Common Coordinate Framework^40^. Data visualization was performed using ImageJ and Brainrender^21^, and quantitative analyses were conducted using custom MATLAB and Python scripts.

To quantitatively define novel GPe projection targets, we calculated projection density for each annotated brain region as the total GPe axonal projection volume within that region divided by the region’s total volume. Projection density values were computed for each brain and then averaged across animals (n = 7 mice). A region was classified as a GPe target if its mean projection density exceeded a predefined threshold. We used the pedunculopontine nucleus (PPN) as a biologically validated reference region to establish this threshold, as it ranks 64th among all quantified GPe projection targets and has independent functional validation in this study. Regions with projection densities greater than or equal to the mean PPN projection density were classified as bona fide GPe targets. Because this thresholding strategy relies on a single benchmark region and emphasizes specificity, it likely provides a conservative estimate and may underestimate the total number of previously unrecognized GPe projection targets.

### Single-nucleus RNA sequencing

#### Dissection, nuclear isolation and flow cytometry

Single-nucleus preparations were performed using microdissected GPe tissue pooled from 24 12-week-old C57BL/6J mice, each injected with AAVretro-hSyn-H2B-mCherry and AAVretro-hSyn-H2B-eGFP into two distinct GPe output targets, following the Allen Institute V1 protocol. Briefly, each pooled sample was homogenized in a chilled 2 mL Dounce homogenizer (Sigma D8938) using 3.5 mL of freshly prepared homogenization buffer containing 3.375 mL nuclear isolation medium (NIM; 10 mM Tris-HCl, 250 mM sucrose, 25 mM KCl, 5 mM MgCl₂), 3.5 μL of 100 mM DTT (Promega P1171), 70 μL of 50× protease inhibitor cocktail (Promega G6521), 17.5 μL of 40 U/μL RNAsin Plus (Promega N2615), and 35 μL of 10% Triton X-100 (Sigma T8787). The homogenate was filtered through a 30 μm cell strainer (Miltenyi Biotec 130-098-458) and centrifuged at 900 × g for 10 min at 4 °C. The resulting pellet was resuspended in 500 μL of freshly prepared blocking buffer consisting of 410 μL nuclease-free water, 50 μL 10× PBS, 40 μL 10% OmniPur BSA (Sigma 2905), and 2.5 μL of 40 U/μL RNAsin Plus. The nuclear suspension was incubated on ice for 15 min. To enrich for neuronal nuclei, the suspension was labeled with mouse anti-NeuN-488 antibody (1:500; MilliporeSigma) and incubated on ice for 30 min, followed by centrifugation at 400 × g for 5 min. The pellet was resuspended in 500 μL blocking buffer and labeled with 0.5 μL DAPI. Prior to DAPI labeling, 5 μL of nuclear suspension was mixed with 5 μL Trypan Blue and loaded onto a disposable hemocytometer slide (Invitrogen C10228) to estimate nuclear yield and assess nuclear integrity. Fluorescence-activated nuclei sorting (FANS) was performed using a Beckman Coulter MoFlo Astrios EQ cell sorter. The instrument was configured with a 405 nm laser for excitation of DAPI-labeled nuclei, a 488 nm laser for GFP excitation and scatter detection, and a 560 nm laser for mCherry excitation. The sample and collection chambers were maintained at ∼7 °C during sorting. DAPI fluorescence was detected using a 448/59 band-pass filter (PMT 341 V, amplifier gain 3). GFP fluorescence was detected using a 513/26 band-pass filter (PMT 360 V, amplifier gain 3). mCherry fluorescence was detected using a 610/20 band-pass filter. DAPI⁺/GFP⁺, DAPI⁺/mCherry⁺, and DAPI⁺/GFP⁺/mCherry⁺ nuclei were sorted as single nuclei into 8-well strip tubes containing SMART-seq lysis buffer with RNase inhibitor (0.17 U/µL; Takara ST0764).

#### Nuclear isolation

Nuclei were isolated using the RAISINs protocol with slight modifications^49^, as described (https://dx.doi.org/10.17504/protocols.io.8epv517xdl1b/v5).

#### Retro-SMART-seq

Following nuclear isolation, individual nuclei were sorted into 8-well strips containing SMART-seq lysis buffer with RNAse inhibitor (0.17 U/µL; Takara Cat#ST0764), frozen on dry ice, stored at -80°C and then shipped to the Allen Institute on dry ice. SMART-seq libraries were prepared and sequenced to a target read depth of 500,000 reads per nucleus, as previously described^28,29^. The full SMART-seq sample preparation, library preparation and bioinformatic workflow is documented at https://celltypes.brain-map.org under “Documentation.”

#### 10x Genomics single nucleus sequencing

For 10x sequencing of unlabeled GPe neurons, six 9-week-old C57BL/6J mice were used. Animals were euthanized in a CO₂ chamber and decapitated. Brains were rapidly removed from the skull and submerged in ice-cold artificial cerebrospinal fluid (aCSF) containing 92 mM NMDG, 2.5 mM KCl, 1.25 mM NaH₂PO₄, 30 mM NaHCO₃, 20 mM HEPES, 25 mM glucose, 5 mM Na-ascorbate, 3 mM Na-pyruvate, 0.5 mM CaCl₂·2H₂O, and 10 mM MgSO₄·7H₂O. Coronal brain slices (400 μm thick) were prepared at 4 °C using a vibrating-blade microtome (V1200S, Leica, USA). The GPe subregion was microdissected in aCSF under a fluorescence dissection microscope (Leica M165FC) and transferred to microcentrifuge tubes. Excess aCSF was removed, and the samples were immediately flash-frozen on dry ice. Following RAISINs-based nuclear isolation, nuclei were quantified and processed using Chromium’s KITXYZ and KITZYX (catalog numbers). We prepared 10x Genomics snMultiome and snRNA-seq (v3.1) samples as previously described https://dx.doi.org/10.17504/protocols.io.bp2l61mqrvqe/v3; https://dx.doi.org/10.17504/protocols.io.dm6gpwd8jlzp/v3). We loaded ∼10,000 nuclei per port. We targeted sequencing depth of 120,000 reads per nucleus and achieved 123,893 reads/nucleus on average (Supplemental Table 1, Sheet “10xMetrics”).

#### Processing RNA-seq data

Processing of single-nucleus transcriptomic data was performed as previously described for SMART-seq^38^ and 10x Genomics data^48^.

#### Analysis of integrated RNAseq datasets

Retro-SMART-seq v4, (SSv4, Clontech), 10x Genomics single-nucleus multiome sequencing, and 10x Genomics single-nucleus RNA sequencing (v3.1) were independently subjected to cell or nucleus QC filtering (**Figure S2A**). For 10xGenomics data, we kept nuclei that satisfied these initial QC filters: doublet finder^50^ doublet_score below 0.3, *Malat1* log_2_CPM ≥ 11, fewer than 10% of transcripts mapping to mitochondrial genes, total UMI counts below the mean + 3 standard deviations, gene counts above the 5^th^ percentile, and gene counts below the mean + 3 standard deviations from the mean. For Retro-SMART-seq (v4), we kept nuclei with over 1000 genes detected, more than 100,000 reads, over 75% alignment rate and cg dinucleotide complexity over 0.5. We kept genes only if they were expressed in a minimum of three nuclei (min.cells = 3).

Across modalities, raw count cell x gene matrices were concatenated. Multiple integration methods and rounds of integration were used to correct for batch and technological artifacts and filter to the desired clusters (**Figures 2A**, and **S2A**). These steps are fully documented at https://github.com/AllenInstitute/gpe. The initial whole dataset integration was performed with multiple approaches: scVI and scANVI^51^, Harmony^52^, RPCA and CCA^53^. Integration benchmarking was performed with scIB^54^. The scANVI integration of the entire (neuronal and non-neuronal) single-nucleus sequencing GPe dissectate dataset was selected as the best integration (Fig. S2A). This dataset was filtered to neurons, re-integrated, then subset by removing clusters and cells that represented likely doublets, cholinergic neurons and neurons that were likely contamination from nearby regions (such as MSNs) based on the ABC Atlas. After performing these subsetting operations, the dataset was re-integrated, yielding a final integrated GPe GABAergic neuron dataset (3,634 neurons).

#### Transcriptomic data visualization and analysis

The final transcriptomic dataset was clustered with Louvain clustering (resolution 0.3) and analyzed with Seurat (5.1.0). GPe GABAergic neuronal clusters were given cell type labels based on their cluster number and key marker genes. These clusters were nearly homogeneous for mapped ABC Atlas (version: AIT21; CCN20230722) cell clusters (type level) (Fig. 2B, Supplemental Table 1). In the case of cluster 0891 NDB-SI-MA-STRv Lhx8 Gaba_4, our clustering further subdivided this cluster, yielding 0891 NDB-SI-MA-STRv Lhx8 Gaba_4_A (2_GPe_Cdh18) and 0891 NDB-SI-MA-STRv Lhx8 Gaba_4_B (3_GPe_Mboat1). Differential expression was calculated using FindAllMarkers (Seurat) using the entire dataset and the transcriptomic clusters as the grouping scheme (Fig. 2D; Supplemental Table 1). Differential expression was also calculated for projection-specific populations, using only the Retro-seq data as the starting matrix (Fig. 3). Figures were produced in R using Seurat 5.1.0, ggplot2, scCustomize, patchwork, ggsankey, and ComplexHeatmap. All genes were scaled across all cells (Seurat::ScaleData) prior to expression visualization.

#### Spatial soma prediction

To predict the soma locations of our single-nucleus transcriptomic data, we trained regression models to map gene expression to 3D spatial coordinates using MERFISH data as a reference. We selected only 8 of the predominant neuronal subclasses in the GPe to use as the training/test set. We kept only genes shared between the MERFISH and snRNA-seq datasets. The MERFISH neuronal dataset was split 80/20 into training and test sets. Three model types were trained independently for each spatial axis (X, Y, Z): multiple linear regression, Random Forest (300 trees, mtry = √p and XGBoost (max depth = 6, η = 0.1, 500 rounds). Model performance on the test set was assessed using MAE, RMSE and R^2^. All three trained models were then applied to Retro-seq SMART-seq v4 neurons using the expression of the overlapping gene panels as input and storing the predicted coordinates as cell-level metadata for plotting. All analyses were performed in R 4.2.3 using Seurat 5.1.0, randomForest, xgboost, and caret.

### fMOST imaging, Whole neuron morphology (WNM) reconstruction and analysis

As described previously^34,55^, resin-embedded, GFP-labeled brains underwent chemical reactivation to recover GFP fluorescence and facilitate wide-field or two-photon block-face imaging. GFP was imaged with an excitation wavelength of 488 nm and a bandpass emission filter of 510–550 nm. A 40X water-immersion lens with NA 0.8 was used to provide an optical resolution (at 520 nm) of 0.35 µm in XY axes and voxel size of 0.35 x 0.35 x 1.0 µm, appropriate for neuron reconstruction. Whole-brain imaging produced a 15–20 TB dataset of approximately 10,000 coronal planes per brain. These data are freely available at https://spelunker.cave-explorer.org/#!%7B%22layers%22:%5B%7B%22type%22:%22new%22%2C%22source%22:%22%22%2C%22tab%22:%22source%22%2C%22name%22:%22new%20layer%22%7D%5D%2C%22selectedLayer%22:%7B%22visible%22:true%2C%22layer%22:%22new%20layer%22%7D%2C%22layout%22:%224panel-alt%22%7D. The metadata for each neuron is recorded in Supplemental Table 2.

### Whole neuron morphology (WNM) reconstruction and analysis

Vaa3D-TeraVR was used for WNM reconstructions of fMOST images. The den- drites and the complete local and long-range axonal arbor was traced using the virtual finger or polyline tool. Post-processing steps were run on completed reconstructions to ensure that there were no errors (i.e., breaks or loops).

### fMOST image registration to CCF

Whole brain fMOST images were registered to the average mouse brain tem- plate of CCFv3 using DeepMAPI^35^. Once images were CCF-aligned, the reconstructed neurons were transformed into the CCFv3 space using the generated deformation fields.

### Calculating the WNM projection matrix

CCF-registered reconstructions were translated such that all somas were positioned in the left hemisphere. SWC files were subsequently resampled to ensure uniform spacing between nodes. To quantify the pattern of axonal projection targets, a projection matrix was derived based on reconstruction node counts per anatomical target structure. Target regions were represented in both ipsilateral and contralateral hemispheres. To better reflect the targets that were synaptically innervated by a neuron, versus those with just fibers of passage, only regions containing a branch or tip node were included in the projection matrix.

### Determining neuron projection groups and visualizing projection data

Neurons located within 20µm of the GPe border were excluded from analysis, as were neurons from *Chat, Tle4,* and *Gnb4* Cre lines due to border proximity, cholinergic characteristic, or aberrant projection patterns. This left 20 reconstructed neurons with soma localized to the GPe. Ipsilateral and contralateral axon lengths were summed for each brain region from the branch/tip masked projection matrix, so as not to count pass-through neurons. Total neuron length was calculated from the un-masked matrix. The branch/tip projection matrix was normalized by each neuron’s total axonal length. Principal component analysis (PCA) was performed on the normalized matrix, and the first 10 components were retained. Neurons were clustered using hierarchical clustering with Ward’s D2 linkage based on the PCA scores, with the optimal cluster number (k = 4) determined by inspecting the within-cluster sum of squares elbow plot. Brain regions were organized at multiple hierarchical levels using the Allen Mouse Brain Common Coordinate Framework (CCF 2020) parcellation: projection targets within the striatum, midbrain, and thalamus were represented at the “child” structure level, while other regions were collapsed to their “parent” structure. A GPe boundary mask was created from using the Python package morph_utils (https://github.com/MatthewMallory/morph_utils). Projection patterns, soma location relative to the boundary mask and cluster assignments were visualized using the R packages tidyHeatmap (1.12.2) and ggplot2 (3.5.2). Projection matrices and analysis code is available at https://github.com/AllenInstitute/GPe_projection.

### Wired inertial measurement unit (IMU) and the WEAR system

The IMU used in this study was part of the WEAR motion sensor system developed by the Champalimaud Hardware Platform and the Costa laboratory (https://www.cf-hw.org/harp/wear). The device is an ultra-lightweight (200 mg) 9-axis motion sensor comprising accelerometer, gyroscope, and magnetometer components. In this study, accelerometer and gyroscope signals were used for behavioral quantification. Motion data were sampled at 200 Hz. The IMU communicates with a HARP-based base station (Champalimaud Hardware Platform), enabling hardware-level synchronization through digital inputs and outputs, camera trigger outputs, analog input, and clock synchronization signals. The IMU can be controlled via software commands or hardware pins. Data streams from the IMU, video cameras, and other acquisition systems were integrated and synchronized using Bonsai visual reactive programming software (Bonsai v2.4, https://bonsai-rx.org/). Time stamps from all devices were aligned using custom MATLAB scripts for downstream behavioral analysis. WEAR hardware and firmware designs are open source and available through the Champalimaud Hardware Platform repository.

### Open-field experiments and synchronized motion and video acquisition

Open-field experiments were conducted in a white acrylic arena (40 × 40 cm) placed inside a custom-built sound-attenuating chamber (1 m × 1 m). Mice were habituated to the head-mounted devices and the open-field arena for 2 days (1 hour per mouse per day) prior to experimentation. On the day of recording, mice were lightly anesthetized with 2%–3% isoflurane to facilitate attachment and removal of the optogenetic patch cord (Plexon, OPT/PC-LC-LCF-200/230-HD), fiber photometry mono fiberoptic patch cord (Doric, MFP_400/430/1100-0.57_2m_SMA-MF1.25_LAF, low autofluorescence), and the wired IMU sensor. Recordings began 15 min after recovery from anesthesia. Behavioral, optogenetic, or fiber photometry sessions lasted 40–90 min depending on the experimental condition. For construction of the behavioral clustering library used in closed-loop optogenetic experiments, 1 hour open-field recordings were collected. During closed-loop optogenetic sessions, actions were self-paced, and 60 trials were collected within 30–90 min sessions, with total duration determined by mouse-initiated behaviors. Fiber photometry recordings of calcium activity from GPe axon terminals were collected in 30 min sessions. IMU data were sampled at 200 Hz using a HARP base station. The same base station generated 30 Hz TTL camera trigger signals to control a top-mounted behavioral camera (Flea3, Point Grey Research) recording at 30 frames per second. A custom Bonsai workflow was used to initiate acquisition and synchronize IMU signals, behavioral video, optogenetic stimulation timestamps, and fiber photometry recordings. Behavioral video and associated timestamps were recorded continuously throughout each session.

### Unsupervised behavior clustering

To unbiasedly identify the natural action repertoire of individual mice, we performed unsupervised behavioral classification using affinity propagation applied to IMU sensor data, following a framework similar to previously described approaches^10,35,41^. In contrast to the original implementation described by Klaus et al. study^10^, which used three behavioral features derived from accelerometer and video measurements (total body acceleration, gravitational acceleration along the anterior–posterior axis, and video-derived head–body rotation), our analysis incorporated four features extracted directly from IMU signals to capture additional aspects of body dynamics. Four features were extracted from the sensor data: 1) gravitational acceleration along the anterior–posterior axis for the discrimination of postural changes, GAap; (2) Raw sensor acceleration along the dorsal–ventral axis to quantify movement momentum, ACCdv; (3) Dorsal–ventral axis of the gyroscope to extract head head–body rotational information, GYRdv; (4) Total body acceleration to differentiate the resting state from movement. Total body acceleration (TotBA) was defined as:

where BAap, BAml and BAdv represent the body acceleration of the anterior–posterior, medial–lateral and dorsal–ventral axis, respectively. Each BA component was calculated by median filtering the raw acceleration signals followed by a fourth-order Butterworth high-pass filter (0.5 Hz cutoff). For estimation of gravitational acceleration, the BA components were subtracted from the median-filtered raw signal along each axis. All four time-series features were segmented into non-overlapping 300 ms windows. For each window, feature values were discretized using fixed thresholds to generate summary distributions. For GAap and ACCdv, ten evenly spaced thresholds were used, with two additional bins capturing values outside the defined limits to approximate the distribution within each window. For GYRdv, five thresholds (0, ±50, ±100) were used to discriminate left and right rotational movements. For TotBA, a single threshold was used to separate movement from resting states. This threshold was kept constant across experiments and defined as the average value separating the bimodal distribution of log(TotBA). For each 300 ms segment, four histograms were generated, one for each feature. These histograms were normalized to probability distributions and used to compute pairwise similarities between segments. We used the Earth Mover’s Distance (EMD) as a measure of similarity^42^: where dEM represents the sum of normalized EMD distances across the four features (GAap, ACCdv, GYRdv, and TotBA). The normalization constrains similarity values to the range [-1,0], where −1 represents maximal dissimilarity and 0 represents identical distributions. The similarity matrix was then clustered using affinity propagation, with the preference parameter set to the minimal value of the similarity matrix, which produced a stable number of behavioral clusters across recordings. Compared with the original implementation described by Klaus et al.^10^, the similarity metric used here incorporates four behavioral features rather than three. Affinity propagation clustering produced a set of discrete behavioral states representing the natural action repertoire of the animal. These clusters were used to construct the behavioral library for downstream behavioral analyses and closed-loop optogenetic experiments.

### Closed-loop optogenetics set-ups during open-field experiment

To unbiasedly identify the natural action repertoire of individual mice and generate a behavioral library for action-specific closed-loop optogenetic manipulation, we first performed one-hour open-field recordings and processed the IMU sensor data to obtain behavior clusters for each individual mouse. This is the ground-truth cluster library for this mouse. Video frames corresponding to each behavioral cluster were then extracted and stitched together to generate cluster-specific video clips. These clips were manually inspected to identify clusters corresponding to immobility and locomotion for each mouse, which were subsequently used as target actions for online closed-loop optogenetic manipulation. Once the target clusters were defined, a testing session was conducted to verify the detection accuracy of the selected behavioral clusters before performing the optogenetic experiments.

During closed-loop optogenetic sessions, mice were lightly anesthetized with 2–3% isoflurane to facilitate attachment and removal of the optogenetic patch cord (Plexon, OPT/PC-LC-LCF-200/230-HD) and the wired IMU sensor. Recordings began 15 min after recovery from anesthesia. An online closed-loop Bonsai script was used to acquire IMU sensor data and record behavioral video as described above for the open-field experiments. In parallel, IMU sensor data were streamed to a custom MATLAB script that performed real-time behavioral clustering. For each incoming 300 ms segment of IMU sensor data, motion feature histograms were computed and compared to the exemplar clusters in the ground-truth behavioral library derived from the one-hour open-field recording of the same mouse. The motion feature histogram was assigned to the action for which comparison with the exemplar produced the lowest EMD score, corresponding to the most similar behavioral cluster. When the identified cluster matched the predefined target action (immobility or locomotion), the Bonsai script generated a trigger signal that was sent via an Arduino board to a Plexon PlexBright 4-channel optogenetic controller to initiate optogenetic stimulation. The latency between detection of the target action and delivery of the optogenetic trigger was estimated to be 35–55 ms^35^. Timestamps for IMU data, behavioral video, behavioral variables, and optogenetic triggers were recorded and synchronized through the Bonsai acquisition script. During closed-loop optogenetic sessions, actions were self-paced, and 60 trials were collected within 30–90 min sessions, with the total duration determined by mouse-initiated behaviors.

For offline analysis of closed-loop optogenetic effects, stimulation-triggered data were aligned to the onset of optogenetic stimulation. Distance and speed were calculated from the animal centroid extracted online in Bonsai and converted from pixels to centimeters. Total body acceleration was quantified from the raw IMU signal sampled at 200 Hz and aligned to the optogenetic trigger onset. For each trial, the acceleration trace during the 5 s stimulation period was extracted, and the mean total body acceleration across this window was used for summary statistics. Behavioral cluster identity was analyzed in 300 ms bins derived from IMU sensor data. Locomotion clusters were defined manually for each mouse by reviewing cluster-specific stitched video clips from each recording session. Forward probability was defined as the fraction of trials in which at least one forward locomotion cluster occurred within the 5 s optogenetic manipulation window. Forward sequence probability was defined as the fraction of trials in which at least one forward locomotion sequence occurred within the same window. A forward locomotion sequence was defined as two consecutive forward locomotion clusters (600 ms total duration), with each cluster representing a 300 ms behavioral segment.

### Lever-pressing task with optogenetic manipulation and fiber photometry

Mice were trained to perform a fixed-ratio operant lever-press task using custom-built operant chambers placed inside item24 sound-attenuating boxes measuring 0.5 meter × 0.5 meter × 0.5 meter or 0.5 meter × 0.5 meter × 1.0 meter (length × width × height). The lever-press box consisted of an enclosed acrylic chamber with interior dimensions of 24 cm (depth), 15.6 cm (width), and 20.66 cm (height). Behavioral control and event detection were implemented using pyControl^43^, a behavioral experiment control system based on a MicroPython microcontroller. Sucrose solution (10%) was delivered through a solenoid valve (LHDA1231515H, The Lee Company) via tubing into the reward magazine (5 μL per reward). Licks at the reward port were detected using an infrared beam sensor. Mouse position and behavior within the chamber were monitored using a side camera (Flea3, Point Grey Research), while overall behavior was recorded using a second camera (Flea3, Point Grey Research) positioned above the chamber. Both cameras recorded at 30 Hz. During training, mice were maintained under food restriction and received approximately 1.5–2.5 g of regular chow daily after training sessions to maintain body weight at approximately 85% of baseline weight. The training paradigm began with 2 days of magazine training, during which sucrose rewards were delivered on a random time schedule to allow mice to learn to retrieve rewards from the reward magazine. This was followed by 7 days of continuous reinforcement (CRF; one press = one reward) consisting of 2 days each of CRF5 and CRF15 and 3 days of CRF30, with progressively increasing maximum reward limits per session. Following CRF training, mice were trained under a progressively more demanding fixed-ratio schedule consisting of 2 days of FR4 (four presses = one reward) followed by 14 days of FR8 training (eight presses = one reward). Under the FR8 schedule, animals were required to complete a sequence of eight lever presses to obtain a sucrose reward (maximum of 30 rewards per session). Over the course of training, mice showed progressive increases in total lever presses and press rate and gradually organized their behavior into self-paced bouts of rapid lever-press sequences. A sequence of lever presses was defined as a bout of presses occurring before delivery of an individual reward. Mice were maintained under the FR8 schedule for 14 days before fiber photometry recordings or optogenetic experiments were conducted. To perform optogenetic excitation of GPe projections to the Pf or PPN, delivery of blue light was triggered through TTL pulses generated by pyControl upon detection of the first lever press in a sequence. TTL pulses were timestamped in Bonsai and transmitted via an Arduino board to a Plexon PlexBright 4-channel optogenetic controller to initiate optogenetic stimulation. In each session, 15–30 stimulation trials (true trials) and a similar number of false trials were recorded. Timestamps of lever presses, magazine entries, and optogenetic stimulation events were recorded and synchronized using the Bonsai acquisition script.

To quantify how optogenetic stimulation affected ongoing lever-press action sequences during the FR8 task, several behavioral variables were defined and extracted from timestamped behavioral events. The number of lever presses during stimulation was defined as the number of lever presses occurring within the 5-second optogenetic stimulation window following the first lever press of an FR8 sequence, providing a measure of the immediate effect of stimulation on the ongoing lever-press bout. The number of lever presses post-stimulation was defined as the number of lever presses occurring after the stimulation window within the same sequence before reward delivery. The number of lever presses per reward was calculated as the total number of lever presses within a sequence divided by the number of rewards obtained. Sequence duration was defined as the time interval between the first and the last lever press within a reward-earning sequence, reflecting the temporal structure and speed of sequence execution. The reinforcement rate was defined as the number of rewards obtained per minute during the session and provided a global measure of task performance. The inter-press interval (IPI) was defined as the time interval between consecutive lever presses within a sequence, and for each sequence the mean IPI was calculated across all presses to quantify the rhythm of lever-press execution. All behavioral metrics were computed separately for stimulation and control conditions and averaged across trials within each animal prior to group-level comparisons.

All behavioral metrics were computed separately for stimulation and control conditions and were averaged across trials within each animal prior to group-level comparisons. Behavioral variables derived from the FR8 task were statistically analyzed using a two-factor design. For each animal, behavioral metrics were first averaged across trials within each session. Group comparisons were performed using a two-way repeated-measures ANOVA, with stimulation condition (LED off, GPe→Pf stimulation, GPe→PPN stimulation) as the within-subject factor and virus group (ChR2 or YFP control) as the between-subject factor. When significant main effects or interactions were detected, post hoc pairwise comparisons were performed using Tukey’s multiple-comparison test. Statistical analyses were conducted using GraphPad Prism or MATLAB. Data are presented as mean ± s.e.m., and individual data points represent individual animals.

### Fiber photometry recording and analysis

One month after surgery, mice were habituated to the fiber photometry patch cord (Doric, MFP_400/430/1100-0.57_2m_SMA-MF1.25_LAF, low-autofluorescence Mono fiberoptic patch cord) and the wired IMU sensor in a white acrylic open-field arena (40 × 40 cm) placed inside a custom sound-attenuating chamber (1 meter × 1 meter) for 2 consecutive days (1 hour per mouse per day). The branching fiber patch cord contains four fibers, of which two were used for recording from the Pf and PPN simultaneously. On the day of recording, axonal calcium activity was recorded for 30 minutes during free exploration. Behavioral video was acquired at 30 Hz and IMU motion sensor data were acquired at 200 Hz. All data streams were synchronized using Bonsai. Fiber photometry recordings during open-field experiments were performed using a Neurophotometrics FP3002 system. Fluorescence signals were acquired at 60 Hz using alternating excitation of 470 nm and 415 nm LEDs, resulting in an effective sampling rate of 30 Hz per wavelength. Excitation at 470 nm was used to record calcium-dependent fluorescence, and excitation at 415 nm served as the isosbestic control channel. Raw fluorescence signals from both channels were exported and analyzed using custom MATLAB scripts. To correct for photobleaching and motion-related artifacts, the 415-nm control signal was first fitted using a biexponential function and then scaled to the 470-nm signal using robust linear regression. The normalized fluorescence trace was calculated as the ratio between the 470-nm signal and the fitted 415-nm control signal and expressed as ΔF/F. To relate neural activity to behavior, IMU sensor data were clustered using affinity propagation as described above. Cluster identities were verified by generating cluster-specific video clips and manually inspecting the associated behavioral videos to identify the actions represented by each cluster. Locomotion onset was defined as the transition into two consecutive locomotion clusters (600 ms), with each cluster representing a 300 ms behavioral segment derived from IMU sensor data. Requiring two consecutive bins ensured reliable detection of sustained locomotion initiation. For event-aligned analysis, peri-event fluorescence traces were extracted using a 3 s pre-event and 3 s post-event window around each locomotion onset. Each trace was baseline-corrected and normalized using a robust baseline z-score computed from the median and median absolute deviation (MAD) of a pre-event baseline window defined from −2.1 s to −1.0 s relative to locomotion onset. This robust normalization approach minimizes the influence of large calcium transients and baseline fluctuations and allows reliable comparison of event-aligned responses across trials and animals. Z-scored peri-event traces were averaged across trials within each session to obtain a mean peri-event response for each animal and region. To quantify locomotion-related neural activity, z-score delta was calculated as the peak-to-trough amplitude of the trial-averaged peri-event z-score trace, defined as the difference between the maximum and minimum z-score values within the analysis window.

Following completion of the open-field photometry experiments, the same cohort of mice was subsequently trained in the lever-pressing task described above. During these experiments, fiber photometry recordings were obtained from GPe projections to the Pf and PPN simultaneously while mice performed the fixed-ratio eight (FR8) operant task. For lever-pressing experiments, mice were connected to the branching fiber photometry patch cord prior to each behavioral session. Fiber photometry signals were acquired using the same FP3002 acquisition configuration (60 Hz alternating excitation; 30 Hz effective sampling per wavelength). Behavioral events, including lever presses and magazine entries, were generated by the pyControl system and transmitted to Bonsai for synchronization with photometry recordings. For event-aligned analysis, neural activity was aligned to the first lever press of each FR8 sequence, which marked the initiation of an action sequence. Peri-event fluorescence traces were extracted around the first lever press and normalized using the same robust baseline z-score procedure described above. Z-scored traces were averaged across trials within each session to obtain a mean peri-event response for each animal and region. To quantify task-related neural activity modulation, z-score delta was calculated as the peak-to-trough amplitude of the trial-averaged peri-event z-score trace, defined as the difference between the maximum and minimum z-score values within the analysis window. This metric captures the magnitude of neural modulation associated with the initiation of lever-press action sequences. Negative z-score deflections indicate decreases in axonal calcium activity relative to baseline, reflecting reduced presynaptic activity in the recorded projection pathway at the onset of the action sequence.

## Statistical analysis

All sample sizes, statistical tests, and significance values are reported in the Results or figure legends. Data are presented as mean ± SEM unless otherwise noted. Statistical analyses were performed using GraphPad Prism (v8.0.2). For comparisons involving repeated measurements within the same subjects, one-way repeated-measures ANOVA with Geisser–Greenhouse correction was used, followed by post hoc multiple comparisons using either Dunnett’s test (when comparing conditions to a control) or Sidak’s test (for pairwise comparisons). For comparisons between two groups, two-tailed paired or unpaired t tests were used as appropriate. For distributional comparisons, two-sample Kolmogorov–Smirnov (KS) tests were applied. Statistical significance was defined as P < 0.05. Significance levels are indicated as follows: ****P < 0.0001, ***P < 0.001, **P < 0.01, *P < 0.05; ns, not significant.

## Notes

### Competing Interest Statement

The authors have declared no competing interest.

