## Supplemental Figures for "The External Globus Pallidus is a Basal Ganglia Output Hub with Action-Specific Circuits"

Supplemental Figure 1, related to Figure 1

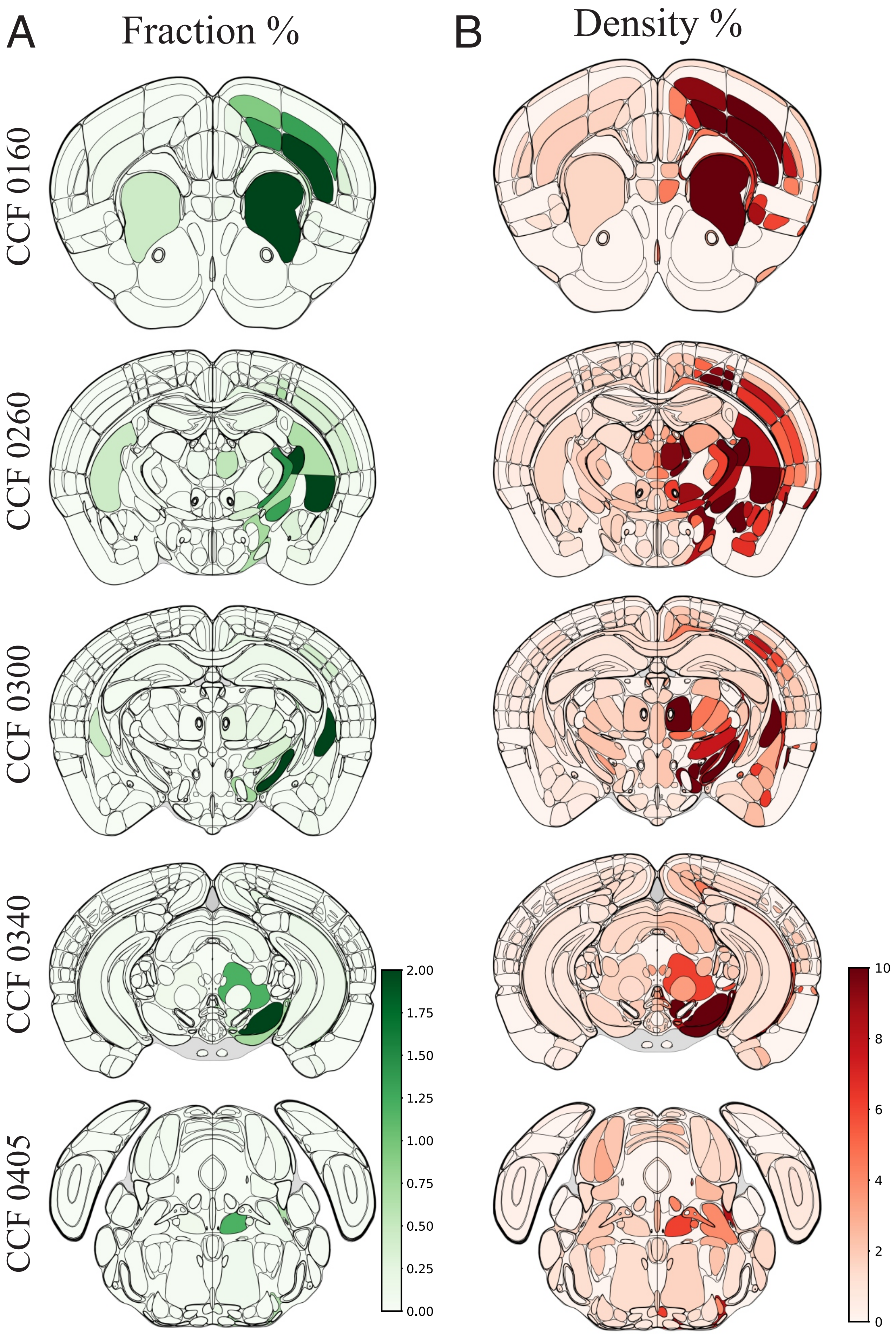

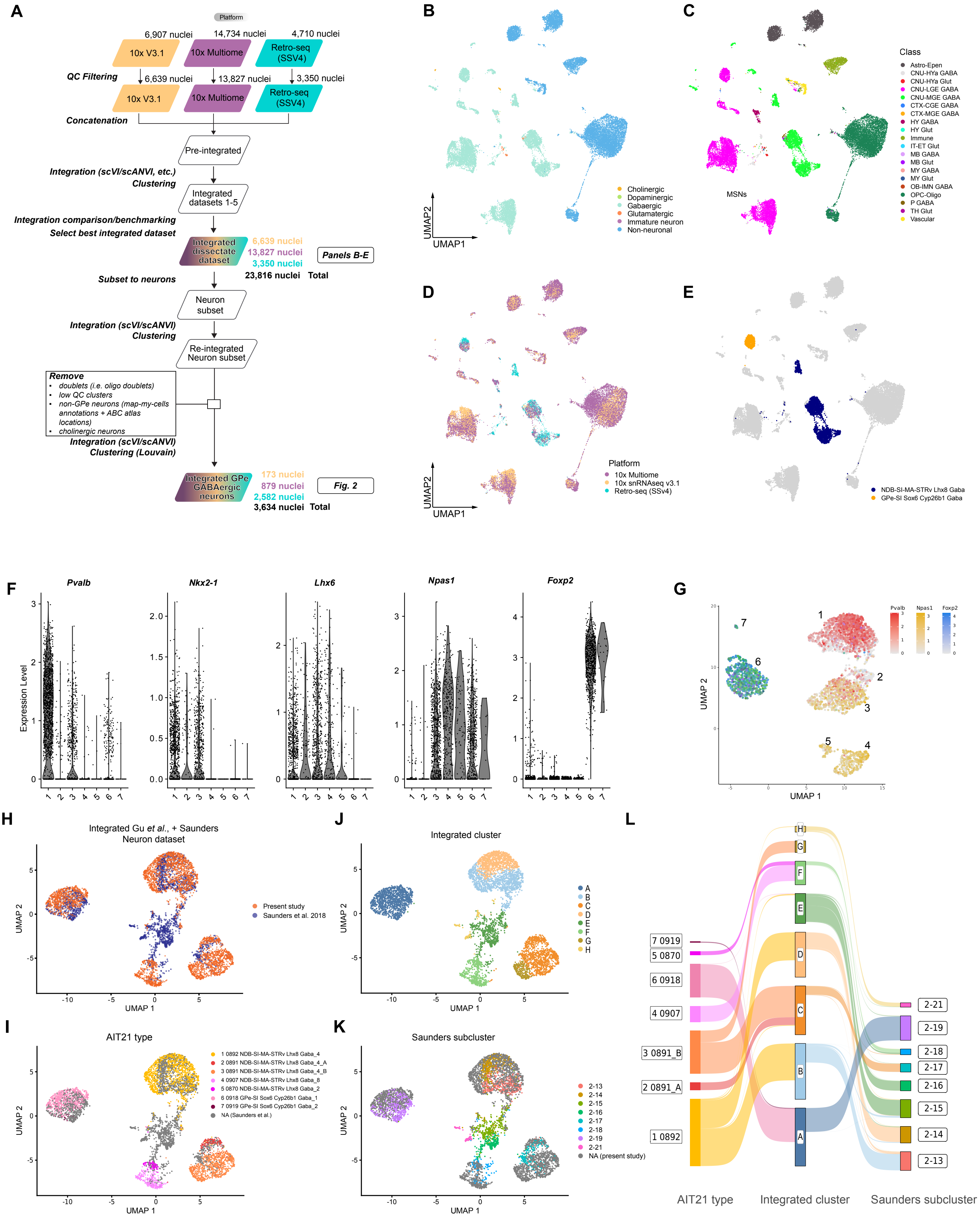
A
Platform
6,907 nuclei
14,734 nuclei
4,710 nuclei
10x V3.1
10x Multiome
Retro-seq (SSV4)
QC Filtering
6,639 nuclei
13,827 nuclei
3,350 nuclei
Concatenation
Pre-integrated
Integration (scVI/scANVI, etc.)
Clustering
Integrated datasets 1-5
Integration comparison/benchmarking
Select best integrated dataset
Integrated dissectate dataset
6,639 nuclei
13,827 nuclei
3,350 nuclei
23,816 nuclei Total
Panels B-E
Subset to neurons
Neuron subset
Integration (scVI/scANVI)
Clustering
Re-integrated Neuron subset
Remove
doublets (i.e. oligo doublets)
low QC clusters
non-GPe neurons (map-my-cells annotations + ABC atlas locations)
cholinergic neurons
Integration (scVI/scANVI)
Clustering (Louvain)
Integrated GPe GABAergic neurons
173 nuclei
879 nuclei
2,582 nuclei
3,634 nuclei Total
Fig. 2
B
UMAP2
UMAP1
Cholinergic
Dopaminergic
Gabaergic
Glutamatergic
Immature neuron
Non-neuronal
C
Class
Astro-Epen
CNU-HYa GABA
CNU-HYa Glut
CNU-LGE GABA
CNU-MGE GABA
CTX-CGE GABA
CTX-MGE GABA
HY GABA
HY Glut
Immune
IT-ET Glut
MB GABA
MB Glut
MY GABA
MY Glut
OB-IMN GABA
OPC-Oligo
P GABA
TH Glut
Vascular
MSNs
D
Platform
10x Multiome
10x snRNAseq v3.1
Retro-seq (SSV4)
UMAP2
UMAP1
E
NDB-SI-MA-STRv Lhx8 Gaba
GPe-SI Sox6 Cyp26b1 Gaba
F
Pvalb
Nkx2-1
Lhx6
Npas1
Foxp2
Expression Level
G
Pvalb
Npas1
Foxp2
UMAP 2
UMAP 1
H
Integrated Gu et al., + Saunders Neuron dataset
Present study
Saunders et al. 2018
UMAP 2
UMAP 1
J
Integrated cluster
A
B
C
D
E
F
G
H
UMAP 2
UMAP 1
L
7 0919
5 0870
6 0918
4 0907
3 0891_B
2 0891_A
1 0892
2-21
2-19
2-18
2-17
2-16
2-15
2-14
2-13
AIT21 type
Integrated cluster
Saunders subcluster
I
AIT21 type
1 0892 NDB-SI-MA-STRv Lhx8 Gaba_4
2 0891 NDB-SI-MA-STRv Lhx8 Gaba_4_A
3 0891 NDB-SI-MA-STRv Lhx8 Gaba_4_B
4 0907 NDB-SI-MA-STRv Lhx8 Gaba_8
5 0870 NDB-SI-MA-STRv Lhx8 Gaba_2
6 0918 GPe-SI Sox6 Cyp26b1 Gaba_1
7 0919 GPe-SI Sox6 Cyp26b1 Gaba_2
NA (Saunders et al.)
UMAP 2
UMAP 1
K
Saunders subcluster
2-13
2-14
2-15
2-16
2-17
2-18
2-19
2-21
NA (present study)
UMAP 2
UMAP 1

### Supplemental Figure 3, related to Figure 3

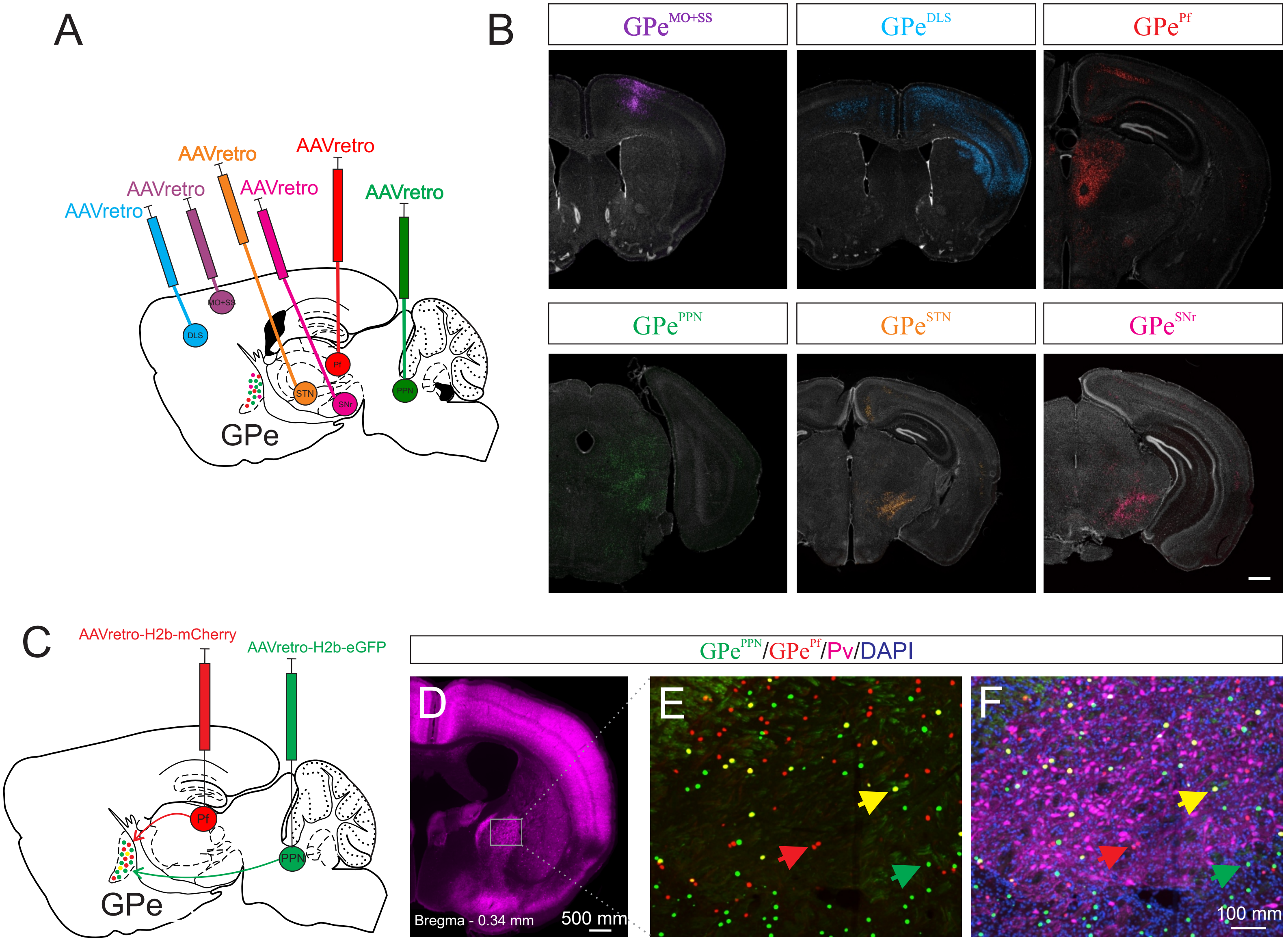

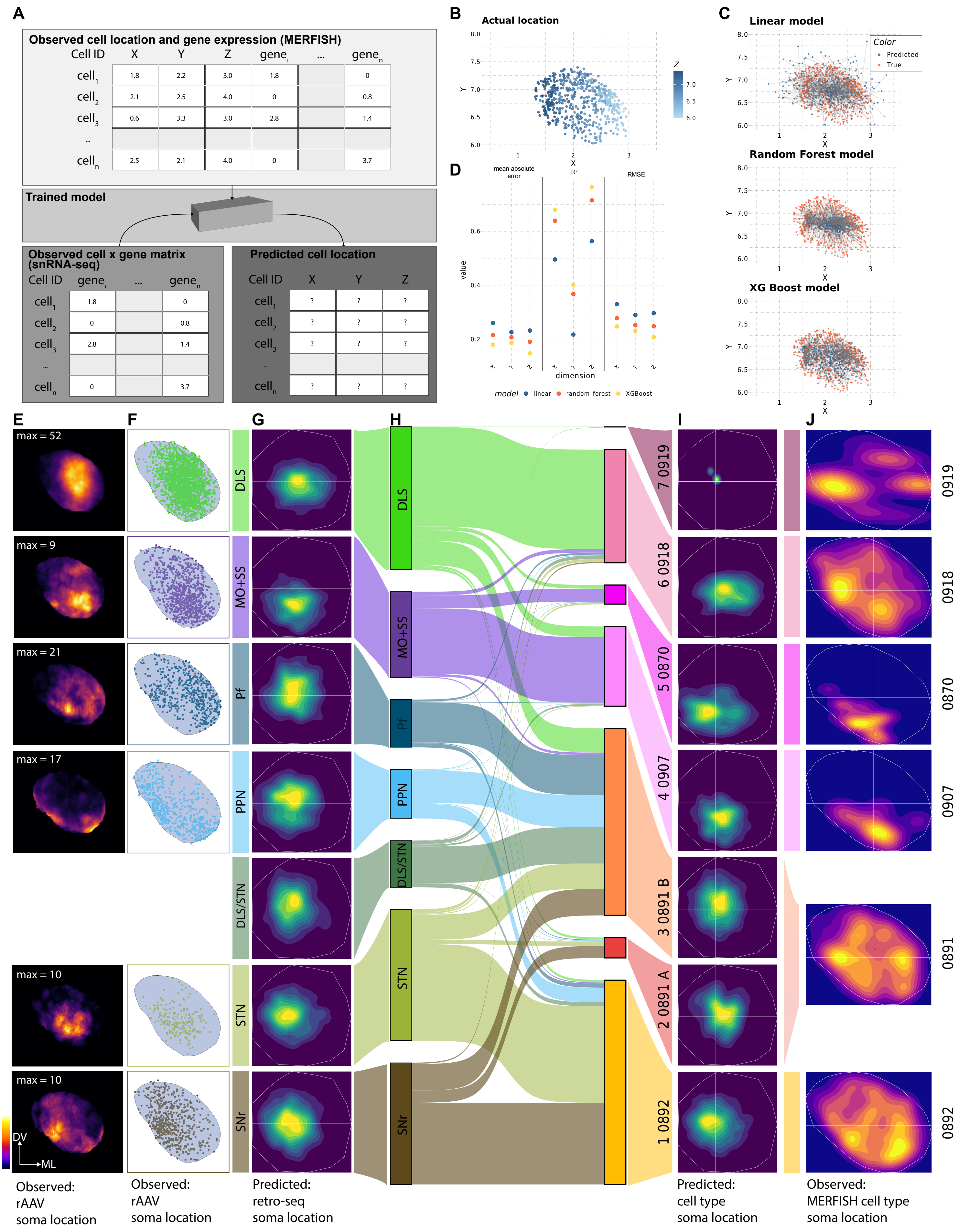
A
Observed cell location and gene expression (MERFISH)
Cell ID X Y Z gene ... gene
Trained model
Observed cell x gene matrix (snRNA-seq)
Predicted cell location
B
Actual location
C
Linear model
Random Forest model
XG Boost model
Color
Predicted
True
D
mean absolute error
R²
RMSE
value
dimension
model linear random_forest XGBoost
E
F
G
H
I
J
max = 52
max = 9
max = 21
max = 17
max = 10
max = 10
DLS
MO+SS
Pf
PPN
DLS/STN
STN
SNr
7 0919
6 0918
5 0870
4 0907
3 0891 B
2 0891 A
1 0892
0919
0918
0870
0907
0891
0892
DV
ML
Observed: rAAV soma location
Observed: rAAV soma location
Predicted: retro-seq soma location
Predicted: cell type soma location
Observed: MERFISH cell type soma location

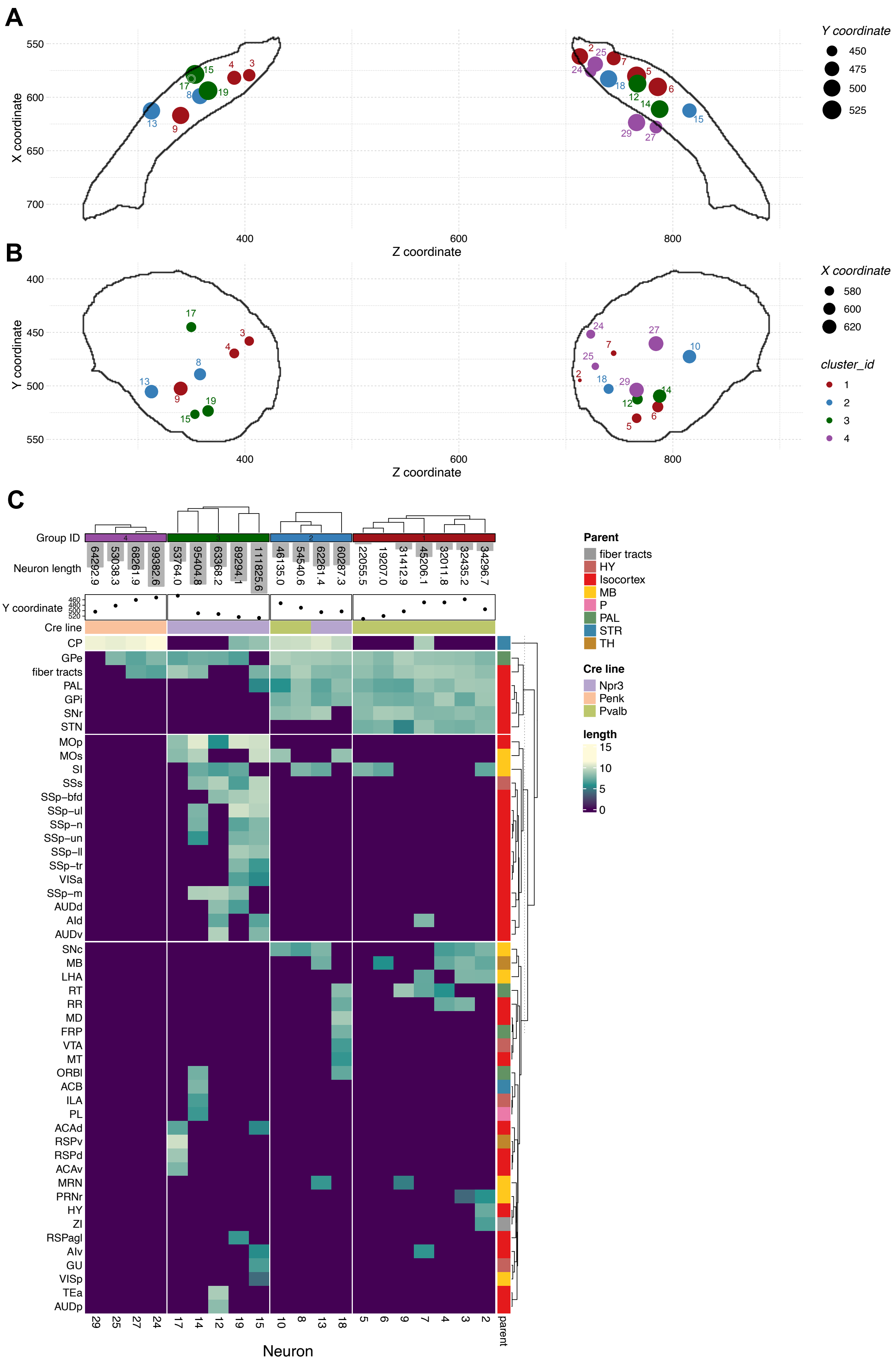
A
X coordinate
Z coordinate
Y coordinate
450
475
500
525
B
Y coordinate
Z coordinate
X coordinate
580
600
620
cluster_id
1
2
3
4
C
Group ID
Neuron length
64292.9
53038.3
68261.9
99382.6
53764.0
95404.8
63368.2
89294.1
111825.6
46135.0
54540.6
62261.4
60287.3
22055.5
19207.0
31412.9
45206.1
32011.8
32435.2
34296.7
Y coordinate
Cre line
CP
GPe
fiber tracts
PAL
GPi
SNr
STN
MOp
MOs
SI
SSs
SSp−bfd
SSp−ul
SSp−n
SSp−un
SSp−ll
SSp−tr
VISa
SSp−m
AUDd
AId
AUDv
SNc
MB
LHA
RT
RR
MD
FRP
VTA
MT
ORBl
ACB
ILA
PL
ACAd
RSPv
RSPd
ACAv
MRN
PRNr
HY
ZI
RSPagl
AIv
GU
VISp
TEa
AUDp
29 25 27 24 17 14 12 19 15 10 8 13 18 5 6 9 7 4 3 2
parent
Neuron
Parent
fiber tracts
HY
Isocortex
MB
P
PAL
STR
TH
Cre line
Npr3
Penk
Pvalb
length
15
10
5
0

Supplemental Figure 6, related to Figure 4

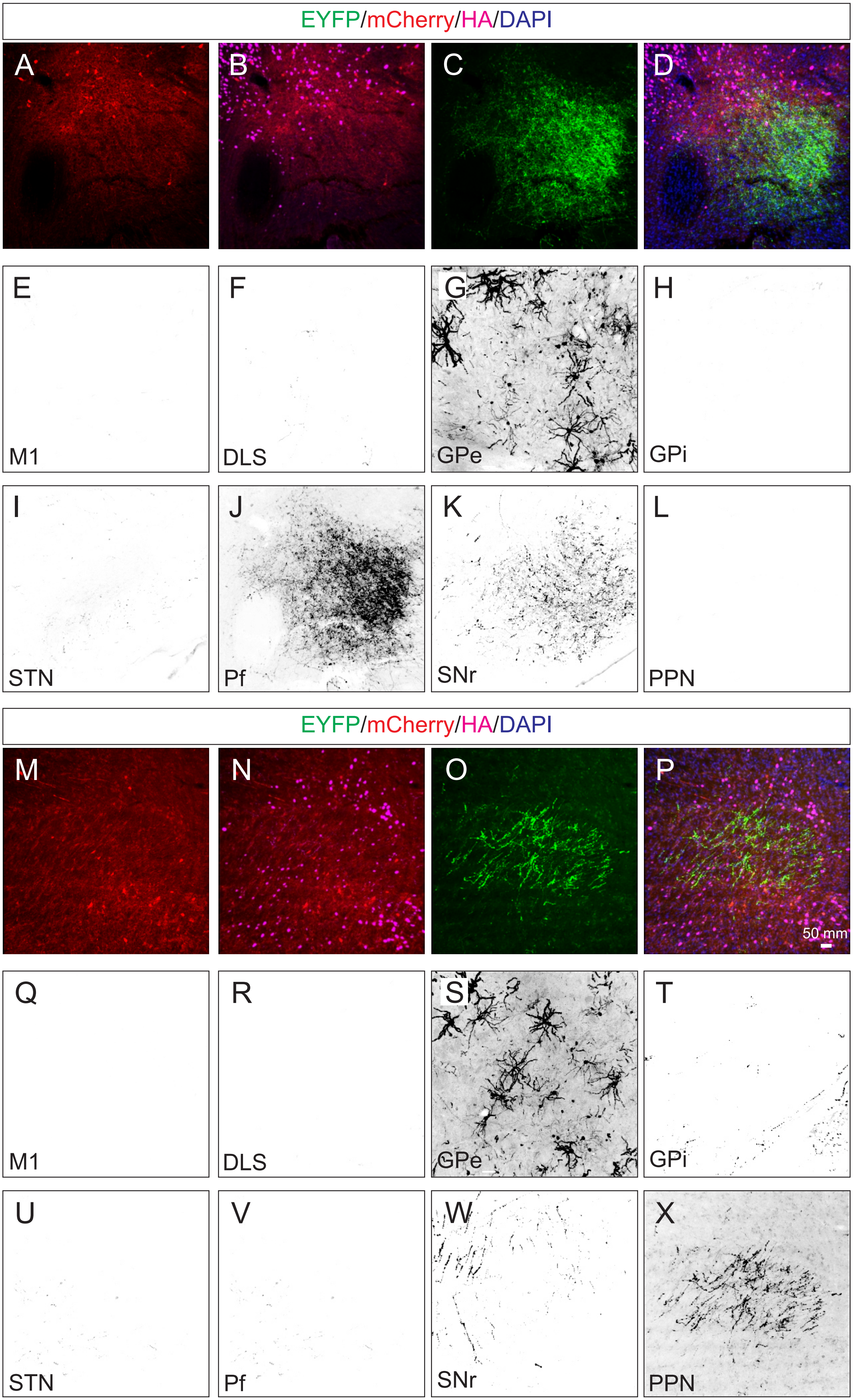

Supplemental Figure 7, related to Figure 6

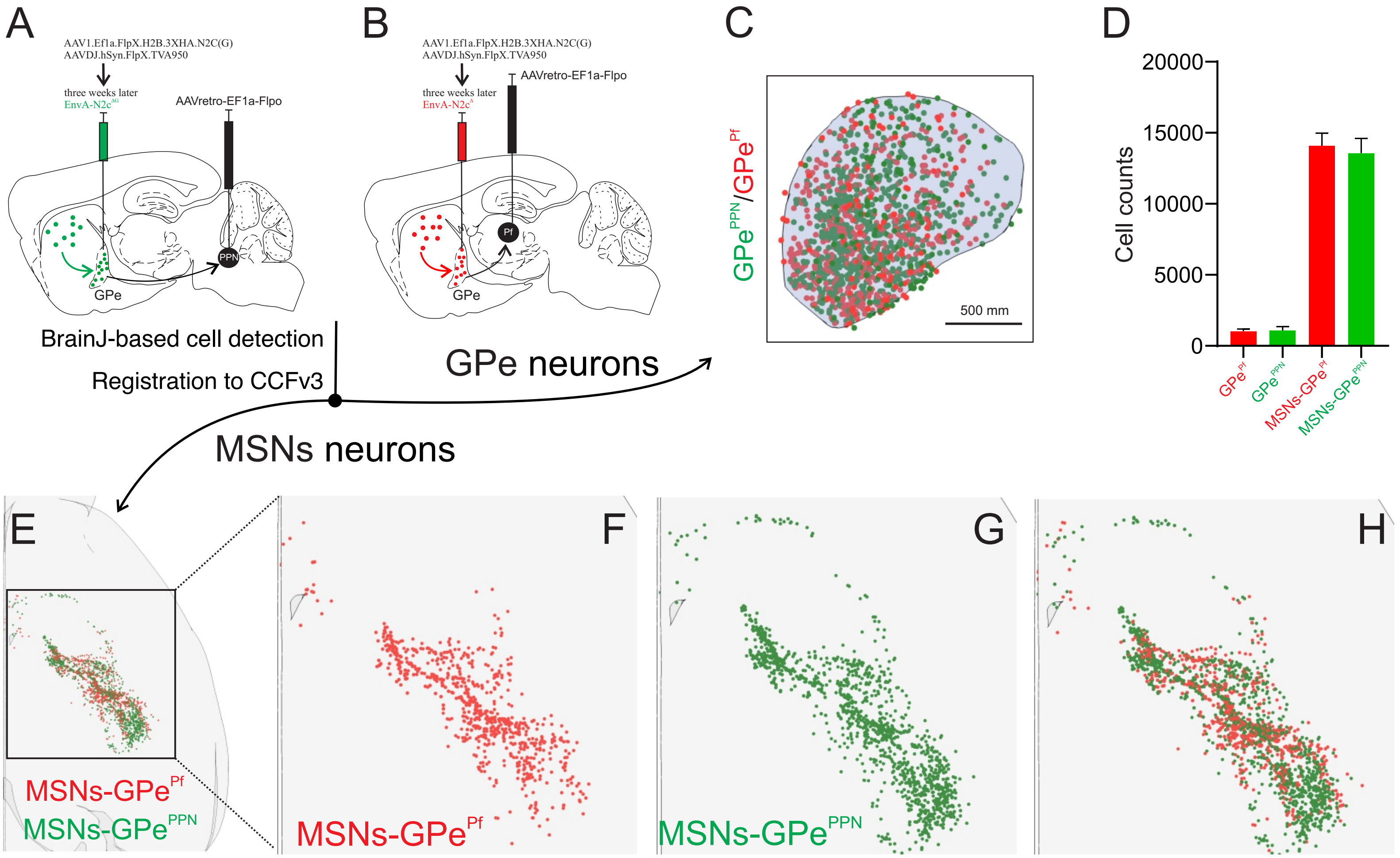

### Supplemental Figure 8, related to Figure 6

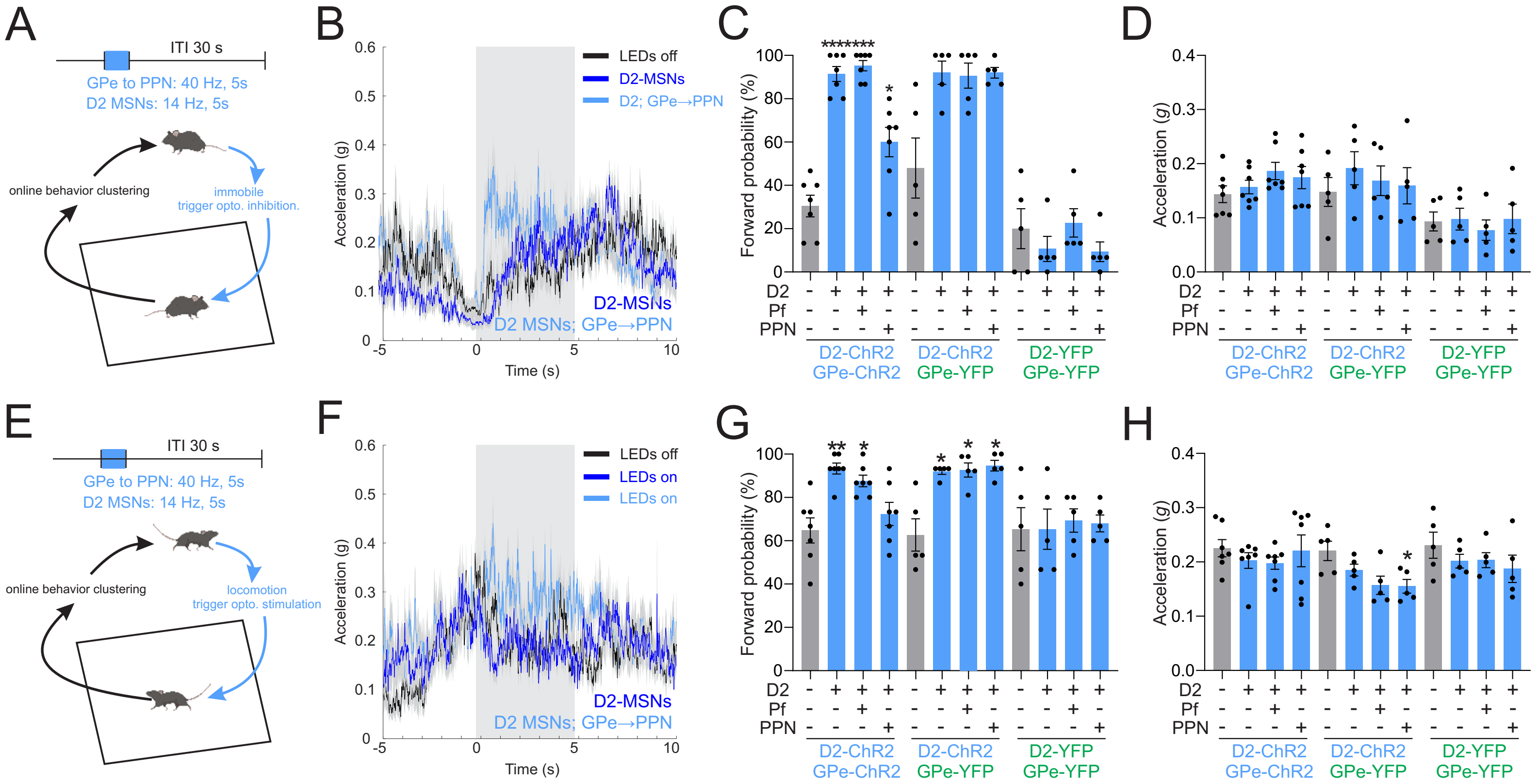
